# Whole-genome duplication reshaped adaptive evolution in a relict plant species, *Cyclocarya paliurus*

**DOI:** 10.1101/2022.09.04.506500

**Authors:** Yinquan Qu, Xulan Shang, Ziyan Zeng, Yanhao Yu, Guoliang Bian, Wenling Wang, Li Liu, Li Tian, Shengcheng Zhang, Qian Wang, Dejin Xie, Xuequn Chen, Zhenyang Liao, Yibin Wang, Jian Qin, Wanxia Yang, Caowen Sun, Xiangxiang Fu, Xingtan Zhang, Shengzuo Fang

## Abstract

*Cyclocarya paliurus* is a relict plant species that survived the last Glacial period and shows a population expansion recently. Its leaves have been traditionally used to treat obesity and diabetes with the well-known active ingredient cyclocaric acid B. Here, we present three *C. paliurus* genomes from two diploids with different flower morphs and one haplotype-resolved tetraploid assembly. Comparative genomic analysis revealed two rounds of recent whole-genome duplication events and identified 691 genes with dosage-effect that likely contribute to adaptive evolution through enhanced photosynthesis and increased accumulation of triterpenoids. Re-sequencing analysis of 45 accessions uncovered two bottlenecks, consistent with the known events of environmental changes, and many selectively swept genes involved in critical biological functions, including plant defense and secondary metabolism biosynthesis. We also proposed the biosynthesis pathway of cyclocaric acid B based on multi-omics data and identified key genes, in particular gibberellin related genes, associated with heterodichogamy in the species. Our research sheds light on evolutionary history and provides genomics resources to study the medicinal herb.

## Introduction

The “relict” plants consist of the remaining population of previously widely distributed species but restricted to limited geographic distribution currently. The severe population bottleneck is likely caused by large-scale environmental changes, such as global dramatic temperature decline, that have a fundamental impact on the ecosystem of the previously abundant species (Schweiger et al., 2008). Evidence reveals that the population dynamics are recorded in the population genomes (The 1001 Genomes Consortium, 2016). Recently developed sequencing technologies provide a practical approach to uncover the disaster during long-term evolutionary history (Li & Durbin, 2011; Liu & Fu, 2015). However, how relict plants survive previous environmental changes and flourish in the new territory remains unclear.

*Cyclocarya paliurus* (Batal.) Iljinskaja (wheel wingnut), the sole species in the genus *Cyclocarya* Iljinskaja (Juglandaceae), is not only a well-known multi-function tree species (Fang et al., 2006; Qin et al., 2021), but also has the character of heterodichogamy, a transitional form in the evolution of plants from monoecism to dioecism (Mao et al., 2019; Qu et al., 2021). Previous studies indicated that its leaves are often used in traditional Chinese medicine to treat hypertension and diabetes (Zhang et al., 2010; Zhou et al., 2021) due to its high biological activity and favorable safety. Furthermore, the antihyperglycemic tea of *C. paliurus* was the first health tea approved by Food and Drug Administration (FDA) in 1999 (Lin et al., 2020; Wang et al., 2020). These evidences show that *C. paliurus* leaves contain multiple bioactive compounds, including triterpenoids, flavonoids, phenolic acids, steroids, and polysaccharides, contribute to protecting humans against chronic diseases by antidiabetic, antioxidant, and antimicrobic properties (Mao et al., 2019; Sun et al., 2020; Xiao et al., 2020).

Triterpenoids are synthesized from six C5 (isopentenyl diphosphate) units from the common precursor 2,3-oxidosqualene by oxidosqualene cyclases (OSCs) (Hamberger et al., 2013). These carbon skeletons are further oxidized by cytochrome P450 monooxygenases (P450s) and glycosylated by UDP-dependent glycosyltransferases (UGTs), resulting in diverse triterpenoid structures (Seki et al., 2015; Miettinen et al., 2017). Triterpenoids constitute a vast family of natural products that play an important role in their significant biological and pharmacological effects, such as Cyclocaric acid B (CA-B) extracted from *C. paliurus* leaves (Zhu et al., 2015). CA-B has pharmacological activity on diabetes and it can enhance glucose uptake by involving AMP-activated protein kinase (AMPK) activation and improving insulin sensitivity in adipocytes (Zhu et al., 2015). However, the evolutionary history and functions of CA-B-related genes remain unknown, although ≥ 40 different triterpenoid compounds have been isolated from *C. paliurus* species (Wu et al., 2017; Jiang et al., 2019; Zheng et al., 2019). Therefore, the elucidation of biosynthetic pathways leading to the production of CA-B is greatly needed for heterologous bioproduction and a high public health priority.

With the importance in pharmaceutical values, a considerable production of *C. paliurus* leaves are required for production and medical use. However, *C. paliurus* seedlings can only be propagated from seeds but its seed quality is deficient with the seed vigor of 0-10% due to its heterodichogamy (Fu et al., 2011). Heterodichogamy in *C. paliurus* possess two temporally complementary morphs, protandry (PA) or protogyny (PG), in monoecious population. The stigma matures before pollen dispersal in PG whereas pollen scatters before stigma maturation in PA. Hence, the female and male function segregation within PA or PG significantly affects seed filling index and quality. Altougth heterodichogamy is especially common in the Fagales, Magnoliales, Malvales, Laurales, Sapindales, Canellales, Ranunculales, Zingiberales, Trochodendrales, Rosales, Caryophyllales, Malpighiales, and Apiales (Endress et al., 2020), the related genetic mechanism is far from well-studied.

The species *C. paliurus* is circumscribed with approximately two ploidy levels, including diploid (2n = 2x = 32) and auto-tetraploid (2n = 4x = 64), which mainly distribute across subtropical mountainous areas in China. Polyploidy or ancient whole-genome duplication (WGD) is a major driver of plant evolution (Guo et al., 2019) that contributes to variation in genome size, abundant genetic materials, phenotypic and functional diversification of plants (Jiao et al., 2014; Soltis et al., 2016). Although one polyploid *C. paliurus* genome was reported recently, the collapsed assembly missed haplotypic variations that may underlie important functions (Zheng et al., 2021), In addition, the exact roles that WGD played in origin and evolution of *C. paliurus* have not been clearly elucidated. Herein, we present three chromosome-scale genomes, containing two diploid *C. paliurus* that represent two different flower morphs (protogynous and protandrous), and one haplotype-resolved genome for auto-tetraploid. The comprehensive genomic resources of this species allows us to uncover the genetic mechanisms behind the special characteristics of *C. paliurus* species, including heterodichogamy, the origination and polyploidization, and the pathway of CA-B biosynthesis.

## Results

### Genome assemblies and annotation of the three *C. paliurus* genomes

We sequenced and assembled three *C. paliurus* genomes, including two diploid and one tetraploid accessions (**Fig. 1 and Table 1**). The two diploid *C. paliurus* represent two different reproduction types in hermaphroditism, with one being protandrous (PA-dip) and another protogynous (PG-dip), while the tetraploid is protandrous (PA-tetra). Karyotype analysis identified 32 chromosomes in the diploid genomes and 64 in tetraploid (**Supplementary Figure 1**). Taking the male florals of PA-tetra *C. paliurus* as experimental materials, the homologous chromosomes synapsis at the early stage of meiosis in pollen mother cells of PA-tetra *C. paliurus* were studied. Multivalents (quadrivalent) phenomena were observed in pollen mother cells (**Supplementary Figure 2**), strongly indicating that the PA-tetra is an auto-tetraploid. Using diploid *Pterocarya stenoptera* (600 Mb) as an internal reference, we estimated the genome size of PA-dip, PG-dip, and PA-tetra *C. paliurus* by flow cytometry (FCM) to be about 606 Mb (1C), 659 Mb (1C), and 2,460 Mb (1C = 1,230 Mb), respectively (**Supplementary Figure 3**). A total of 134.9, 75.5 and 271.8 Gb subreads were generated on the PacBio Sequel II platform, comprising ∼223×, 115× and 221× coverage of the estimated genome sizes (by FCM, 1C) for PA-dip, PG-dip and PA-tetra, respectively (**Supplementary Table 1**). The initial contigs were assembled using CANU assembler, resulting in three contig-level assemblies with N50s of 1.9 Mb in PA-dip, 1.4 Mb in PG-dip and 431 Kb in PA-tetra (**Supplementary Table 2**). The assembled genome sizes were 586.62 Mb for PA-dip and 583.45 Mb for PG-dip, accounting for 97.8% and 97.2% of estimated genome size by flow cytometry (**Supplementary Figure 3**), respectively. 2.38 Gb sequences were assembled in PA-tetra, almost four times of the haploid size (**Supplementary Table 2**). The chromosomal level assemblies were achieved using High-throughput Chromatin Conformation Capture (Hi-C) technology (**Supplementary Table 3**). For the diploid PA and PG genomes, 543.53 Mb (92.65%) and 553.87 Mb (94.93%) of sequences were integrated into 16 pseudo-chromosomes, respectively **(Fig. 1g)**. The PA-tetra genome comprised 64 pseudo-chromosomes with four sets of monoploid chromosomes using ALLHiC phasing algorithm that anchored 2.17 Gb (91.08%) of genomic sequences, representing a haplotype-resolved assembly of the tetraploid species **(Fig. 1g, Supplementary Figure 4, and Supplementary Table 4)**. We identified 12,688,530 haplotypic variations among the haplotype-resolved A/B/C/D homologous chromosomes, affecting 5,152 functional genes (**Supplementary Table 5)**. These genes were significantly enriched in some primary biological pathways, such as aminoacyl-tRNA and fatty acid biosynthesis, glycerolipid metabolism, mitochondrial genome maintenance, and single strand break repair **(Supplementary Figure 5)**. We further assessed the quality of genome assemblies, showing more than 95.2% of BUSCO completeness (**Supplementary Table 6**). Comparison with the previously published *C. paliurus* genome shows an improved BUSCO completeness (95.2% v.s. 91%) with well-resolved duplicated genes (i.e., allelic genes; BUSCO duplication 82.3% v.s. 8.4% in **Supplementary Table 6**). (Zheng et al., 2021). Our assessment using Illumina short reads showed at least 98.89% of global mapping ratio and 93.72% of properly paired reads **(Supplementary Table 7)**. Comparison among the three genomes revealed high levels of syntenic relationship with a large number of genes (26,760) located in the syntenic regions (**Fig. 1g**). The Hi-C contact heatmaps also confirmed the high consistency of genome structure and quality, which also shows an improved chromosomal-scale assembly in the comparison with the previously published genome (Zheng et al., 2021) **(Supplementary Figures 6-8)**.

**Table 1.**
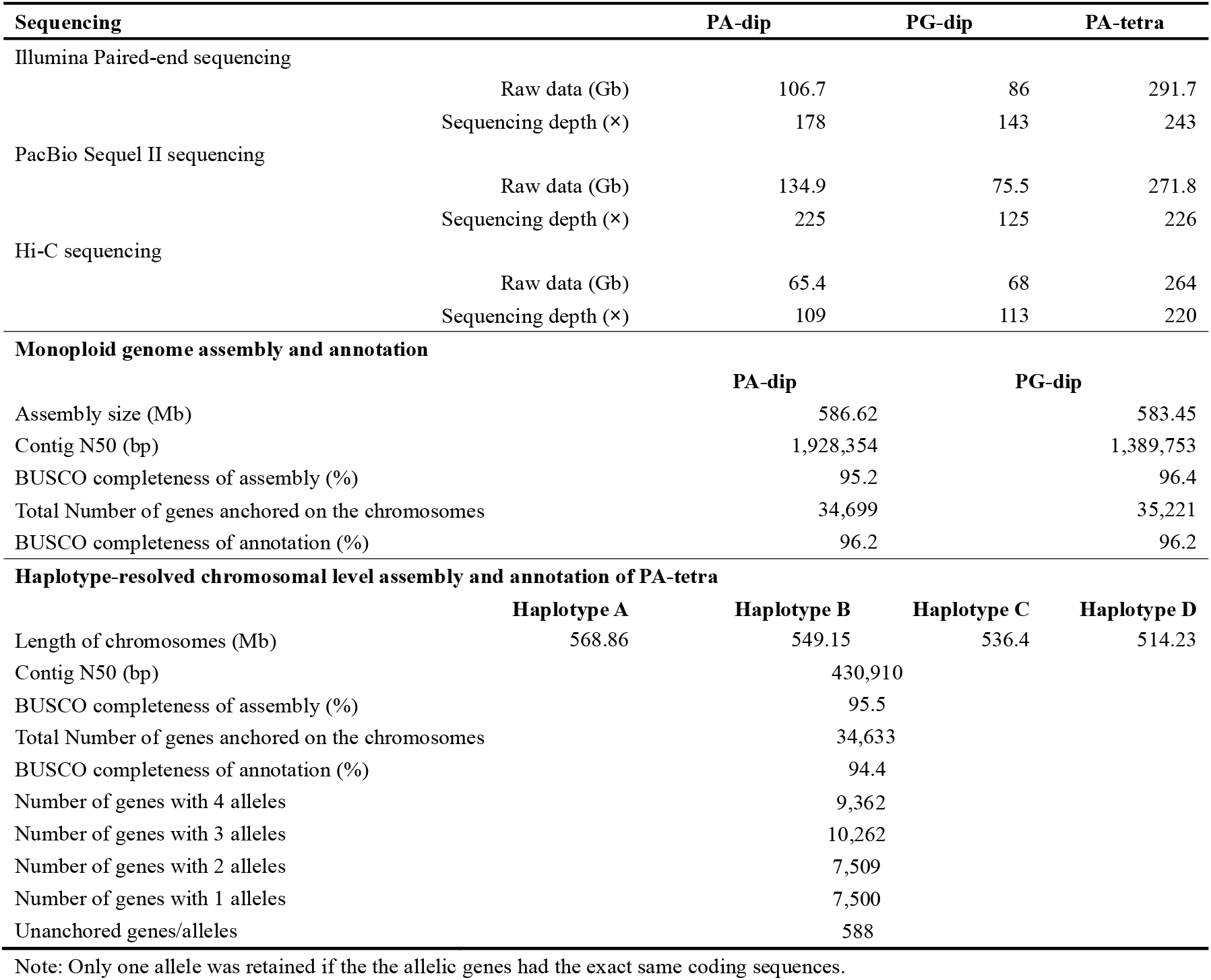
Summary on genome assembly and annotation of *C. paliurus*

**Fig. 1.**
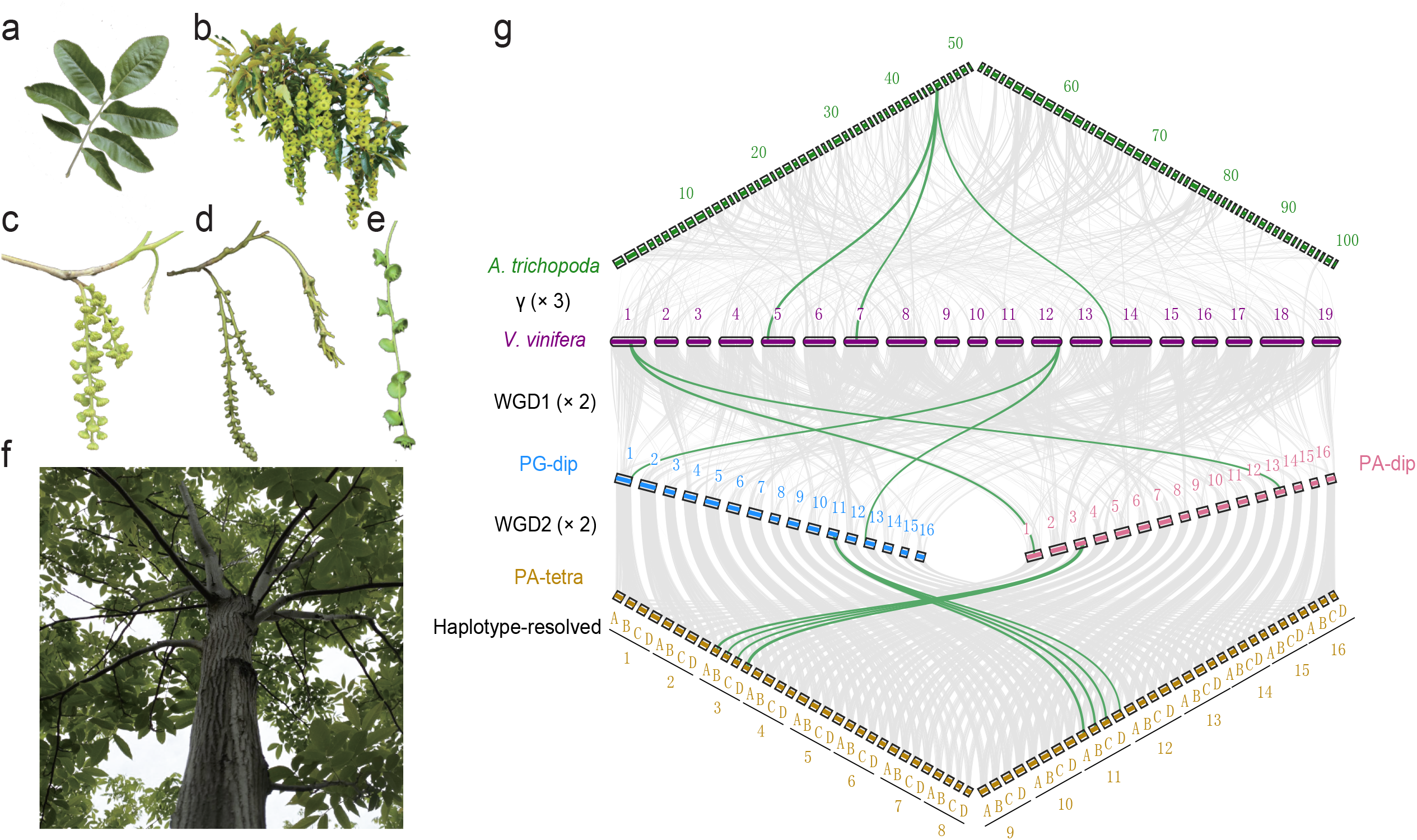
Morphology and genome duplications of *C. paliurus*. (a) Imparipinnate leaves. (b) Mature fruits. (c) Female and male flowers of PA-dip. (d) Female and male flowers of PG-dip. (e) Fruit wings that start to spread. (f) Tetraploid plant. (g) Genomic alignments between the basal angiosperm *Amborella trichopoda*, and the basal eudicot *Vitis vinifera*, as well as PG-dip, PA-dip, and PA-tetra *C. paliurus* are shown. The conserved collinear blocks are shown as the gray lines in the background and the green lines indicate cases in each round of whole genome duplication.

We annotated 34,699 protein-coding genes in PA-dip and 35,221 in PG-dip. The initial annotation of the tetraploid genome resulted in 90,752 gene models; however, this number mixed the concept of genes and allelic genes. To identify allelic genes that have at least one base substitution, we adopt the same strategy in our previously published sugarcane genome (Zhang et al., 2018), leading to 34,633 allele-defined protein-coding genes in the tetraploid genome (**Supplementary Table 8-9**). Our assessment of the annotation showed 96.2%, 96.2% and 94.4% of completeness for PA-dip, PG-dip, and PA-tetra, respectively, according to the 1,375 conserved genes in BUSCO assessed using embryophyta_odb10 database **(Supplementary Table 10)**.

A total of 282.25 Mb (48.1% of the assembled genome), 316.95 Mb (54.3%) and 1,154.35 Mb (48.4%) repetitive sequences were identified in the PA-dip, PG-dip and PA-tetra genomes, respectively, showing a slight increase in PG-dip genome **(Supplementary Table 11)**. The ratio of repetitive elements in the previously reported assembly by Zheng (14.94%) is much lower than our results, possibly due to a large proportion of collapsed sequences. Retroelements account for approximately three-quarters of the repetitive sequences, ranging from 35.7 to 37.3% in the three genomes. However, in contrast to other published plant genomes, such as pineapple (Chen et al., 2019), sugarcane (Zhang et al., 2018) and banyan tree (Zhang et al., 2020), LINE is the most prominent family in *C. paliurus*, spanning from 12.16% to 12.59% of the assembled genomes. In comparison, *Copia* and *Gypsy* account for only ∼5.47% and ∼5.94% of the assembled genomes on average in *C. paliurus* genomes **(Supplementary Table 11)**.

### Phylogeny and polyploidy evolution

Maximum likelihood tree using 302 single-copy gene families from nine plant species reconstructed the phylogenetic relationship among *C. paliurus* and related species. The estimated divergence time between *C. paliurus* and *Pterocarya stenoptera* was approximately 46.07 million years (Mya) ago **(Fig. 2a)**, consistent with the previously reported divergence time of different genera of Juglandaceae (Sun, 2005). Analysis of the gene families showed that a large number of families experienced expansion (285) and contraction (264) compared to *P. stenoptera* **(Fig. 2a)**. In addition, we found 1,738 gene families specific to the *C. paliurus* genome, while 9,917 gene families were shared in the selected species, demonstrating evolutionary conservation **(Fig. 2b)**.

**Fig. 2.**
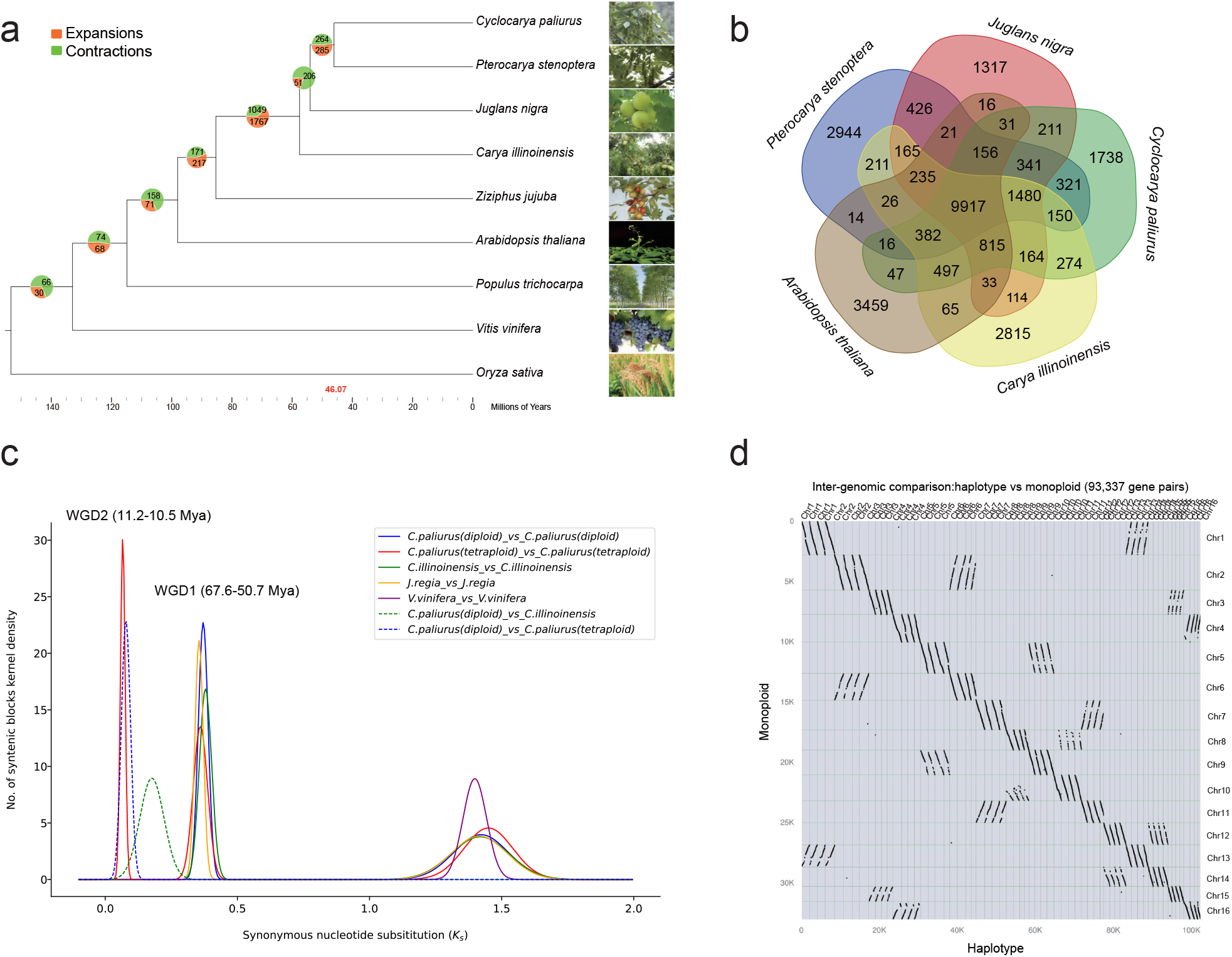
Phylogenetic and Comparative Analysis of *C. paliurus*. (a) Phylogenetic relationship of *C. paliurus, C. illinoinensis, J. nigra, P. stenoptera, A. thaliana, Ziziphus jujuba, Vitis vinifera, Populus trichocarpa*, and *Oryza sativa*. The divergence times among different plant species are labeled in the bottom. (b) Venn diagram of orthologous and species-specific genes in different plant genomes. (c) Evolutionary analysis of the diploid and tetraploid *C. paliurus* genomes. The distribution of synonymous substitution rates (*Ks*) of orthologs. (d) Synteny analysis between PA-tetra and PA-diploid genomes. The full name ‘monoploid’ indicates a reference genome assembly with only one representative haplotype retained, while the ‘haplotype’ indicates fully phased genome with all the four haplotypes.

The distribution of synonymous substitution ratios (Ks) of each homologous gene pair within *C. paliurus* showed three peaks **(Fig. 2c)**, representing three whole-genome duplication events. In addition to the ancient whole-genome triplication (WGT) event shared with grape, *C. paliurus* experienced two recent WGDs. Synteny analysis between PA-dip and PG-dip validated the early WGD (i.e., WGD1), dating back to ∼67.6-50.7 Mya (**Supplementary Figure 9 and Fig. 2c**). Most of the duplicated chromosomes maintained high levels of syntenic relationship and completeness compared to their counterparts. However, several structural variations were observed (**Supplementary Figure 9**), for instance, an inversion between Chr4 and Chr16, and a large deletion in Chr8 compared with Chr10, indicating the diploidization process in the diploid *C. paliurus* after the early WGD. The most recent WGD event (i.e., WGD2) happened in ∼11.2-10.5 Mya and contributed to the tetraploidy in *C. paliurus* species (**Fig. 2c-d**). To check whether each of two WGDs made specific contribution to specific gene family expansion in *C. paliurus*. We counted 1,351 and 2,024 genes, which experienced WGD1 and WGD2 events, respectively. Functional enrichment analysis showed that genes involved in WGD1 event were significantly enriched in ribosomal subunit assembly, mannosyltransferase activity, N-Glycan biosynthesis, and the genes involved in WGD2 event were mostly enriched in terpene biosynthesis, such as sesquiterpenoid, triterpenoid, and monoterpenoid biosynthesis (**Supplementary Figure 10**). Meanwhile, the genomic dot plots between *C. paliurus* and *Vitis vinifera* validated that tetraploid *C. paliurus* experienced two WGD events during the polyploidy evolution (**Supplementary Figures 11-12**). The fully phased haplotypes facilitate us to investigate the evolution of polyploidy in *C. paliurus*. According to the method described by Mitros (2020) that depend on clustering of chromosome-specific *K*-mers (*K*=13), we found that four haplotypes within each homologous group were consistently partitioned into a same branch (**Supplementary Figure 13**), indicating a similar evolutionary history of the four haplotypes. In addition, the smudge pot analysis using heterozygous *K-*mer pairs extracted from Illumina sequencing reads suggested a highly heterozygous tetraploidy evidenced by the dominant component of AAAB pattern, accounting for 57% of tested *K*-mer pairs (**Supplementary Figure 4d**). These evidences collectively suggest that PA-tetra is likely an auto-tetraploid species with a high heterozygosity of 1.97% (**Supplementary Figure 4c**).

### Expansion of P450 gene families associated with elevated triterpenoids biosynthesis

Evidences showed that triterpenoid, sterols, flavones, and phenols acids were enriched in *C. paliurus* leaves (Fu et al., 2009; Yang et al., 2018). It is possibly associated with expansion of specific gene families related to biosynthesis of these secondary metabolism. GO and KEGG functional enrichment analysis showed that many of the 2,234 expanded genes (**Fig. 2b**) were enriched in sesquiterpene, terpene, monoterpenoid, and triterpenoid biosynthesis pathways (**Supplementary Figures 14-15**). For instance, we found 15% (42/280 in total) of cytochrome P450 monooxygenases (P450s) expanded, among which 23 genes were clustered on Chr1, Chr4 and Chr12 (**Supplementary Figure 16**). Phylogenetic analysis further identified that, among the 42 expanded P450s, 16 genes belong to CYP89 family that participates in biosynthesis of tetracyclic triterpenoids (Zhou et al., 2016), 18 genes belong to CYP706 family, and six genes belong to CYP82 family, possibly contributing to the increased flavonoids (Awasthi et al., 2016) (**Supplementary Table 12 and Supplementary Figure 17**).

### Genes associated with heterodichogamy in *C. paliurus*

The most typical feature of *C. paliurus* is heterodichogamy with female and male functionally separated within protandry (PA) or protogyny (PG) individuals, promoting outbreeding in an independent population. To investigate the genes triggering heterodichogamy in *C. paliurus*, we performed a comparative floral bud transcriptome analysis of PG and PA samples, respectively. These samples contain two tissue types (female and male floral buds) at five different developmental stages, namely from S0 to S4 (Chen et al., 2019).

Pairwise comparison between PG and PA female samples (PG-F v.s. PA-F) identified 958 differentially expressed genes (DEGs) that were consistently upregulated or downregulated in at least two stages. Similarly, 2,373 DEGs were found between PA and PG male samples (PA-M v.s. PG-M; **Supplementary Figure 18a-b**). Functional analysis revealed that these DEGs were enriched in a series of biological processes involving in floral organ formation and development (**Supplementary Figure 18c-d and 19-20**). Notably, many of up-regulated DEGs (128/855 in PG-F and 58/1539 in PA-M) were related to hormone biosynthesis and signaling pathways, indicating that hormone may play an important role in contributing to the heterodichogamy morphs with asynchronous flowering. We further tested endogenous hormone contents in PA and PG floral buds, including gibberellin (GA_3_), auxin (IAA), and abscisic acid (ABA). The results displayed similar levels of IAA and ABA at each of the five stages especially at S0, an initial developmental stage of floral buds (**Supplementary Figure 21**). However, significantly increased levels of GA_3_ in PG samples were detected at S0 stage compared to PA (**Supplementary Figure 21**), indicating that GA_3_ content likely plays a crucial role in regulating floral buds physiological differentiation and was responsible for the asynchronous flowering.

To investigate the co-expression networks during floral buds development, we identified co-expressed gene sets via WGCNA package based on the 22 RNA-seq data (**Supplementary Figure 22**). After filtering genes with low expression, a total of 20,829 genes were retained, distributing in 26 modules (**Supplementary Figure 22-23**). We observed that genes in three modules (darkorange: 47 genes; pink: 912 genes; red: 1,244 genes) showed a high correlation (R^2^ ≥ 0.56, *P* ≤ 0.007, Pearson test) with GA_3_ content (**Supplementary Figure 24-25**). In addition to ‘regulation of hormone and gibberellin biosynthetic process’, these genes were also functionally enriched in signal transduction, response to stimulus and stress, and biological regulation (**Supplementary Figure 26-27**).

Key hub genes including transcription factors (TFs) were identified in the WGCNA analysis. For instance, *Trihelix-1* (*CpaF1st06806*), involving in endogenous hormone signaling and flower development (Nagata et al., 2010), and *ERF066* (*CpaF1st15865*), responsible for embryos development and stress signal transduction (Rashotte et al., 2006), have the most edges (342 and 279, respectively; **Supplementary Figure 24e**) in pink module network. In red module network, the top two most frequently connected hub genes, *ERF090* (*CpaF1st01445*) and *WRKY55* (*CpaF1st00113*) (**Supplementary Figure 24f**), were identified that may play important functions in regulating the development of floral organs (Chuck et al., 1998) and phytohormone-mediated signal transduction process (Zhang et al., 2004).

### Dosage-effect contributed to enhancing photosynthesis and increasing accumulation of terpenoid

Investigation of the seedlings’ morphology, anatomical structure, and photosynthetic capacity of leaves showed enhanced photosynthesis and acceleration of plant growth in the tetraploid *C. paliurus*. We observed that a series of growth indices in the tetraploid plants were significantly higher than that in diploid individuals (*p* < 0.01, Duncan-test), including seedling size, leaf area, length of compound leaf (**Supplementary Figure 28 and Supplementary Table 13**), the thickness of upper and lower epidermal cells, palisade mesophyll, sponge tissue, and blade, stomatal size, stomatal density (**Fig. 3a and Supplementary Figures 29-30**), net photosynthetic rate, and chlorophyll content (**Fig. 3b-c**). In addition, we also detected three growth indices including blade aspect ratio, leaf moisture content, and leaf specific weight, which were significantly lower in the tetraploid than that in diploid individuals.

**Fig. 3.**
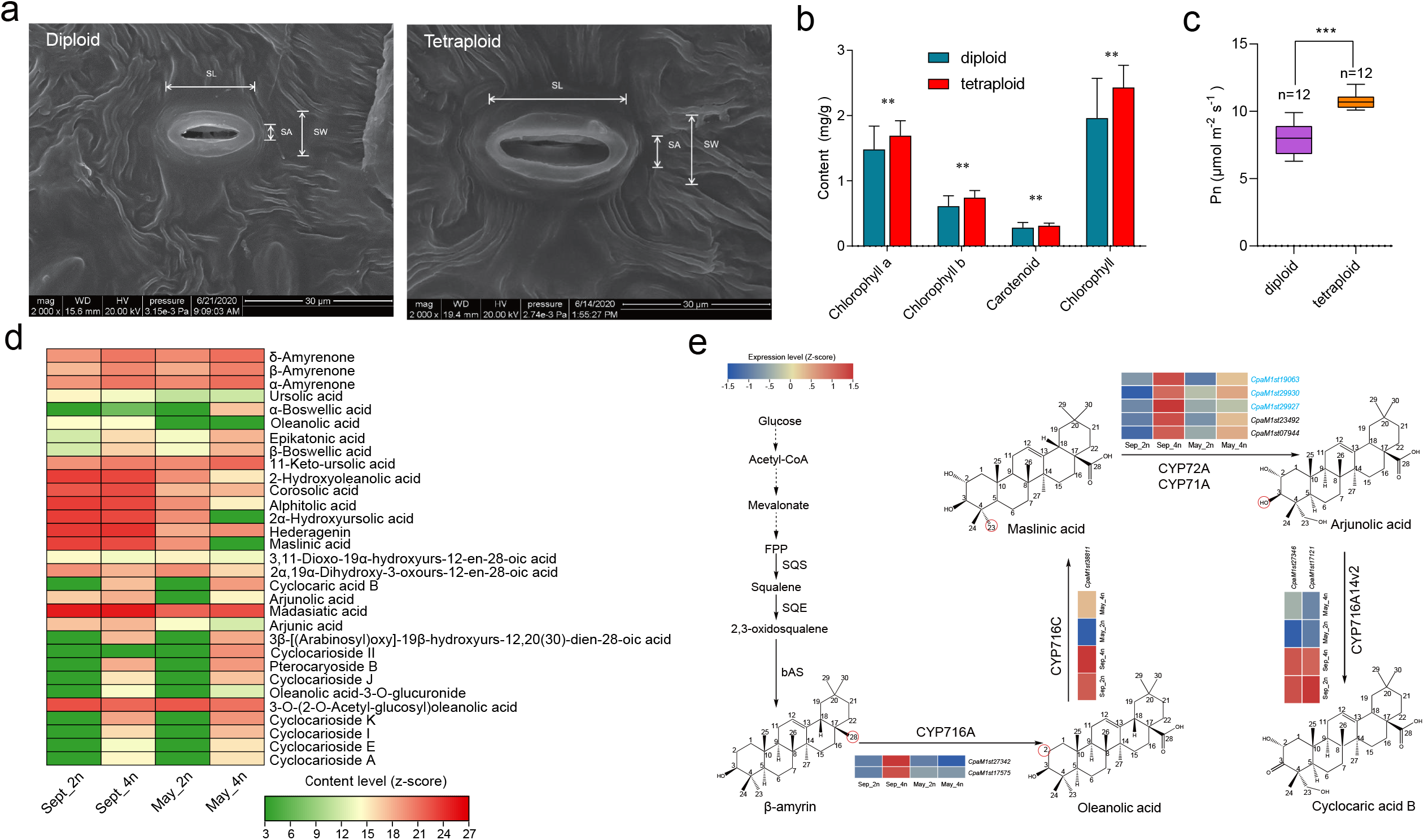
Dosage-effect contributes to increased growth adaptability and accumulation of terpenoids. (a) Scanning electron microscopy (SEM) of stomata in diploid and tetraploid *C. paliurus* leaves. (b) The comparison of chlorophyll content between diploid and tetraploid *C. paliurus*. Statistical significance (n = 5) was determined using the two-sided Student’s t test. Error bars indicate mean±SD of indicated replicates. **, P value < 0.01. (c) The comparison of net photosynthetic rate (Pn) between diploid and tetraploid *C. paliurus*. ***, P value < 0.001.(d) Heatmap showing the accumulation patterns of triterpenoids among the four samples (Sept_2n: diploid samples collected in September; Sept_4n: tetraploid samples collected in September; May_2n: diploid samples collected in May; May_4n: tetraploid samples collected in May). (e) Expression profiles of genes associated with Cyclocaric acid B (CA-B) synthesis in *C. paliurus* tender and mature leaves for different ploidy. The scale ranging from blue (low) to red (high) indicates the expression magnitude of the FPKM values. The genes of CYP72A subfamily are shown in blue. The functions of CYP716C and CYP716A14v2 were identified in Karel (2017) and Moses’s (2015) report.

To investigate genes underlying the functional impact of polyploidization, we identified 691 genes that showed significantly elevated expression in the tetraploid samples compared to diploid samples (Fold Change ≥ 2 and P-value ≤ 0.05; **Supplementary Figure 31**), which were considered as dosage-effect genes. Furthermore, functional annotation showed that these genes were abundantly enriched in some primary biological pathways (**Supplementary Figure 32**). Notably, genes, which are involved in carbohydrate, starch, sucrose, alanine, aspartate, and glutamate metabolism, phosphatidylinositol signaling system, and ion channels, play vital roles in photosynthesis, and are indispensable for plant growth, development, and stress responses (Jone & Varela, 1998; Boriboonkaset et al., 2013; Tegeder et al., 2014). We also explicitly investigated the *A. thaliana* key homologous proteins in the photosynthetic pathway. Carbonic anhydrase (CA), phosphoenolpyruvate carboxylase kinase (PPCK), ribulose-1,5-bisphosphate carboxylase/oxygenase (Rubisco), fructose-1,6-bisphosphatase (FBP), sedoheptulose-1,7-bisphosphatase (SBPASE) have significantly higher expression in tetraploid than that in diploid (*P* ≤ 0.05; **Supplementary Figure 33**). Meanwhile, we identified 759 dosage compensation effect genes with higher expression in diploid than in tetraploid (Fold Change ≥ 2 and P-value ≤ 0.05). GO and KEGG functional enrichment analysis showed that many of the dosage compensation effect genes were enriched in regulation of DNA recombination, maltose metabolic process, cysteine and methionine metabolism, amino sugar, and nucleotide sugar metabolism pathways (**Supplementary Figure 34**).

Interestingly, we also noticed that many of these genes were significantly enriched in sesquiterpenoid and triterpenoid biosynthesis, terpenoids metabolism (P value<0.05; **Supplementary Figure 32**), and likely contributed to the increased accumulation of some triterpenoid components in the PA-tetra (**Fig. 3d**). Furthermore, we observed that 22 dosage-effect genes belong to the Cytochrome P450 monooxygenases (P450s) family based on BLAST results in public databases and phylogenetic analysis (**Supplementary Figures 35-36 and Supplementary Note**). Among them, three P450 subfamilies (CYP716A, CYP72A and CYP71A) might be vitally important in the biosynthesis of Cyclocaric acid B (CA-B; **Fig. 3e**) via modification of different C positions. Previous researches showed that CYP716A12 and CYP716A1 can catalyze the oxidation at the C-28 position of β-amyrin, forming the triterpene oleanolic acid (Fukushima et al., 2011; Yasumoto et al., 2016). While, the maslinic acid might be hydroxylated by CYP71A16 and CYP72A397 specifically at C-23 position into triterpene arjunolic acid (Castillo et al., 2013; Han et al., 2018). Our study identified two homologous genes in *CYP716A*, two in *CYP71A*, and three in *CYP72A* showing dosage-effect in the tetraploid (**Fig. 3e**) and they were likely the key genes contributing to biosynthesis pathway of CA-B (specific triterpene to *C. paliurus*).

### Population structure and evolutionary history

To explore the population structure and evolutionary history of the *C. paliurus* population, we re-sequenced 45 accessions, including ten diploid and 35 tetraploid individuals native to the south of China, and one walnut species (*Juglans regia*) as an outgroup (**Fig. 4a and Supplementary Table 14**). A total of 3.89 and 26.67 million variants were respectively identified from diploid and tetraploid populations based on our stringent filtering criteria (**Methods**), including 3.55 and 23.08 million SNPs, and 0.34 and 3.60 million indels. We also identified that the two populations contained 3,845 and 38,899 variants in genic regions, including 3,753 and 23,764 synonymous, 5,503 and 35,241 nonsynonymous, respectively (**Supplementary Table 15**).

**Fig. 4.**
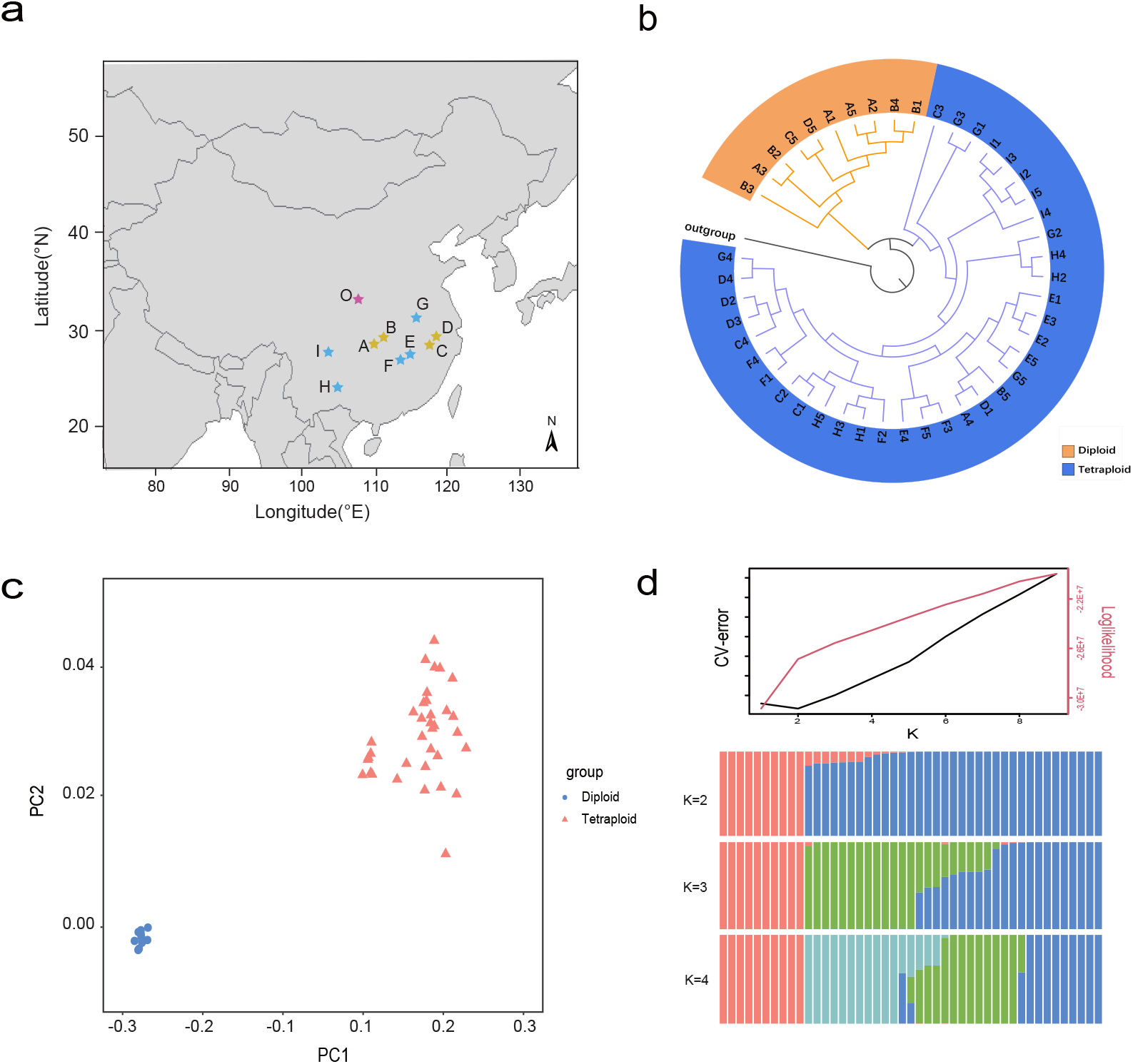
Phylogenetic splits among *C. paliurus* populations. (a) Dispersion of 45 individuals sampled from 9 sites across most of the geographic range of *C. paliurus*. Populations are plotted with dots color-coded based on dispersion by latitude and longitude. Yellow color stars represent the co-presence of diploid and auto-tetraploid distribution, blue color stars represent only auto-tetraploid distribution, and purple color star represents the outgroup (*J. regia*) (more details were shown in Supplementary Table 14). The world map was constructed from the Natural Earth dataset (http://www.naturalearthdata.com). (b) A phylogeny for *C. paliurus* accessions, estimated from SNPs in neutrally evolving sites. (c) PCA shows clear separation between diploid and auto-tetraploid populations. (d) ADMIXTURE plot for *C. paliurus* showing the distribution of K=L2, 3, 4, and 5 genetic clusters, among them, K=L2 (representing the divergence within diploid and auto-tetraploid clades) genetic clusters indicates the smallest cross-validation error. Cross-validation plots display CV-error versus K suggests K = 2 is also the best fit.

A phylogeny of these *C. paliurus* individuals collected from nine geographic locations partitioned these samples into two distinct groups (**Fig. 4b**). The ten diploid samples were clustered in the first group and most closely related to the outgroup *Juglans regia*. The remaining 35 tetraploid samples were represented in the second group. Both principal component analysis (PCA) and ADMIXTURE analyses supported the population structure (**Fig. 4c-d**). These results indicated a single origin of the last WGD event in *C. paliurus* species rather than multiple origins observed in other polyploid species, such as sugarcane (Zhang et al., 2018).

To identify the candidate genes responsible for coordinated local adaptation, we further analyzed selective sweeps based on the SweeD analysis in both diploid and auto-tetraploid genomes. A total of 25.88 Mb and 165.18 Mb genomic sequences were under purifying selection in diploid and tetraploid genomes, respectively (**Supplementary Figure 37**). These selectively swept regions were evenly distributed in 16 chromosomes of PA-dip and 64 chromosomes of PA-tetra *C. paliurus*, with a number of them showing high levels of selective sweeps (**Supplementary Figure 37**). These swept genomic regions overlapped with 1,529 protein-coding genes in diploid, and 4,812 allele-defined genes in the tetraploid group. Global GO and KEGG functional enrichment analysis of the swept genes also revealed that the number of 114 and 568 genes respectively in diploid and tetraploid groups were involved in important functions, such as secondary metabolism, regulation of DNA recombination, and maltose metabolic process (**Supplementary Figures 38-41**).

Among these swept genes, 262 were shared between the diploid and tetraploid genomes, and also located in the syntenic regions. In addition, the selectively swept genes specific to tetraploid were mostly enriched in terpene synthase activity and terpene biosynthesis (**Supplementary Figures 42-43**), probably caused by stronger environmental adaptation (Jiang et al., 2019) and stress tolerance (Boncan et al., 2020) than diploid. Further, three genes were verified as orthologs of the *A. thaliana* genes *FAR1-RELATED SEQUENCE* (*FRS10, CpaM1st41668*), *ARABIDOPSIS RESPONSE REGULATOR1* (*ARR1, CpaM1st05982*), and *PHYTOCHROME C* (*PHYC, CpaM1st08040*), which was associated with general environmental variables such as light regulation, temperature, and precipitation (Wei et al., 2021).

The reconstruction with the ancestral alleles from the diploid *C. paliurus* reference genome was used to estimate the site frequency spectrum for 45 *C. paliurus* accessions using ANGSD and to generate a stairway plot elucidating effective population size (*Ne*) history over time. With an estimate mutation rate of 2×10^−9^ per generation and a generation time of 8 years, the stairway plot revealed two population bottlenecks over deep time (**Supplementary Figure 44**). An early *Ne* bottleneck (dating back to ∼4.5-3.0 Mya) appeared during the upper Pliocene, consistent with the known events of environment change that boundary between the Pliocene and Miocene, was a regional transition from the warmer to the cooler stages (Zheng et al., 2000). The last *Ne* drop (dating back to ∼0.6-0.16 Mya) corresponds with the great extinction event at the Pleistocene glaciation (Paillard et al., 1998), followed by a rapid population expansion. Facilitated by the comparison of the demographic bottlenecks in three *Camellia sinensis*, we uncovered similar bottlenecks in *Cyclocarya paliurus* (0.6-0.16 Mya) and *C. sinensis* (2.5–0.7 Mya), both coinciding with known periods of environmental change (Zhang et al., 2021).

## Discussion

*Cyclocarya paliurus* is well-known as the sweet tea tree and traditionally used as an herbal medicine (Kakar et al., 2020). The diverse ploidy and heterodichogamy in this species make it an ideal model to investigate the functional impact of WGDs and the development of flowers. We have generated three references, including two diploid and one haplotype-resolved tetraploid genomes, by incorporating the newly developed sequencing technologies and chromosome phasing algorithm. Facilitated by the comparison of these genomes and 45 re-sequenced individuals, we are able to investigate the evolutionary history and uncover functional genes underlying environmental adaptation as well as factors contributing to enhanced photosynthesis and the biosynthesis of Cyclocaric acid B, one of the active components for the treatment of hypertension and diabetes (Zhu et al., 2015).

Studies have shown that *Cyclocarya* is an ancient genus, likely originating in the late Paleocene and becoming extinct in the Early Miocene except for *C. paliurus* from the subtropical China (Wu et al., 2017). The remaining *C. paliurus* is a relict plant species within the genus, serving as an excellent model to study the adaptive evolution of relict plants. Our population analysis uncovered two bottlenecks in the species, with each coinciding with dramatic climate changes. However, the rapid demographic decline was recovered by population expansion as shown in the stairway plot, posing questions about how this species survived during the long-term evolutionary history. Increasing evidence has shown that WGD is a pivotal contributor to adaptation in angiosperms (Wu et al., 2019). Two rounds of WGDs after the ancient WGT event shared by eudicots are observed in *C. paliurus*, occurring at ∼67.6-50.7 Mya and ∼11.2-10.5 Mya, respectively. Population genetics analysis clusters the ten diploid and 35 tetraploid re-sequenced individuals into two distinct groups, indicating one single origination of the latest WGD event rather than multiple WGD events in a different location. Comparison between the auto-tetraploid and diploid genomes shows that the dosage effect after the most recent WGD involved genes contributing to adaptive evolution and improvement of photosynthesis, such as genes in terpenoid metabolic biosynthetic pathways, carbohydrate, starch, and sucrose metabolism. This is highlighted by the biosynthesis of Cyclocaric acid B, which shows a significant increase in the auto-tetraploid genome compared to the diploid *C. paliurus*, attributing to elevated copy numbers of CYP716A, CYP72A and CYP71*A* subfamilies genes by WGDs. We also observed that the Pn, stomatal size, chlorophyll content in the leaves of tetraploid *C. paliurus* are significantly higher than diploid, which is a benefit to improve its growth and environment adaptability. Moreover, combining the experimental phenomenon, the pollen viability of diploid *C. paliurus* is significantly lower than that in tetraploid (**Supplementary Figure 45**). In conclusion, we propose that the tetraploid *C. paliurus* is more superior on the physiological and ecological characteristics than diploids.

As a typical heterodichogamy species, *C. paliurus* possess two complementary morphs with asynchronous flowering, which may effectively prevent selfing, reduce intramorph inbreeding, and heavily contribute to the pattern of genetic diversity in the process of species evolution (Bai et al., 2010). Our results uncover that GA_3_ content plays an important role in the asynchronous flowering, which is also evidenced by up-regulated expression of GA related genes in PG-F and PA-M. In addition to GA related signaling pathway, co-expression network analysis identify that hub genes, including *Trihelix-1, TCP, ERF090*, and *WRKY55*, likely contributing to heterodichogamy trait in *C. paliurus*.

## Methods

### Homologous chromosomes synapsis analysis of PA-tetra *C. paliurus*

The male florals at the early stage of meiosis of PA-tetra *C. paliurus* were transferred into Carnoy’s fluid (75% methanol, 25% glacial acetic acid) at 4°C for 24 hours under dark condition. Then 5 anthers were transferred to glass slide with 45% glacial acetic acid for 2 minutes acid hydrolysis. After covering the coverslip, the pollen mother cells were observed using phase contrast microscope, and the effective tablets were stored at −80°C for 24 hours. Then the 100% ethanol was added in materials at room temperature for 30 minutes dehydration. Finally, 15 ul DAPI was added and the tablet was examined by fluorescence microscope.

### Genome sequencing and assembly of three *C. paliurus*

The *C. paliurus* chromosome level assemblies combined three technologies from Single-Molecule Real-Time (SMRT) sequencing with the PacBio Sequel technology, chromatin conformation capture (Hi-C), and short reads polished based Illumina HiSeq sequencing. Briefly, ∼223× coverage for PA-dip, ∼115× coverage for PG-dip, and ∼221× coverage for PA-tetra raw data were generated with PacBio Sequel II platform and ∼176× coverage, ∼131× coverage, and ∼237× coverage, respectively by paired-end reads on the Illumina Novaseq6000 platform (**Supplementary Note**).

The initial contig-level assemblies were accomplished using a series of PacBio assemblers. More specifically, the longest coverage of subreads from PacBio SMRT Sequencer was self-corrected using CANU v.1.7 (Koren et al., 2017). Then, error-corrected reads were assembled into genomic contigs using three widely-used PacBio assembler CANU v.1.7 (Koren et al., 2017) with parameter corOutCoverage=100. Evaluation of N50s, the assembled genome size (**Supplementary Table 2**), and the complete BUSCO ratio (**Supplementary Table 6**) were used to inspect the quality of each round of assemblies. Last, the best results for subsequent analysis were selected through careful manual inspection. Illumina paired-end reads were further used to polish the PacBio assemblies using Pilon v.1.18 (Walker et al., 2014). Young leaves of *C. paliurus* were prepared for Hi-C libraries construction according to the standard protocol described previously (Belton et al., 2012). The paired-end sequencing libraries were generated from chimeric fragments, followed by Illumina sequencing. The paired-end Hi-C reads were aligned to the contig-level assembly and mis-joined contigs were then corrected using the 3D-DNA pipeline v.201008 (Dudchenko et al., 2017) for abrupt long-range contact patterns detecting. The contigs corrected by Hi-C interactions were successfully linked into 16 pseudo-chromosomes in PG-dip and PA-dip, and 64 pseudo-chromosomes with four sets of haplotypes in PA-tetra *C. paliurus* using the ALLHiC pipeline (Zhang et al., 2019), following the guideline that was used to assemble an auto-tetraploid sugarcane genome (https://github.com/tangerzhang/ALLHiC/wiki/ALLHiC:-scaffolding-an-auto-polyploid-sugarcane-genome).

### Genome annotation

#### RNA extraction and sequencing

Total RNA was isolated from stems, leaves, leaf buds and floral buds using E.Z.N.A Plant RNA Kit (OMEGA, USA), and then purified with RNase-Free DNase I (Takara Company, China). 1% agarose gel was used to evaluate the RNA contamination and degradation. The purity was further monitored using ultraviolet spectrophotometer (IMPLEN, CA, America). Samples with RIN values higher than 8 were used for downstream cDNA library preparation. The cDNA libraries construction was performed with NEBNext® UltraTM RNA Library Prep Kit according to the manufacturer’s instruction. The Agilent Bioanalyzer 2100 system was performed for library quality assessing and short paired-end reads were generated from the libraries preparations based on Illumina Novaseq sequencing platform.

#### Repeat annotation

Repetitive sequences were identified in the three *C. paliurus* genomes based on the same pipeline. First, RepeatModeler v.2.0.1 (see **URLs**) and RepeatMasker v.4.0.5 (see **URLs**) were used to *de novo* predict and discover the known transposable elements (TEs). Next, TE class v.2.1.3 (Abrusán et al., 2009) was further to categorize the unknown TEs. Then, two pipelines Tandem Repeat Finder (TRF) v.4.07 (Benson et al., 1999) and LTR_Finder v.1.05 (Xu & Wang, 2007) were performed to detect intact long terminal repeat (LTR) retrotransposons and tandem repeat, respectively. Finally, LTRharvest v1.5.10 (Ellinghaus et al., 2008) and LTR_retriever (Qu et al., 2018) were used to construct a high-quality LTR library.

#### Gene annotation

MAKER2 v.2.31.9 computational pipeline (Cantarel et al., 2008) was used to annotate genes in the three *C. paliurus* genomes, through a comprehensive strategy from RNA-seq-based prediction, homology-based prediction, and ab *initio* gene prediction. Briefly, RNA-seq data of different tissues of *C. paliurus* were assembled by Trinity v.2.6.5 (Haas et al., 2013) software with default parameters, genome-guided assembly and *de novo* assembly. The fragments per kilobase per million (FPKM) expression value of assembled transcripts was quantified by RSEM (Li & Dewey, 2011), the transcripts were removed if FPKM less than 1. The PASA v.r09162010 (Haas et al., 2003) program was applied to construct a comprehensive transcript library from the filtered transcripts. The almost “full-length” transcripts selected from PASA were aligned to the Uniprot protein database, and protein sequences with coverage greater than 95% were reserved as candidate sequences. Afterward, the MAKER2 pipeline was used to integrate coding evidence of three annotation strategies [SNAP v.29-11-2013. (Korf et al., 2004), GENEMARK v.4.28 (Lomsadze et al., 2005) and AUGUSTUS v.3.2.3 (Stanke et al., 2006)] and annotate protein-coding genes. After the first round, the predicated gene models with annotation edit distance (AED) values < 0.2 were selected for model re-training. Finally, the gene annotation improvement was generated from the second round of MAKER2. Further, the RNA-seq reads were aligned to reference genomes using HiSAT2 v.2.0.4 with default parameters and re-assembled by StringTie v.2.2.0 (Pertea et al., 2016). The assembled RNA-seq transcripts and homologous proteins from *Oryza sativa, Vitis vinifera, Carica papaya, Morus notablis, solanum tuberosum* and *Arabidopsis thaliana* were imported to MAKER2 pipeline. After filtering putative gene models of transposon-derived, a total of 34,699, 35,221, and 90,752 gene models were annotated in PA-dip, PG-dip and PA-tetra *C. paliurus*, respectively. BUSCO analyses for PA-dip, PG-dip, and PA-tetra *C. paliurus* were used to evaluate completeness of the protein-coding annotation (**Supplementary Table 6**).

### Phylogenetic tree reconstruction

To identify gene family groups, we analyzed protein-coding genes from 9 species, *C. paliurus, C. illinoinensis, J. nigra, P. stenoptera, A. thaliana, Ziziphus jujuba, Vitis vinifera, Populus trichocarpa*, and *Oryza sativa* genomes. Gene family clustering was performed using OrthoFinder v.2.2.7 (Emms et al., 2019) based on 35,221 predicted genes of *C. paliurus*, and the *O*.*sativa* as outgroup. Phylogenetic tree was constructed for *C. paliurus* and 8 other plant species based on coding sequence alignment of 302 single-copy gene families using Fasttree v. 2.1.11 (Price et al., 2009) software. The divergence time among 9 species was estimated by the r8s v.1.8.1 (Sanderson et al., 2003) program. For estimation of divergence time, we selected two calibration points from articles and calibrated the age of the nodes between *O. sativa* and *A. thaliana* (308-115 Mya), *J. nigra* and *P. stenoptera* (76-36 Mya), and *V. vinifera* and *A. thaliana* (135-107 Mya) according to the TimeTree website. The contraction and expansion of the gene families were observed by comparing the differences of cluster size between *C. paliurus* and each species using CAFE v.4.2.1 (Bie et al., 2006) method.

### Analysis of genome collinear and whole-genome duplication

For the comparative genomics analysis, the species we chose including C. paliurus, Pterocarya stenoptera, Juglans nigra, and Carya illinoinensis all belong to the family Juglandaceae. The Ziziphus jujuba shares the same typical feature of heterodichogamy with Juglandaceae. Arabidopsis thaliana and Populus trichocarpa were also chosen as the model plants for comparison. Vitis vinifera is a basal eudicot that has experienced an known whole-genome triplication (WGT) event, without a recent independent whole-genome duplication. Oryza sativa as a monocotyledon, is used as an outgroup. Hence, the genomic information of these plant species are significant for comparative genomics analysis. We performed collinearity searches to identify collinear blocks within *C. paliurus* using MCScanX v.1.1.11(Wang et al., 2012). Synonymous substitutions per synonymous site (*Ks*) between collinear genes were estimated based on WGDI v.0.1.6 (Sun et al., 2021) software with YN model. In brief, the collinear blocks were constructed by performing similarity search for all-against-all protein sequence using BLASTP v.2.8.1 (Altschul et al., 1997) (cutoff *E* value of 1e-10), then homologous block was built through MCScanX software. Finally, the *Ks* of each homologous gene pair was calculated.

### Analysis of the tetraploidy signatures

#### Identification of chromosome-enriched K-mers

To determine whether the genome is auto-tetraploid or allo-tetraploid, we adopt and modified a similar method that was used to study an allo-tetraploid *Miscanthus sinensis* genome (Mitros et al., 2020) based on counting of chromosome-enrich 13-mers. Briefly, 13-mers were identified across the whole genome using Jellyfish v.2.2.6 (Marcais et al., 2011) and only *K*-mers that were meet the following two conditions were retained: 1) *K*-mers occurring at least 1,000 times globally; 2) *K*-mers that were enriched in any chromosome with at least two-fold difference. Applying of the filtering strategy resulted in a total of 11,783 chromosome-enriched 13-mers. We further clustered these selected *K*-mers basing on the number of these 13-bp short sequences presenting across the whole genome. The results showed that every homologous chromosome group that comprises of 4 haplotypes were grouped together, providing strong evidence of auto-tetraploidy signatures.

#### Smudge plot analysis

We also performed the smudge plot analysis (Ranallo et al., 2020) to investigate the polyploid signatures in this tetraploid species. This tool utilized heterozygous paired *K*-mers extracted from raw reads in the sequenced genome and analyzed the genome structure by comparing the total coverage of paired *K*-mers (i.e., coverage A+coverage B) to the relative coverage of the minor one (i.e., coverage B/ (coverage A+coverage B), where A and B are paired heterozygous *K*-mers, A indicate the dominant *K*-mer and B is the minor one).

### Identification of differentially expressed genes between PG and PA

Male and female floral buds in various protandry (PA) and protogyny (PG) individuals were collected at five different stages including 1) S0; physiological differentiation period, 2) S1; dormancy period, 3) S2; germination period, 4) S3; inflorescence elongation period; and (5) S4; maturation period. Three biological replicates of the extracted RNA from flowers (in PA and PG individuals) were sequenced on Illumina Novaseq platform, and 6 Gb RNA-seq raw data for each sample was generated. Paired-end short reads were aligned to PG-dip *C. paliurus* genome using Hisat2 (Kim et al., 2015). The expected number of FPKM fragments mapped were calculated using RSEM v.1.3.0 (Li & Dewey, 2011) program, which was implanted in Trinity package (Haas et al., 2013). Moreover, the DESeq2 R package v.1.30.0 (Love et al., 2014) was applied to identify the DEGs. The clusterProfiler R package v.3.12.0 (Yu et al., 2012) was used for GO and KEGG enrichment analysis.

### Genome screening for P450s and gene clusters

The HMM (hidden Markov model) profile for the P450s (PF00067) was obtained from the Pfam database (see **URLs**). Then, the P450s were identified using the HMMER v3.3.2 (Prakash et al., 2017) with default parameters by searching against the *C. paliurus* genome. According to definite standards with 40% for family variants described by Xiong et al. (2021), P450s were divided into 68 families by alignment with P450 database (Nelson et al., 2009). Maximum likelihood phylogenetic trees were constructed using the RAxML package v.8.2.11 (Stamatakis et al., 2006) with full-length protein sequences.

The GFF (General Feature Format) files of 42 expanded P450s were used to search for gene clusters. The gene clusters were verified by following criteria: one gene cluster should contain at least three adjacent P450s; the gene clusters were be ruled out if the distance between adjacent P450s were more than 0.8 Mb.

### Identification of expressed genes and triterpenoid compounds for *C. paliurus*

Leaves in various PA-dip and PA-tetra individuals were collected in May and September, respectively. Three biological replicates of the extracted RNA were sequenced on Illumina Novaseq platform, and 6 Gb RNA-seq raw data for each sample was generated. Paired-end short reads were aligned to PA-dip *C. paliurus* genome using Hisat2. Meanwhile, the same leaf samples were prepared for UPLC-MS/MS analysis. UPLC-MS/MS system (UPLC, SHIMADZU Nexera X2; MS/MS, Applied Biosystems 4500 Q TRAP) was used to analyse the differences in triterpenoids accumulation between diploid and tetraploid plant leaves. Sample extraction details were described in Supplementary Note. The UPLC operation parameters were as follows: Chromatographic separation was carried out using an Agilent SB-C18 HPLC analytical column (2.1 mm ×100 mm, 1.8 µm); the mobile phase consists of ultrapure water (A) and acetonitrile (B), which both contains 0.1% formic acid. Gradient elution was as follows: original ratio of 5% B; B ratio linearly increases to 95% within 9 min and maintained for 1 min; B ratio decreased to 5% during 10 to 11.1 min and kept for 2.9 min. The flow rate was kept at 0.35 ml/min, temperature 40°C, and injection volume 4µL. The electrospray ionization (ESI)-triple quadrupole-linear ion trap (Q TRAP)-MS system were carried out for MS experiment. The operating parameters of the ESI source were as follows: positive ion spray voltage (IS) 5500V and negative ion mode −4500 V; source temperature 550°C; curtain gas (CUR) 25 psi, ion source gas I (GSI) 50 psi, and gas I (GSII) 60 psi, respectively. The 10 and 100μmol/L polypropylene glycol solution were respectively implemented for mass calibration and instrument tuning in linear ion trap (LIT) modes and triple quadrupole (QQQ).

### Population genetic structure

More than coverage of 10× per sample for tetraploid and diploid *C. paliurus* were generated from 74 and 35 billion 150-bp Illumina short reads, respectively. Clean data was obtained by removing adapters and low-quality sequences (Q < 30) from paired-end raw reads, followed by aligning against the reference genome of PA-dip *C. paliurus* by bwa v.0.7.17 (Li et al., 2009) with default parameters. The variant calling was carried out by GATK v.4.0.3.0 (Mckenna et al., 2010) following the best practices workflow. The general variants were identified for each individual using GATK HaplotypeCaller, and then combined by GenotypeGVCFs function to a single variant calling file. This two-step approach was carried out to ensure variant accuracy, which included re-genotyping and quality recalibration in the combined vcf file. SNPs were then identified using samtools/bcftools with default parameters based on alignments of all Illumina short reads. SNPs were filtered by following parameters: 1) SNPs were only present in one of the two pipelines (SAMtools/BCFtools and GATK); 2) SNPs with read depth more than 1,000 or less than 5; 3) non-biallelic SNPs; 4) SNPs with missing rate more than 40%; 5) SNPs in repeat regions; 6) SNPs were removed if the distance with nearby variant sites less than 5bp. A phylogenetic gene tree was constructed based on SNPs in the single-copy genes regions. Two popular programs [RAxML (Stamatakis et al., 2014) with GTRCAT model and IQ-Tree (Nguyen et al., 2015) with self-estimated best substitution model] were applied to construct the maximum likelihood (ML) tree. Ancestral population structure among nine *C. paliurus* populations was estimated from ancestral population sizes K = 1-5 by ADMIXTURE software v.1.3.0 and the population size with the smallest cross-validation error (K = 2) was determined. Admixture analysis was performed following the parameter standard errors, which estimated by bootstrapping (bootstrap = 200).

### Identification of selective sweeps

To identify selective sweeps, SweeD v.3.0 (Pavlidis et al., 2013) program was used to identify regions that display significant variations in the Site Frequency Spectrum (SFS) by the composite likelihood ratio (CLR) statistic. 0.3 percent with maximum missing data was allowed in per site, and top 5 percent as the threshold to screen candidate selective sweeps. Ten diploid and 35 tetraploid individuals were compared to PA-dip and PA-tetra reference genomes, respectively.

## Supporting information

supplemental material_preprint

## URLs

RepeatModeler, http://www.repeatmasker.org/RepeatModeler/; RepeatMasker, http://www.repeatmasker.org/; Estimate_genome_size.pl, https://bioinformatics.uconn.edu/genome-size-estimation-tutorial/; Pfam, http://pfam.xfam.org/.

## Acknowledgments

This work was funded by the Natural Science Foundation of China (No. 31971642, 32071750, and 31470637). This work was also supported by the Key Research and Development Program of Jiangsu Province (BE2019388) and the Priority Academic Program Development of Jiangsu Higher Education Institutions (PAPD). We received editing assistance from Life Science Editors.

## Author contributions

Shengzuo Fang, Xiangxiang Fu, Yinquan Qu, and Xingtan Zhang designed this genome project and coordinated research activities. Yinquan Qu, Xulan Shang, Yanhao Yu, Li Liu, Qian Wang, Caowen Sun, Wanxia Yang, and Shengzuo Fang collected and provided plant materials. X.S., Guoliang Bian and Yinquan Qu performed the Karyotype anaysis and flow cytometry. Ziyan Zeng performed the figures and tables in paper. Wenling Wang and Yinquan Qu performed population genetics analysis. Xingtan Zhang and Shengcheng Zhang assembled, annotated and analyzed the genomes. Yinquan Qu, Ddejin Xie, and Xiangxiang Fu performed RNA-seq analysis and identified genes associated with heterodichogamy. Yinquan Qu and Shengzuo Fang identified the key enzyme genes in the triterpenoids synthesis pathway. Xuequn Chen completed statistics of genome information. Zhengyang Liao performed the polyploidy evolution. Yibin Wang performed auto-tetraploid identify. Jian Qin modified the format of text. Xingtan Zhang, Yinquan Qu, Xiangxiang Fu and Shengzuo Fang interpreted the data and contributed to the manuscript writing.

## Data Accessibility

The whole genome sequencing raw data including Illumina short reads, PacBio long reads, Hi-C interaction reads, and transcriptome data have been submitted to the Genome Sequence Archive (GSA) database, under the accession number CRA004671 and Bioproject accession number: PRJCA00598727. The genome assemblies and annotation were temporarily deposited in Google Drive (https://drive.google.com/file/d/1YDqAk_jvoX_ilb7kSxqs56QfbEiHj9Nm/view?usp=sharing) and Baidu cloud (https://pan.baidu.com/s/1XoKEQAjYw0GFgDYvdRT7aw, extraction number: bm0u) for review. We are also submitting the assemblies, annotation and sequencing short reads to NCBI and GSA databases, however, still waiting for accession numbers. All of the data generated in this study will be publicly released once our manuscript is accepted.

