## supplemental material_preprint for "Whole-genome duplication reshaped adaptive evolution in a relict plant species, *Cyclocarya paliurus*"

### Supplementary Notes

**Genome sequencing and assembly of three *C. paliurus***

***Plant material.*** Tissues of two diploid *C. paliurus* (PG-dip and PA-dip) and one autotetraploid (PA-tetra) for genome sequencing were collected from plants grown in germplasm bank of *C. paliurus*, which locates in Baima experimental, Nanjing, Jiangsu province, China. Tissues were immediately frozen in liquid nitrogen and stored at -80℃.

***Illumina short reads sequencing.*** Total DNA was isolated and extracted using the QIAGEN DNaesy Plant Mini Kit and sequenced in Illumina Novaseq6000 with 150-bp paired-end (PE) reads length.

***Pacbio library construction and Sequencing.*** Approximately 20 μg of high-molecular-weight genomic DNA was sheared to an ~20 kb targeted size, followed by damage repair and end repair, blunt-end adaptor ligation and size selection with a Blue Pippin system. The final libraries were subsequently sequenced using a PacBio Sequel II with P6-C4 chemistry. After data filtering and preprocessing, 134.9, 75.5, and 271.8 Gb raw data were generated for PA-dip, PG-dip, and PA-tetra *C. paliurus*, respectively (**Supplementary Table 1**).

***Hi-C library construction and sequencing.*** Hi-C libraries were created from tender leaves of *C. paliurus* plants as described before (Dudchenko et al., 2017). Briefly, the leaves were fixed with lysed, formaldehyde, and then the cross-linked DNA digested with *HindIII* over-night. Sticky blunt ends were biotinylated and proximity-ligated to form chimeric junctions, and then physically sheared to a size of 500-700 bp. Chimeric fragments representing the original cross-linked long-distance physical interactions were then processed into paired-end sequencing libraries. A total of 196, 190, and 687 million of 150-bp paired-end reads were produced on Illumina Novaseq6000 system. Hi-C sequencing was assessed using HiC-Pro program and the results showed a decent proportion of validate reads (no less than 80% in three *C. paliurus* Hi-C libraries), suggesting high quality of the Hi-C data (**Supplementary Table 3**).

***Estimation of C. paliurus genome size.*** To estimate the genome size of the three *C. paliurus*, Illumina short reads were recruited to determine the distribution of *K*-mer values using published PERL scripts (see **URLs**). The genome size was estimated using the following formula: the number of *K*-mers divided by the depth of peaks (*K*-mer-number/depth). The size of the haploid genomes for PA-dip and PG-dip *C. paliurus* were estimated to be approximately 538.16 Mb and 574.97 Mb, which are similar with the genome size of *P. stenoptera*. In addition, the estimated genome size is 2065.41 Mb, and high heterozygosity that comes from PA-tetra *C. paliurus*. To obtain estimation of nuclear DNA content, we followed previous method (Loureiro et al. 2007) to perform flow cytometry. The estimated genome sizes are consistent with our *K*-mer based estimation with an average 606 Mb for PA-dip, 659 Mb for PG-dip, and 2460 Mb for PA-tetra *C. paliurus* (**Supplementary Figure 3**).

**Estimation of genome heterozygosity rate.** Ideally each *K*-mer in a set of genomes should be a single occurrence, and there is no repetitive sequence or heterozygosity. In actual samples, due to the influence of heterozygous and repetitive sequences, the corresponding frequency of each *K*-mer is uncertain. We classified *K*-mer according to the frequency of appearance (i). Then, We calculated the percentage of each type of *K*-mer number: $a_i = n_{i,Kspecies}/n_{Kspecies}$, and the percentage of the number: $bi = n_{i,Kindividuals}/n_{Kindividuals}$. Considering the distribution of *K*-mer frequencies in the genome is [1, m], and the expected depth of each category is $C_i = i*C$, the probability function of the number of *K*-mer species in the genome with heterozygosity and repetitive sequences could be obtained ($P_{Kspecies}(x)$ represents the Poisson distribution with the desired depth Ci):

$$\mathbf{P_{Kspecies}(x) = \sum_{i=1}^{m} a_i \times P_{Kspecies,i}(x)}$$

The probability function of the number of *K*-mers in a genome with heterozygosity and repetitive sequences could be obtained ($P_{Kindividuals}(x)$ represents the deformed Poisson distribution with the desired depth Ci.):

$$\mathbf{P_{Kindividuals}(x) = \sum_{i=1}^{m} b_i \times P_{kindividuals,i}(x)}$$

For heterozygous genomes, all *K*-mers can be divided into two categories: heterozygous *K*-mers and homozygous *K*-mers. There are 2*K heterozygous *K*-mers covered at each heterozygous site. Hence, the expected depth of the heterozygous *K*-mer is C/2 compared to the expected depth of the homozygous *K*-mer.

Using this method, the number of heterozygous sites and genome size could be estimated by $a_{1/2}$ and $n_{Kspecies}$, and the heterozygosity rate was estimated to be 1.97% ($a_{1/2}$ is the percentage of the number of heterozygous *K*-mer species, $n_{Kspecies}$ is the number of all *K*-mer types):

$$\phi = \frac{a_{1/2} \times n_{Kspecies} /(2 \times K)}{n_{Kspecies} - a_{1/2} \times n_{Kspecies} / 2} = \frac{a_{1/2}}{K(2 - a_{1/2})}$$

**Identification of triterpenoid compounds for *C. paliurus***

Standards (Chromatographic purity) were purchased from Sigma-Aldrich (St Louis, MO, USA), MeOH and ACN were all purchased from Merck (Darmstadt, Germany). *C. paliurus* leaves of vacuum freeze-drying were ground into powder (30 Hz, 1.5min), and 100 mg powder were extracted with 1.2 mL methanol solution (70%). The extract was vortexed every 30 minutes and repeat in six sets of 30 seconds, and stored at -20 °C for one night. After centrifugation (rotating speed 12,000 rpm, 10 min), the supernatants were filtered with microporous membrane (0.22 μm pore size) and collected for UPLC-ESI-MS/MS analysis.

***Hi-C scaffolding and chromosome assembly.*** Hi-C reads were uniquely mapped to the contig assemblies and reads within 500 bp regions of *Sau3*AI restriction sites were retained for further analysis. Mis-joined contigs were corrected to detect abrupt long-range contact patterns by 3D-DNA pipeline. The Hi-C corrected contigs were further linked into 16 pseudo-chromosomes in PG-dip and PA-dip, and 64 pseudo-chromosomes with 4 sets of monoploid chromosomes in PA-tetra *C. paliurus* using the ALLHiC pipeline (Zhang et al. 2019). The accuracy of Hi-C based chromosome construction was evaluated by chromatin contact matrix (**Supplementary Figures 4-6**).

**Phylogenetic analysis of P450s subfamilies**

Phylogenetic analysis revealed that 22 dosage-effect genes were clustered into their respective clade for plant P450s family. Furthermore, seven genes clustered to *CYP716A* (*CpaM1st27342* and *CpaM1st17575*), *CYP71A* (*CpaM1st23492* and *CpaM1st07944*), and *CYP72A* (*CpaM1st19063*, *CpaM1st29930*, *and CpaM1st29927*) in *C. paliurus* were deeply compared to public databases by BLAST. Among them, *CpaM1st27342* and *CpaM1st17575* were defined as the ortholog genes of *CYP716A1*, *CYP716A2* in *Arabidopsis thaliana,* respectively. *CpaM1st23492* and *CpaM1st07944* were highly homologous with *CYP71A22* and *CYP71A26,* respectively. *CpaM1st19063* and *CpaM1st29927* were highly homologous with *CYP72A219*. *CpaM1st29930* was an ortholog of *CYP72A15*. For whole genome scale, *CpaM1st27346* and *CpaM1st17121* were defined as *CYP716A14v2* gene in *Artemisia annua, and CpaM1st38811* was an ortholog of *CYP716C* in *A. thaliana*.

**Supplementary Table 1. Sequencing information for the assemblies.**

| Library type | Insert size | Raw data (Gb) | | | Coverage (×) | | |
|---|---|---|---|---|---|---|---|
| | | PA-dip | PG-dip | PA-tetra | PA-dip | PG-dip | PA-tetra |
| Illumina Paired-end | 200 bp | 106.7 | 86 | 291.7 | 176 | 131 | 237 |
| PacBio Sequel II | 20 kb | 134.9 | 75.5 | 271.8 | 223 | 115 | 221 |
| Hi-C | 200 bp | 65.4 | 68 | 264 | 108 | 103 | 215 |
| Total | - | 307 | 229.5 | 827.5 | 507 | 348 | 673 |

**Supplementary Table 2. Contig level assembly.**

| | PA-dip | PG-dip | PA-tetra |
|---|---|---|---|
| No. of contigs | 1101 | 921 | 9,744 |
| Max length (Mb) | 8.68 | 12.50 | 10.32 |
| Assembly size (Mb) | 586.62 | 583.45 | 2380.95 |
| Contig N90 (bp) | 419,278 | 355,229 | 100,226 |
| Contig N50 (bp) | 1,928,354 | 1,389,753 | 430,910 |
| Average (bp) | 532,809 | 633,498 | 244,350 |
| Total No. of contigs (>2kb) | 1,082 | 912 | 9,719 |

**Supplementary Table 3. Statistics of Hi-C mapping**

| | PG-dip | PA-dip | PA-tetra |
|---|---|---|---|
| **Statistics of mapping** | | | |
| Clean Paired-end Reads | 195501972 | 190011721 | 686905160 |
| Unmapped Paired-end Reads | 1793734 | 2033035 | 1986108 |
| Unmapped Paired-end Reads Rate (%) | 0.918 | 1.07 | 0.289 |
| Paired-end Reads with Singleton | 25285318 | 25084739 | 37399248 |
| Paired-end Reads with Singleton Rate (%) | 12.934 | 13.202 | 5.445 |
| Multi Mapped Paired-end Reads | 54214489 | 57828217 | 541930877 |
| Multi Mapped Ratio (%) | 27.731 | 30.434 | 78.895 |
| Unique Mapped Paired-end Reads | 114208431 | 105065730 | 105588927 |
| Unique Mapped Ratio (%) | 58.418 | 55.294 | 15.372 |
| **Statistics of valid reads** | | | |
| Unique Mapped Paired-end Reads | 114208431 | 105065730 | 105588927 |
| Dangling End Paired-end Reads | 1375950 | 1597284 | 8185832 |
| Dangling End Rate (%) | 1.205 | 1.52 | 7.753 |
| Self-Circle Paired-end Reads | 125793 | 94131 | 87361 |
| Self-Circle Rate (%) | 0.11 | 0.09 | 0.083 |
| Dumped Paired-end Reads | 125793 | 94131 | 87361 |
| Dumped Rate (%) | 0.11 | 0.09 | 0.083 |
| Interaction Paired-end Reads | 101609002 | 92261311 | 85094810 |
| Interaction Rate (%) | 88.968 | 87.813 | 80.591 |
| Lib Valid Paired-end Reads | 94294785 | 85581476 | 77436384 |
| Lib Valid Rate (%) | 88.968 | 87.813 | 80.591 |
| Lib Dup (%) | 6.404 | 6.358 | 7.253 |

**Supplementary Table 4. Chromosomal level genome assemblies.**

| | PA-dip (Mb) | PG-dip (Mb) | PA-tetra (Mb) | | | |
|---|---|---|---|---|---|---|
| **ChrID** | | | **Hap A** | **Hap B** | **Hap C** | **Hap D** |
| Chr1 | 46.29 | 46.58 | 41.27 | 38.01 | 42.45 | 50.08 |
| Chr2 | 43.74 | 44.40 | 43.92 | 43.74 | 32.86 | 50.40 |
| Chr3 | 39.60 | 41.58 | 41.77 | 41.49 | 38.96 | 38.16 |
| Chr4 | 38.01 | 38.67 | 39.44 | 42.99 | 36.33 | 35.67 |
| Chr5 | 37.99 | 37.97 | 37.28 | 39.01 | 39.72 | 33.65 |
| Chr6 | 36.49 | 37.30 | 37.88 | 37.41 | 35.25 | 35.28 |
| Chr7 | 36.56 | 37.08 | 44.92 | 35.67 | 40.88 | 25.69 |
| Chr8 | 32.70 | 35.15 | 42.35 | 36.54 | 32.56 | 34.12 |
| Chr9 | 33.84 | 34.68 | 33.98 | 31.55 | 34.36 | 34.52 |
| Chr10 | 32.37 | 32.69 | 31.48 | 32.99 | 31.72 | 30.89 |
| Chr11 | 33.71 | 32.21 | 33.96 | 38.48 | 36.88 | 23.04 |
| Chr12 | 29.00 | 30.21 | 32.31 | 29.82 | 23.49 | 31.52 |
| Chr13 | 28.95 | 29.25 | 27.51 | 29.46 | 35.23 | 22.26 |
| Chr14 | 25.93 | 27.22 | 27.58 | 27.05 | 26.89 | 26.16 |
| Chr15 | 24.52 | 24.85 | 25.30 | 24.02 | 25.61 | 24.33 |
| Chr16 | 23.82 | 24.03 | 27.91 | 20.92 | 23.21 | 18.46 |
| Total length of contigs (Mb) | 586.62 | 583.45 | | | | 2380.95 |
| Total length of chromosome level assembly (Mb) | 543.53 | 553.87 | | | | 2168.65 |
| Anchor rate (%) | 92.65 | 94.93 | | | | 91.08 |

**Supplementary Table 5. Characteristics of genetic variation compared with monoploid genome in PA-tetra *C. paliurus* (Chr1-Chr8)**

| Haplotype | Variation | Chr1 | Chr2 | Chr3 | Chr4 | Chr5 | Chr6 | Chr7 | Chr8 |
|---|---|---|---|---|---|---|---|---|---|
| | SNPs | 247,084 | 273,640 | 214,901 | 259,224 | 195,550 | 231,148 | 251,540 | 201,220 |
| | No. of Indels(1-50bp) | 28,691 | 32,237 | 30,410 | 29,224 | 23,094 | 34,747 | 37,259 | 29,180 |
| | No. of large Indels(>50bp) | 189 | 166 | 171 | 182 | 161 | 242 | 243 | 219 |
| | Size of Indels(1-50bp) | 79,462 | 92,924 | 39,625 | 82,812 | 65,852 | 45,131 | 48,610 | 37,625 |
| | Size of Indels(>50bp) | 1,724,06 | 1,775,361 | 237,599 | 1,922,516 | 793,481 | 316,305 | 362,227 | 313,569 |
| A | No. of Repeat expansion/contraction | 728 | 762 | 19 | 688 | 559 | 20 | 42 | 14 |
| | Size of Repeat expansion/contraction | 21,838,7 | 27,314,80 | 44,820 | 29,455,39 | 16,657,83 | 33,525 | 55,352 | 30,237 |
| | SNPs | 232,152 | 244,049 | 205,512 | 232,980 | 215,731 | 270,483 | 259,684 | 219,201 |
| | No. of Indels(1-50bp) | 27,206 | 28,914 | 22,438 | 34,833 | 24,662 | 30,335 | 30,780 | 23,170 |
| | No. of large Indels(>50bp) | 194 | 183 | 172 | 208 | 148 | 191 | 213 | 460 |
| | Size of Indels(1-50bp) | 74,728 | 82,546 | 61,345 | 44,319 | 72,552 | 81,780 | 85,730 | 70,067 |
| | Size of Indels(>50bp) | 251,611 | 1,348,559 | 1,608,094 | 328,247 | 521,757 | 746,460 | 1,976,019 | 1,495,230 |
| B | No. of Repeat expansion/contraction | 425 | 734 | 535 | 25 | 627 | 705 | 737 | 590 |
| | Size of Repeat expansion/contraction | 24,707,9 | 32,067,05 | 20,458,59 | 58,905 | 21,037,94 | 17,777,06 | 26,863,05 | 18,863,84 |
| | SNPs | 269,864 | 204,468 | 198,378 | 201,211 | 215,496 | 215,327 | 250,694 | 174,369 |
| | No. of Indels(1-50bp) | 30,923 | 24,034 | 20,638 | 23,853 | 32,673 | 26,210 | 28,349 | 19,087 |
| | No. of large Indels(>50bp) | 234 | 165 | 135 | 145 | 243 | 177 | 178 | 128 |
| | Size of Indels(1-50bp) | 87,642 | 65,167 | 58,059 | 63,514 | 42,158 | 72,556 | 79,567 | 53,719 |
| | Size of Indels(>50bp) | 989,417 | 500,171 | 480,640 | 1,411,969 | 304,556 | 1,221,549 | 2,550,433 | 590,166 |
| C | No. of Repeat expansion/contraction | 869 | 615 | 567 | 553 | 33 | 598 | 694 | 423 |
| | Size of Repeat expansion/contraction | 27,331,7 | 17,847,63 | 19,302,04 | 25,253,11 | 38,521 | 22,445,69 | 26,379,66 | 20,308,99 |
| | SNPs | 259,901 | 249,763 | 206,110 | 197,296 | 216,555 | 185,559 | 196,663 | 174,885 |
| | No. of Indels(1-50bp) | 38，586 | 38,552 | 22,359 | 23,660 | 27,141 | 25,240 | 22,293 | 20,225 |
| | No. of large Indels(>50bp) | 265 | 243 | 152 | 149 | 180 | 148 | 143 | 115 |
| | Size of Indels(1-50bp) | 51，012 | 49,555 | 62,028 | 62,038 | 74,918 | 62,306 | 61,094 | 54,498 |
| | Size of Indels(>50bp) | 318,450 | 341,187 | 914,262 | 491,312 | 909,661 | 402,157 | 410,094 | 893,222 |
| D | No. of Repeat expansion/contraction | 34 | 34 | 557 | 572 | 632 | 554 | 575 | 465 |
| | Size of Repeat expansion/contraction | 92，999 | 81,902 | 20,441,34 | 22,573,70 | 24,047,21 | 18,868,87 | 15,252,79 | 20,180,20 |

**Supplementary Table 5. Characteristics of genetic variation compared with monoploid genome in PA-tetra *C. paliurus* (Chr9-Chr16)**

| Haplot | Variation | Chr9 | Chr10 | Chr11 | Chr12 | Chr13 | Chr14 | Chr15 | Chr16 |
|---|---|---|---|---|---|---|---|---|---|
| | SNPs | 189,675 | 227,892 | 213,606 | 190,418 | 168,096 | 154,562 | 137,768 | 158,013 |
| | No. of Indels(1-50bp) | 22,067 | 23,954 | 23,056 | 28,533 | 19,100 | 22,678 | 15,419 | 23,421 |
| | No. of large Indels(>50bp) | 139 | 117 | 171 | 173 | 127 | 135 | 104 | 185 |
| | Size of Indels(1-50bp) | 58,845 | 66,340 | 64,758 | 37,108 | 53,486 | 29,870 | 41,476 | 31,187 |
| | Size of Indels(>50bp) | 962,431 | 218,625 | 859,936 | 286,170 | 596,097 | 215,875 | 316,853 | 265,111 |
| **A** | No. of Repeat expansion/contraction | 5,287,368 | 630 | 566 | 21 | 499 | 27 | 389 | 29 |
| | Size of Repeat expansion/contraction | 25,269,476 | 16,767,825 | 21,099,563 | 68,411 | 20,290,338 | 101,962 | 5,434,321 | 38,911 |
| | SNPs | 177,427 | 226,131 | 211,135 | 176,264 | 178,162 | 142,865 | 135,328 | 152,941 |
| | No. of Indels(1-50bp) | 20,023 | 32,673 | 29,694 | 20,655 | 21,011 | 15,229 | 15,547 | 18,564 |
| | No. of large Indels(>50bp) | 145 | 150 | 202 | 130 | 146 | 107 | 107 | 126 |
| | Size of Indels(1-50bp) | 55,285 | 43,441 | 38,376 | 58,015 | 59,620 | 42,723 | 42,765 | 49,345 |
| | Size of Indels(>50bp) | 1,296,276 | 234,532 | 2,797,225 | 1,066,474 | 761,442 | 318,129 | 477,268 | 444,471 |
| **B** | No. of Repeat expansion/contraction | 3,867,662 | 22 | 27 | 524 | 502 | 418 | 377 | 412 |
| | Size of Repeat expansion/contraction | 25,269,476 | 41,523 | 32,970 | 15,460,267 | 20,262,057 | 11,884,804 | 13,046,205 | 12,491,353 |
| | SNPs | 199,733 | 220,365 | 225,310 | 160,280 | 165,371 | 139,320 | 140,654 | 175,543 |
| | No. of Indels(1-50bp) | 22,617 | 24,121 | 23,058 | 19,194 | 24,296 | 15,480 | 20,528 | 20,752 |
| | No. of large Indels(>50bp) | 156 | 148 | 157 | 121 | 186 | 109 | 115 | 113 |
| | Size of Indels(1-50bp) | 63,619 | 64,676 | 67,378 | 51,872 | 32,196 | 41,699 | 27,101 | 56,179 |
| | Size of Indels(>50bp) | 1,358,312 | 352,356 | 2,092,302 | 386,661 | 353,247 | 620,705 | 161,771 | 357,232 |
| **C** | No. of Repeat expansion/contraction | 536 | 629 | 595 | 491 | 27 | 465 | 20 | 483 |
| | Size of Repeat expansion/contraction | 18,846,793 | 16,678,274 | 23,146,240 | 14,608,217 | 59,179 | 14,468,290 | 37,831 | 17,462,352 |
| | SNPs | 203,320 | 204,460 | 147,127 | 202,337 | 128,868 | 147,385 | 134,194 | 132,600 |
| | No. of Indels(1-50bp) | 29,836 | 21,665 | 16,955 | 24,390 | 15,239 | 15,457 | 15,120 | 16,183 |
| | No. of large Indels(>50bp) | 181 | 165 | 106 | 155 | 99 | 128 | 109 | 109 |
| | Size of Indels(1-50bp) | 39,041 | 59,138 | 44,970 | 68,096 | 40,819 | 41,515 | 41,892 | 42,411 |
| | Size of Indels(>50bp) | 244,615 | 771,103 | 372,033 | 741,573 | 825,236 | 817,971 | 840,415 | 419,681 |
| **D** | No. of Repeat expansion/contraction | 29 | 592 | 380 | 607 | 390 | 409 | 372 | 389 |
| | Size of Repeat expansion/contraction | 52,274 | 13,895,575 | 14,101,988 | 20,242,633 | 16,116,574 | 10,475,463 | 13,467,245 | 16,634,406 |

**Supplementary Table 6. BUSCO assessment of genome assemblies.**

| Description | PA-dip | | PG-dip | | PA-tetra | |
|---|---|---|---|---|---|---|
| | Number | Percentage(%) | Number | Percentage(%) | Number | Percentage(%) |
| Complete BUSCOs(C) | 1308 | 95.2 | 1326 | 96.4 | 1312 | 95.5 |
| Complete and single-copy BUSCOs(S) | 1178 | 85.7 | 1194 | 86.8 | 181 | 13.2 |
| Complete and duplicated BUSCOs(D) | 130 | 9.5 | 132 | 9.6 | 1131 | 82.3 |
| Fragmented BUSCOs(F) | 14 | 1.0 | 9 | 0.7 | 8 | 0.6 |
| Missing BUSCOs(M) | 53 | 3.8 | 40 | 2.9 | 55 | 3.9 |
| Total BUSCO groups searched | 1375 | 100 | 1375 | 100 | 1375 | 100 |

**Supplementary Table 7. Statistics of Illumina short reads remapped to the assemblies.**

| | Numbers | | | Percentage | | |
|---|---|---|---|---|---|---|
| | **PA** | **PG** | **PA-tetra** | **PA** | **PG** | **PA-tetra** |
| Total reads | 315,235,824 | 313,292,807 | 1,321,149,523 | 100.00% | 100.00% | 100.00% |
| Mapped reads | 311,747,061 | 310,862,209 | 1,314,624,201 | 98.89% | 99.22% | 99.51% |
| Paired reads | 311,704,982 | 311,704,982 | 1,318,071,852 | 98.88% | 99.49% | 99.77% |
| Properly paired | 292,135,204 | 299,878,500 | 1,278,400,172 | 93.72% | 96.21% | 96.99% |
| Unmapped reads | 1,073,915 | 667,698 | 1,242,784 | 0.34% | 0.21% | 0.09% |

**Supplementary Table 8. Allele annotation in Auto-tetraploid *C. paliurus* genome.**

| | Total no. of genes | No. of Genes with 4 alleles | No. of Genes with 3 alleles | No. of Genes with 2 alleles | No. of Genes with 1 allele |
|---|---|---|---|---|---|
| Chr1 | 2,777 | 608 | 913 | 669 | 587 |
| Chr2 | 2,851 | 639 | 924 | 748 | 540 |
| Chr3 | 2,035 | 608 | 486 | 415 | 526 |
| Chr4 | 2,349 | 639 | 743 | 531 | 436 |
| Chr5 | 2,598 | 548 | 913 | 642 | 495 |
| Chr6 | 2,277 | 612 | 717 | 489 | 459 |
| Chr7 | 2,364 | 848 | 705 | 430 | 381 |
| Chr8 | 2,229 | 372 | 595 | 543 | 719 |
| Chr9 | 2,093 | 639 | 608 | 426 | 420 |
| Chr10 | 2,210 | 769 | 619 | 395 | 427 |
| Chr11 | 2,405 | 623 | 589 | 446 | 747 |
| Chr12 | 1,912 | 471 | 699 | 381 | 361 |
| Chr13 | 1,924 | 470 | 632 | 453 | 369 |
| Chr14 | 1,663 | 591 | 354 | 323 | 395 |
| Chr15 | 1,426 | 468 | 346 | 276 | 336 |
| Chr16 | 1,520 | 457 | 419 | 342 | 302 |
| Gene with annotated alleles | 34,633 | 9,362 | 10,262 | 7,509 | 7,500 |
| Unanchored genes/alleles | 588 | - | - | - | - |

**Supplementary Table 9. Genes number of the three *C. paliurus* genomes.**

| Chromosome Number | PA-dip | PG-dip | PA-tetra | | | | |
|---|---|---|---|---|---|---|---|
| | | | Gene Number | No. of alleles in Hap A | No. of alleles in Hap B | No. of alleles in Hap C | No. of alleles in Hap D |
| Chr1 | 2,814 | 2,834 | 2,777 | 1,771 | 1,674 | 1,761 | 1,890 |
| Chr2 | 2,916 | 2,914 | 2,851 | 1,862 | 1,845 | 1,780 | 1,877 |
| Chr3 | 2,019 | 2,016 | 2,035 | 1,392 | 1,300 | 1,293 | 1,261 |
| Chr4 | 2,242 | 2,308 | 2,349 | 1,694 | 1,636 | 1,501 | 1,452 |
| Chr5 | 2,538 | 2,575 | 2,598 | 1,681 | 1,619 | 1,731 | 1,679 |
| Chr6 | 2,162 | 2,230 | 2,277 | 1,613 | 1,621 | 1,475 | 1,327 |
| Chr7 | 2,366 | 2,440 | 2,364 | 1,768 | 1,650 | 1,630 | 1,700 |
| Chr8 | 1,641 | 1,692 | 2,229 | 1,454 | 1,358 | 1,138 | 1,128 |
| Chr9 | 1,954 | 2,058 | 2,093 | 1,418 | 1,336 | 1,408 | 1,490 |
| Chr10 | 2,104 | 2,154 | 2,210 | 1,528 | 1,621 | 1,512 | 1,489 |
| Chr11 | 1,902 | 1,824 | 2,405 | 1,425 | 1,440 | 1,399 | 1,634 |
| Chr12 | 1,790 | 1,824 | 1,912 | 1,409 | 1,170 | 1,178 | 1,347 |
| Chr13 | 1,858 | 1,843 | 1,924 | 1,268 | 1,335 | 1,337 | 1,111 |
| Chr14 | 1,522 | 1,601 | 1,663 | 1,142 | 1,114 | 1,142 | 1,069 |
| Chr15 | 1,321 | 1,274 | 1,426 | 950 | 952 | 969 | 927 |
| Chr16 | 1,522 | 1,499 | 1,520 | 1,091 | 1,035 | 1,020 | 925 |
| Anchored | 32,671 | 33,086 | 34,633 | - | - | - | - |
| Total Number | 34,699 | 35,221 | 90,752 | 23,466 | 22,706 | 22,274 | 22,306 |

**Supplementary Table 10. BUSCO analysis of annotation completeness.**

| Description | PA-dip | | PG-dip | | PA-tetra | |
|---|---|---|---|---|---|---|
| | Number | Percentage (%) | Number | Percentage (%) | Number | Percentage (%) |
| Complete BUSCOs(C) | 1,323 | 96.2 | 1,323 | 96.2 | 1,299 | 94.4 |
| Complete and single-copy BUSCOs(S) | 1,203 | 87.5 | 1,200 | 87.3 | 178 | 12.9 |
| Complete and duplicated BUSCOs(D) | 120 | 8.7 | 123 | 8.9 | 1,121 | 81.5 |
| Fragmented BUSCOs(F) | 25 | 1.8 | 30 | 2.2 | 18 | 1.3 |
| Missing BUSCOs(M) | 27 | 2 | 22 | 1.6 | 58 | 4.3 |
| Total BUSCO groups searched | 1,375 | 100 | 1,375 | 100 | 1,375 | 100 |

**Supplementary Table 11. Repeat annotation of the three *C. paliurus* genomes.**

| | Repeat Number | | | Repeat Length (Mb) | | | Percent of repeat sequences (%) | | | Percent of assembled genome (%) | | |
|---|---|---|---|---|---|---|---|---|---|---|---|---|
| | PA-dip | PG-dip | PA-tetra | PA-dip | PG-dip | PA-tetra | PA-dip | PG-dip | PA-tetra | PA-dip | PG-dip | PA-tetra |
| Total repeat fraction | 955,148 | 965,878 | 3,979,983 | 282.25 | 316.95 | 1,154.35 | 100.00 | 100.00 | 100.00 | 48.10 | 54.32 | 48.41 |
| Class I: Retroelement | 456,693 | 468,102 | 1,863,889 | 211.85 | 217.88 | 851.72 | 75.06 | 68.74 | 73.78 | 36.10 | 37.34 | 35.72 |
| LTR Retrotransposon | 198,738 | 209,954 | 772,133 | 115.38 | 119.20 | 451.36 | 40.88 | 37.61 | 39.10 | 19.66 | 20.43 | 18.93 |
| Ty1/Copia | 48,976 | 54,939 | 199,474 | 31.46 | 34.29 | 123.30 | 11.15 | 10.82 | 10.68 | 5.36 | 5.88 | 5.17 |
| Ty3/Gypsy | 53,278 | 50,928 | 187,167 | 36.63 | 37.25 | 124.15 | 12.98 | 11.75 | 10.76 | 6.24 | 6.38 | 5.21 |
| Other | 96,484 | 104,087 | 385,492 | 47.29 | 47.67 | 203.91 | 16.75 | 15.04 | 17.66 | 8.06 | 8.17 | 8.55 |
| non-LTR Retrotransposon | 169,512 | 169,358 | 719,074 | 76.21 | 77.89 | 312.45 | 27.00 | 24.57 | 27.07 | 12.99 | 13.35 | 13.10 |
| LINE | 137,465 | 137,330 | 567,561 | 71.87 | 73.46 | 289.88 | 25.46 | 23.18 | 25.11 | 12.25 | 12.59 | 12.16 |
| SINE | 32,047 | 32,028 | 151,513 | 4.34 | 4.43 | 22.57 | 1.54 | 1.40 | 1.96 | 0.74 | 0.76 | 0.95 |
| Unclassified retroelement | 88,443 | 88,790 | 372,682 | 20.26 | 20.79 | 87.91 | 7.18 | 6.56 | 7.62 | 3.45 | 3.56 | 3.69 |
| Class II: DNA Transposon | 200,854 | 204,885 | 843,928 | 61.25 | 64.39 | 257.05 | 21.70 | 20.32 | 22.27 | 10.44 | 11.03 | 10.78 |
| CMC | 13,622 | 10,427 | 53,767 | 10.36 | 8.76 | 36.43 | 3.67 | 2.76 | 3.16 | 1.76 | 1.50 | 1.53 |

| | | | | | | | | | | | | |
|---|---|---|---|---|---|---|---|---|---|---|---|---|
| hAT | 29,910 | 34,166 | 122,713 | 11.42 | 13.28 | 45.15 | 4.05 | 4.19 | 3.91 | 1.95 | 2.28 | 1.89 |
| Mutator | 5,082 | 2,708 | 14,995 | 2.47 | 2.07 | 8.79 | 0.87 | 0.65 | 0.76 | 0.42 | 0.35 | 0.37 |
| PIF/Harbinger | 6,305 | 4,268 | 26,036 | 2.67 | 1.61 | 12.11 | 0.94 | 0.51 | 1.05 | 0.45 | 0.28 | 0.51 |
| Other | 145,935 | 153,316 | 626,417 | 34.34 | 38.67 | 154.57 | 12.16 | 12.20 | 13.39 | 5.85 | 6.63 | 6.48 |
| Helitron | 3,372 | 1,750 | 7,842 | 1.59 | 0.89 | 4.22 | 0.56 | 0.28 | 0.37 | 0.27 | 0.15 | 0.18 |
| Tandem Repeats | 206,563 | 202,418 | 889,607 | 13.87 | 12.90 | 72.59 | 4.91 | 4.07 | 6.29 | 2.36 | 2.21 | 3.04 |
| Unkown | 34,628 | 35,897 | 143,126 | 9.40 | 10.11 | 40.05 | 3.33 | 3.19 | 3.47 | 1.60 | 1.73 | 1.68 |

**Supplementary Table 12. The expanded P450s families in the *C. paliurus* genome.**

| Gene family | Number of genes | Gene ID |
|---|---|---|
| CYP71 | 1 | CpaM1st11714 |
| CYP82 | 6 | CpaM1st11392 |
| | | CpaM1st11409 |
| | | CpaM1st11411 |
| | | CpaM1st11418 |
| | | CpaM1st25816 |
| | | CpaM1st25822 |
| CYP89 | 16 | CpaM1st00845 |
| | | CpaM1st00847 |
| | | CpaM1st00850 |
| | | CpaM1st00853 |
| | | CpaM1st00858 |
| | | CpaM1st00863 |
| | | CpaM1st00864 |
| | | CpaM1st00866 |
| | | CpaM1st00872 |
| | | CpaM1st00873 |
| | | CpaM1st00874 |
| | | CpaM1st00881 |
| | | CpaM1st00885 |
| | | CpaM1st14090 |
| | | CpaM1st14091 |
| | | CpaM1st35816 |
| CYP706 | 18 | CpaM1st06185 |
| | | CpaM1st09353 |
| | | CpaM1st17603 |
| | | CpaM1st17613 |
| | | CpaM1st17625 |
| | | CpaM1st17635 |
| | | CpaM1st20465 |
| | | CpaM1st26335 |
| | | CpaM1st26346 |
| | | CpaM1st26520 |
| | | CpaM1st26526 |
| | | CpaM1st28507 |
| | | CpaM1st28509 |
| | | CpaM1st28521 |
| | | CpaM1st28524 |
| | | CpaM1st28527 |
| | | CpaM1st28613 |
| | | CpaM1st28615 |
| CYP727 | 1 | CpaM1st30617 |

**Supplementary Table 13. Comparison of growth indices between diploid and tetraploid *C. paliurus* seedling, one year after stumping**

| Determination standard | Ploidy | Extremum | Mean±SD | Character difference | Coefficient of variation |
|---|---|---|---|---|---|
| Seedling height/cm | Diploid | 57.00-152.00 | 105.98±19.38 | 11.38 | 0.22 |
| | Tetraploid | 80.00-172.40 | 117.36±17.51* | | 0.15 |
| Ground diameter/mm | Diploid | 4.26-17.30 | 11.16±2.49 | 6.3 | 0.22 |
| | Tetraploid | 6.94-27.27 | 17.46±4.37** | | 0.25 |
| Compound leaf fresh weight/g | Diploid | 0.43-5.26 | 2.53±1.01 | 1.74 | 0.40 |
| | Tetraploid | 1.29-7.76 | 4.27±1.58** | | 0.37 |
| Compound leaf dry weight/g | Diploid | 0.14-1.96 | 0.77±0.35 | 0.72 | 0.45 |
| | Tetraploid | 0.39-3.22 | 1.49±0.64** | | 0.43 |
| Leaf moisture content | Diploid | 0.24-0.44 | 0.30±0.03 | 0.05 | 0.10 |
| | Tetraploid | 0.25-0.68 | 0.35±0.08 | | 0.23 |
| Compound leaf area/$cm^2$ | Diploid | 43.31-503.39 | 253.10±93.79 | 136.29 | 0.37 |
| | Tetraploid | 86.39-636.79 | 389.39±138.52** | | 0.36 |
| Leaf specific weight/($g/cm^2$) | Diploid | 0.0016-0.0076 | 0.0031±0.0010 | 0.0008 | 0.33 |
| | Tetraploid | 0.0027-0.0056 | 0.0039±0.0008 | | 0.21 |
| Number of saw teeth | Diploid | 24-94 | 61±12 | 3 | 0.20 |
| | Tetraploid | 48-97 | 64±13 | | 0.20 |
| Blade length/cm | Diploid | 2.95-19.10 | 9.71±2.25 | 3.06 | 0.23 |
| | Tetraploid | 4.74-16.50 | 12.77±2.86** | | 0.22 |
| Blade width/cm | Diploid | 1.47-5.50 | 2.96±0.78 | 1.77 | 0.26 |
| | Tetraploid | 2.74-6.70 | 4.73±1.09** | | 0.23 |
| Blade aspect ratio | Diploid | 1.19-4.51 | 3.34±0.50** | -0.6 | 0.15 |
| | Tetraploid | 1.46-3.64 | 2.74±0.47 | | 0.17 |

Note: * * and * indicate significant differences at 0.01 and 0.05 levels, respectively. Character difference = Mean of tetraploid - diploid.

**Supplementary Table 14.** ***C. paliurus*** **and outgroup (*Juglans regia*) species used for the population genomics analysis**

| Position | No of individuals | Country | Location | ID | Altitude(m) | Latitude (° E) | Longitude (° N) | Ploidy | X-mean | Reference | Coverage (×) |
|---|---|---|---|---|---|---|---|---|---|---|---|
| A | 5 | China | Hefeng County, Hubei | A1 | 1516 | 110.456345 | 29.955051 | Diploid | 10,169 | 19,009 | 21.83 |
| | | | | A2 | 1340 | 110.460312 | 29.971876 | Diploid | 10,171 | 19,270 | 10.33 |
| | | | | A3 | 1494 | 110.462555 | 29.965052 | Diploid | 10,139 | 18,670 | 11.00 |
| | | | | A4 | 970 | 110.433548 | 29.855425 | Auto-tetraploid | 20,779 | 11,417 | 15.67 |
| | | | | A5 | 1340 | 110.460060 | 29.971657 | Diploid | 10,945 | 19,855 | 11.00 |
| B | 5 | China | Houhe National Nature Reserve, Hubei | B1 | 1600 | 110.543877 | 30.069838 | Diploid | 9,890 | 18,292 | 10.33 |
| | | | | B2 | 1629 | 110.544258 | 30.064539 | Diploid | 11,139 | 19,871 | 11.00 |
| | | | | B3 | 1650 | 110.567062 | 30.066883 | Diploid | 10,413 | 18,835 | 10.83 |
| | | | | B4 | 1630 | 110.600578 | 30.111212 | Diploid | 10,726 | 19,360 | 10.83 |
| | | | | B5 | 1400 | 110.541718 | 30.085014 | Auto-tetraploid | 21,019 | 11,422 | 10.50 |
| C | 5 | China | Huangshan, Anhui | C1 | 711 | 118.165787 | 30.104895 | Auto4-tetraploid | 18,751 | 10,308 | 10.58 |
| | | | | C2 | 716 | 118.166092 | 30.105141 | Auto-tetraploid | 19,334 | 10,037 | 15.50 |
| | | | | C3 | 630 | 118.171951 | 30.100531 | Auto-tetraploid | 19,285 | 10,349 | 10.42 |

| | | | | | | | | | | | |
|---|---|---|---|---|---|---|---|---|---|---|---|
| | | | | C4 | 967 | 118.169220 | 30.159826 | Auto-tetraploid | 19,132 | 10,562 | 15.50 |
| | | | | C5 | 1316 | 118.167465 | 30.152409 | Diploid | 9,865 | 17,568 | 21.00 |
| D | 5 | China | Qingliangfeng Nature Reserve, Anhui | D1 | 680 | 118.893478 | 30.149855 | Auto-tetraploid | 19,005 | 10,207 | 15.33 |
| | | | | D2 | 907 | 118.893867 | 30.139042 | Auto-tetraploid | 18,595 | 9,907 | 10.92 |
| | | | | D3 | 685 | 118.894279 | 30.149760 | Auto-tetraploid | 20,008 | 10,466 | 15.17 |
| | | | | D4 | 674 | 118.856438 | 30.151596 | Auto-tetraploid | 18,590 | 10,122 | 15.17 |
| | | | | D5 | 1406 | 118.842934 | 30.127003 | Diploid | 8,920 | 15,833 | 10.33 |
| E | 5 | China | Chiu-ling Mountains, Jiangxi | E1 | 771 | 115.324646 | 29.091516 | Auto-tetraploid | 17,192 | 8,858 | 19.92 |
| | | | | E2 | 780 | 115.264275 | 29.015097 | Auto-tetraploid | 18,105 | 9,951 | 10.58 |
| | | | | E3 | 771 | 110.442207 | 29.965858 | Auto-tetraploid | 19,122 | 10,165 | 10.00 |
| | | | | E4 | 577 | 115.222290 | 29.029156 | Auto-tetraploid | 19,629 | 10,867 | 10.17 |
| | | | | E5 | 580 | 115.223457 | 29.033163 | Auto-tetraploid | 19,819 | 11,574 | 11.08 |
| F | 5 | China | Tonggu County, Jiangxi | F1 | 875 | 114.219116 | 28.519285 | Auto-tetraploid | 19,905 | 11,494 | 11.25 |
| | | | | F2 | 781 | 114.206985 | 28.530428 | Auto-tetraploid | 19,391 | 10,981 | 10.83 |
| | | | | F3 | 793 | 114.207760 | 28.539653 | Auto-tetraploid | 19,336 | 10,509 | 11.33 |
| | | | | F4 | 812 | 114.220161 | 28.516615 | Auto-tetraploid | 18,581 | 10,251 | 11.67 |
| | | | | F5 | 762 | 114.208870 | 28.530558 | Auto-tetraploid | 18,945 | 10,298 | 11.25 |
| G | 5 | China | Shucheng [illegible] | G1 | 798 | 116.540108 | 31.037395 | Auto-tetraploid | 20,020 | 11,302 | 11.33 |

| | | | | | | | | | | | |
|---|---|---|---|---|---|---|---|---|---|---|---|
| | | | | G2 | 831 | 116.541092 | 31.037144 | Auto-tetraploid | 18,987 | 10,369 | 10.83 |
| | | | | G3 | 744 | 116.539253 | 31.039656 | Auto-tetraploid | 20,421 | 11,645 | 10.75 |
| | | | | G4 | 764 | 116.539048 | 31.039019 | Auto-tetraploid | 19,074 | 10,917 | 11.42 |
| | | | | G5 | 766 | 116.539162 | 31.037907 | Auto-tetraploid | 19,636 | 10,223 | 10.50 |
| | | | | H1 | 1810 | 104.950639 | 24.617250 | Auto-tetraploid | 20,681 | 10,922 | 15.08 |
| | | | | H2 | 1829 | 104.952258 | 24.617875 | Auto-tetraploid | 21,125 | 11,951 | 11.00 |
| H | 5 | China | Jinzhongshan Nature Reserve, Guangxi | H3 | 1819 | 104.954131 | 24.617858 | Auto-tetraploid | 21,361 | 10,270 | 10.75 |
| | | | | H4 | 1698 | 104.953714 | 24.616278 | Auto-tetraploid | 20,620 | 10,870 | 11.00 |
| | | | | H5 | 1593 | 104.950094 | 24.612328 | Auto-tetraploid | 20,276 | 11,120 | 10.83 |
| | | | | I1 | 1217 | 103.791064 | 24.966956 | Auto-tetraploid | 20,636 | 10,749 | 19.92 |
| | | | | I2 | 1230 | 103.792557 | 28.965565 | Auto-tetraploid | 21,076 | 11,385 | 10.92 |
| I | 5 | China | Muchuan County, Sichuan | I3 | 1228 | 103.791656 | 28.965809 | Auto-tetraploid | 19,762 | 10,718 | 10.58 |
| | | | | I4 | 1192 | 103.793526 | 28.969936 | Auto-tetraploid | 18,226 | 9,970 | 10.75 |
| | | | | I5 | 1200 | 103.791069 | 28.966961 | Auto-tetraploid | 18,658 | 9,868 | 11.25 |
| Outgroup | 1 | China | Mei, haanxi, China | O | - | 107.75 | 33.82 | Diploid | - | - | 23.10 |

Note: X-mean represents the fluorescence intensity of the sample peak which is proportional to the DNA content of the cells. Reference mean of internal reference (diploid or tetraploid *C. paliurus*) for flow cytometry analysis. The data from Outgroup was referred to Zhang's report (Zhang et al. 2022).

**Supplementary Table 15. Summary of genetic variations in *C. paliurus* populations**

| Category | Diploid | Tetraploid |
|---|---|---|
| **Sequence variants** | | |
| SNPs | 3,545,162 | 23,076,276 |
| Indels | 341,670 | 3,598,719 |
| Variants with effects on genes | 38,874 | 282,037 |
| SNPs that introduce stop codons | 145 | 1,068 |
| SNPs that disrupt stop codons | 19 | 114 |
| SNPs that induce alternative splicing | 472 | 3,237 |
| Indels located in genic regions | 3,845 | 38,899 |
| Frameshift variants | 314 | 2,088 |
| Nonsynonymous variants | 5,503 | 35,241 |
| Synonymous variants | 3,753 | 23,764 |

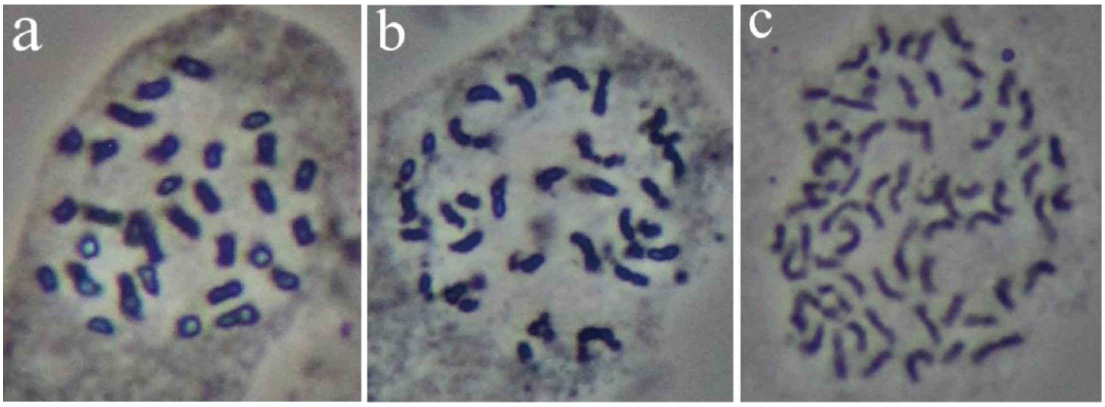


**Supplementary Figure 1. Chromosome karyotypes of *C. paliurus*.** (a) PA-dip. (b) PG-dip. (c) PA-tetra.

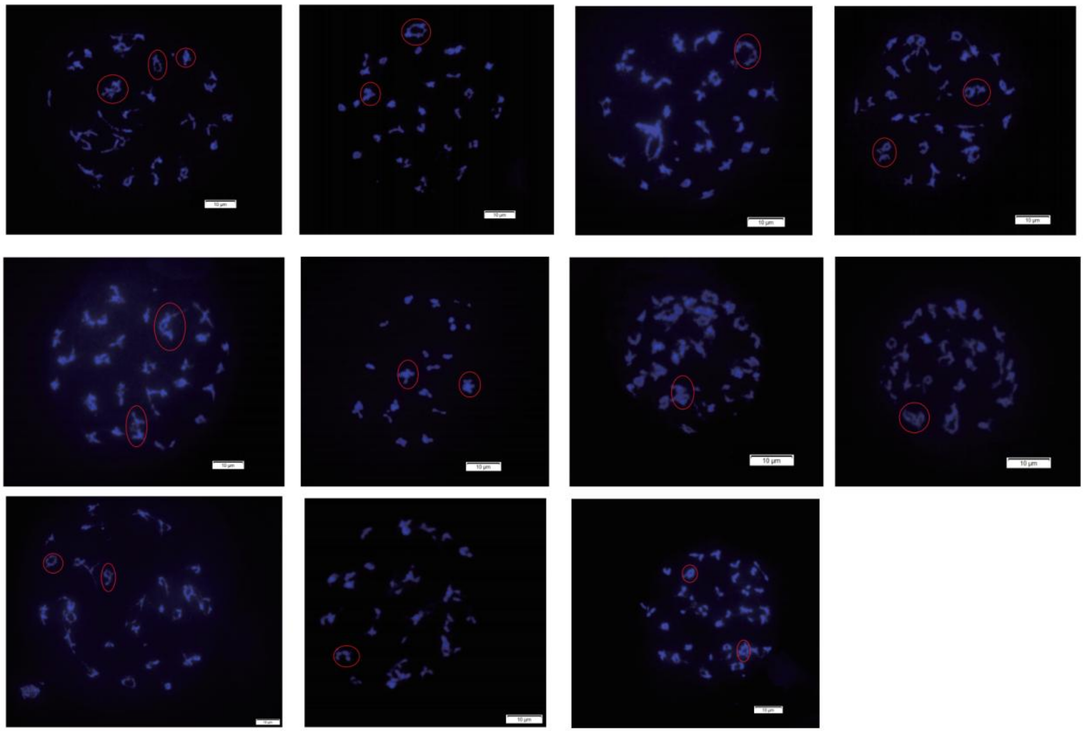

**Supplementary Figure 2. The phenomena of homologous chromosomes synapsis at the early stage of meiosis I in PA-tetra *C. paliurus*.** A total of 11 pollen mother cells were selected to observe and the red rounds represent the quadrivalent.

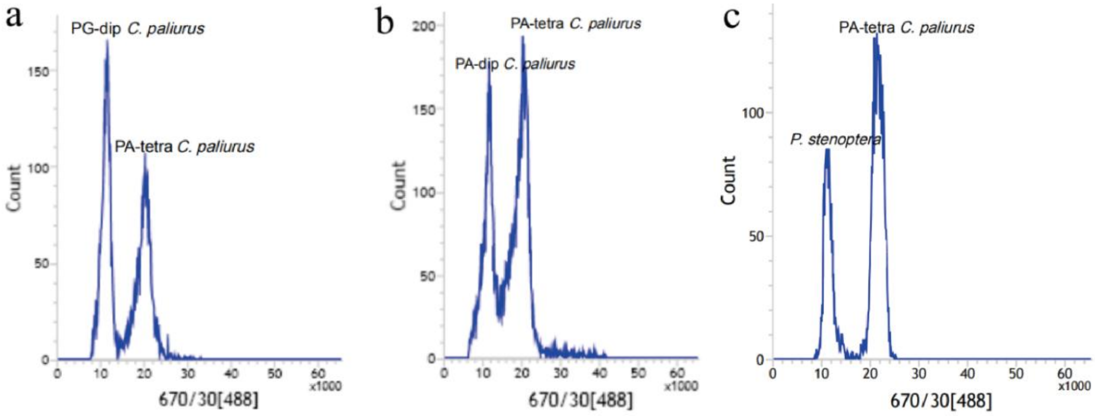


**Supplementary Figure 3. Using flow cytometry to estimate genome size of *C. paliurus*.** (a) PG-dip. (b) PA-dip. (c) PA-tetra. The *Pterocarya stenoptera* (*P. stenoptera*) genome (2n=2x=~600 Mb) and PA-tetra *C. paliurus* were used as an internal reference standard.

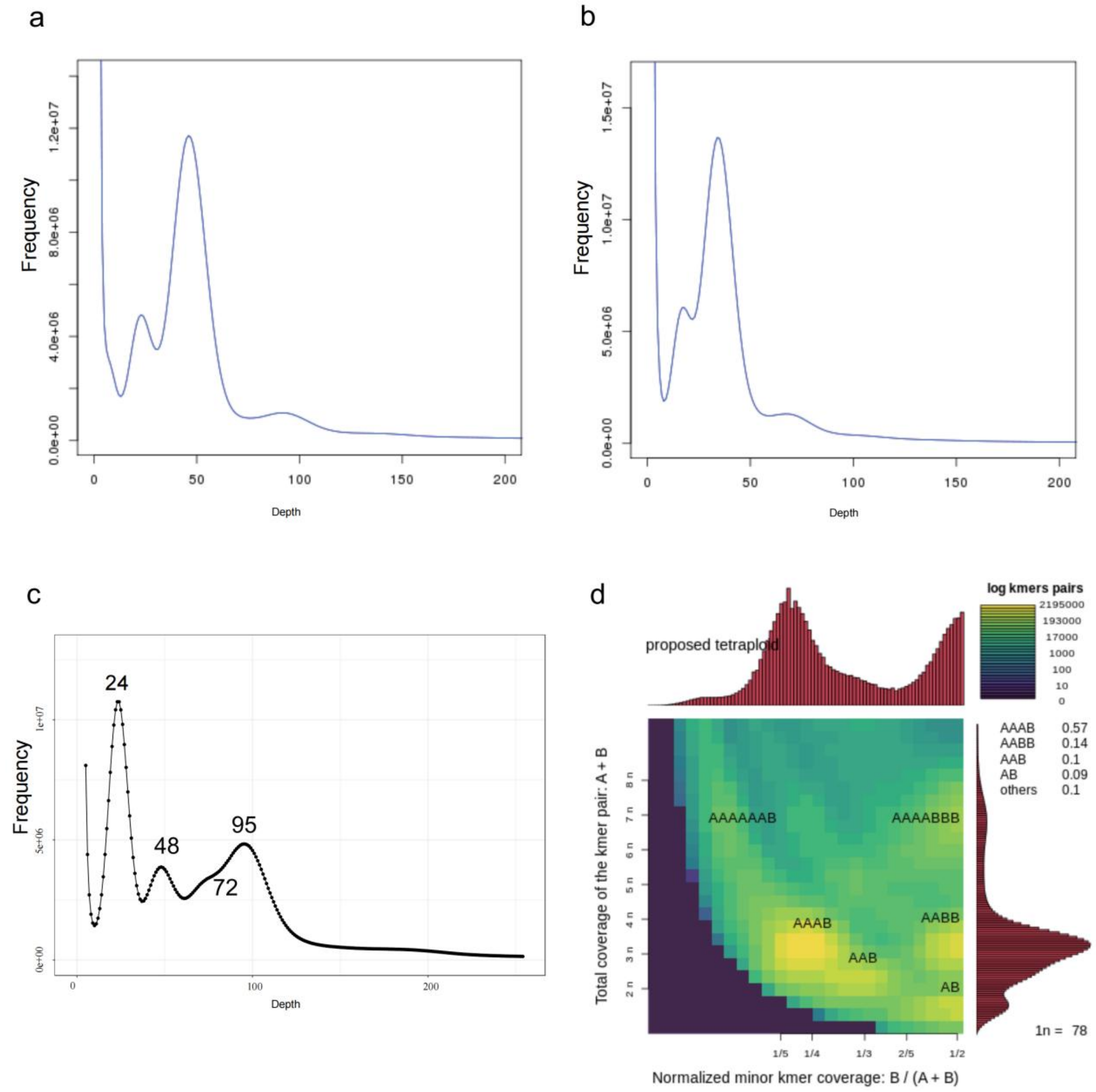


**Supplementary Figure 4. *K*-mer (21-mer) distribution and estimation of genome size and heterozygosity of *C. paliurus*.** (a)PA-dip. (b) PG-dip (b). (c) PA-tetra. (d). Total coverage of the *k*-mer pair (A+B) in PA-tetra *C. paliurus*.

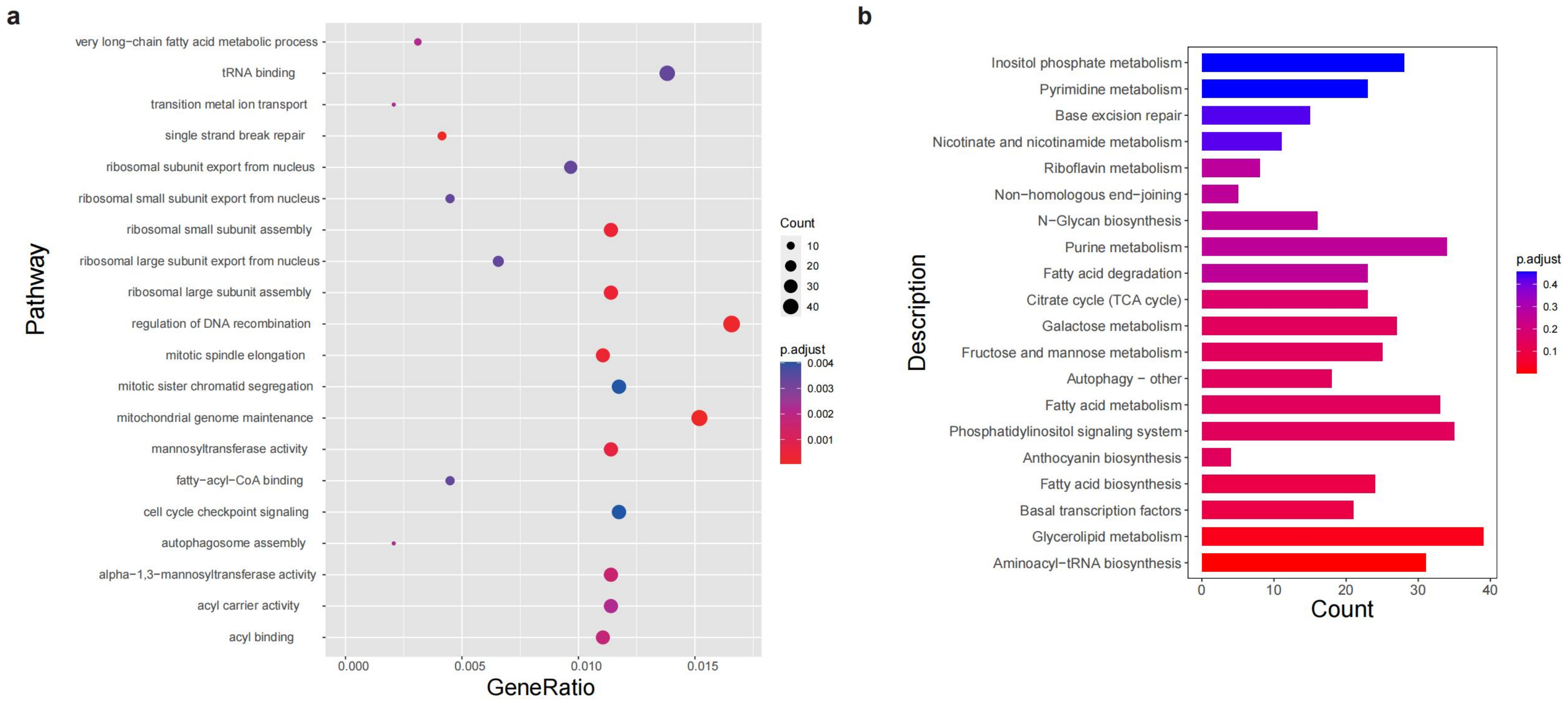


**Supplementary Figure 5. GO (a) and KEGG (b) enrichment analysis of selected genes involved in haplotypic variations in PA-tetra genome.**

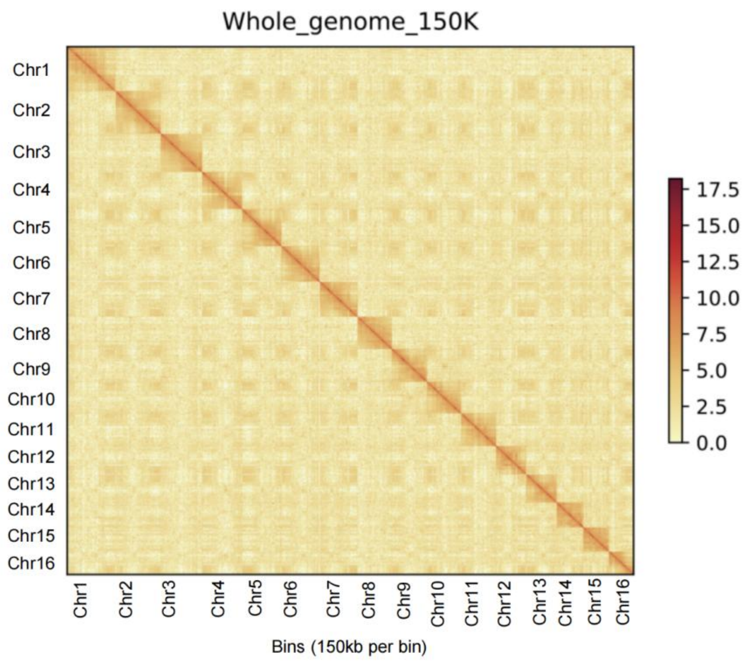


**Supplementary Figure 6. Genome-wide analysis of chromatin interactions at 150-kb resolution in PA-dip genome.** The colored bar on the right represents the strength of interaction.

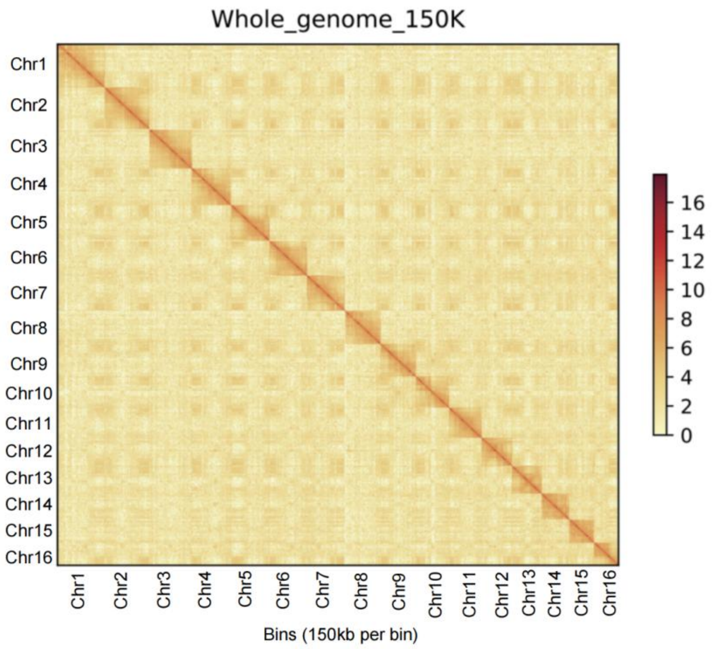


**Supplementary Figure 7. Genome-wide analysis of chromatin interactions at 150-kb resolution in PG-dip genome.**

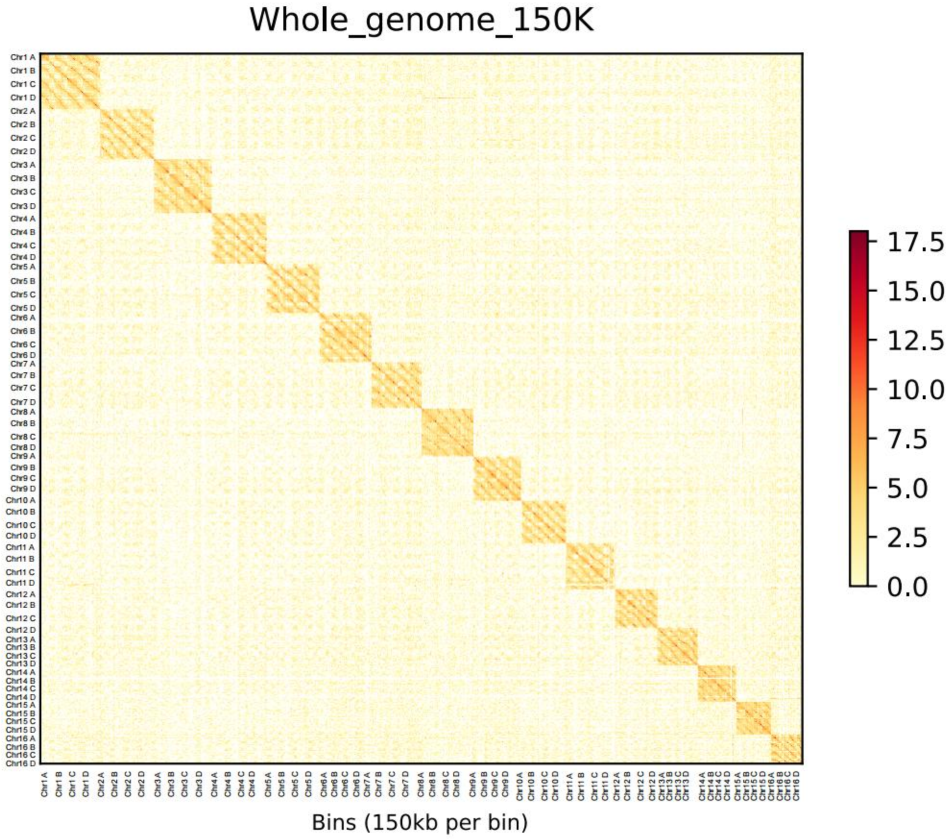


**Supplementary Figure 8. Genome-wide analysis of chromatin interactions at 150-kb resolution in PA-tetra genome.**

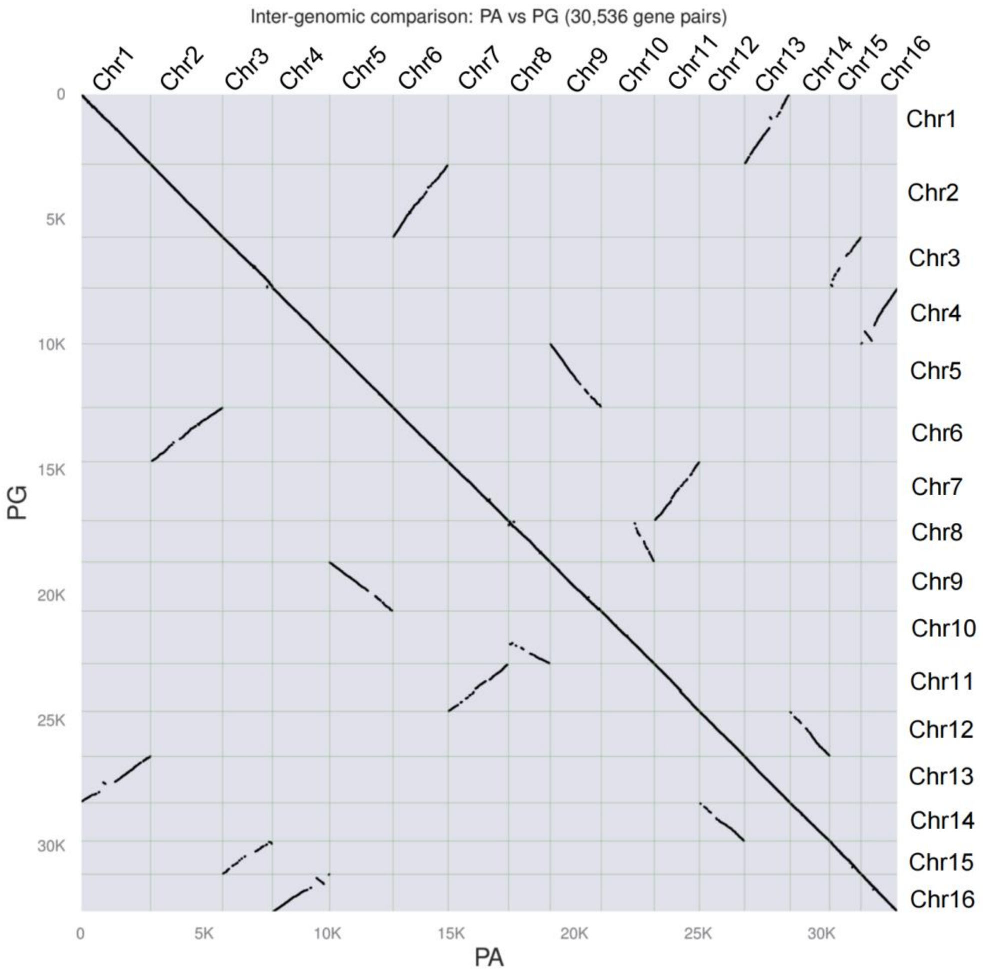


**Supplementary Figure 9. Synteny analysis between PA and PG diploid genomes.**

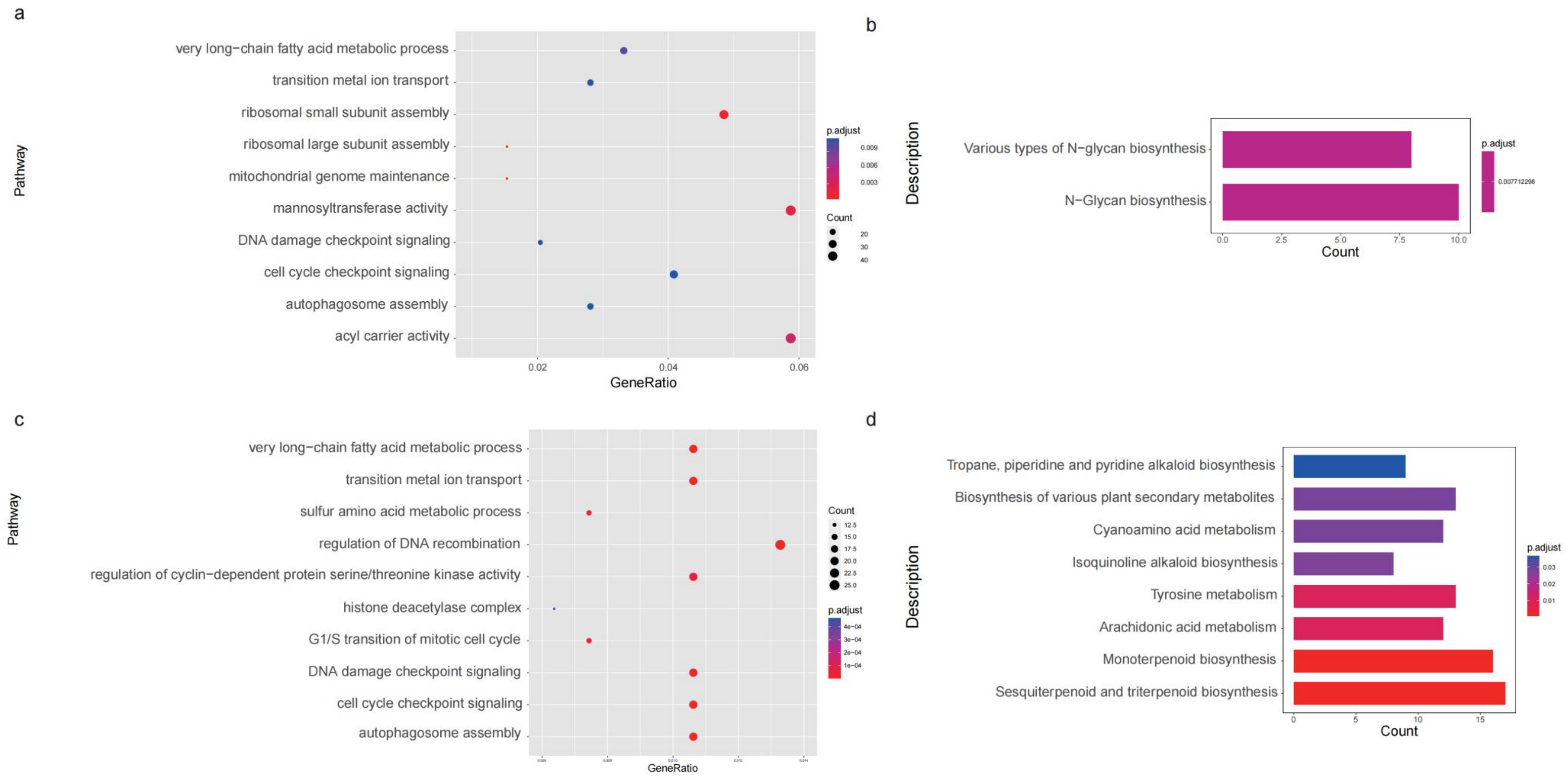


**Supplementary Figure 10. GO (a and c) and KEGG (b and d) enrichment analysis of selected genes involved in WGD1 (a and b) and WGD2 (c and d) events.**

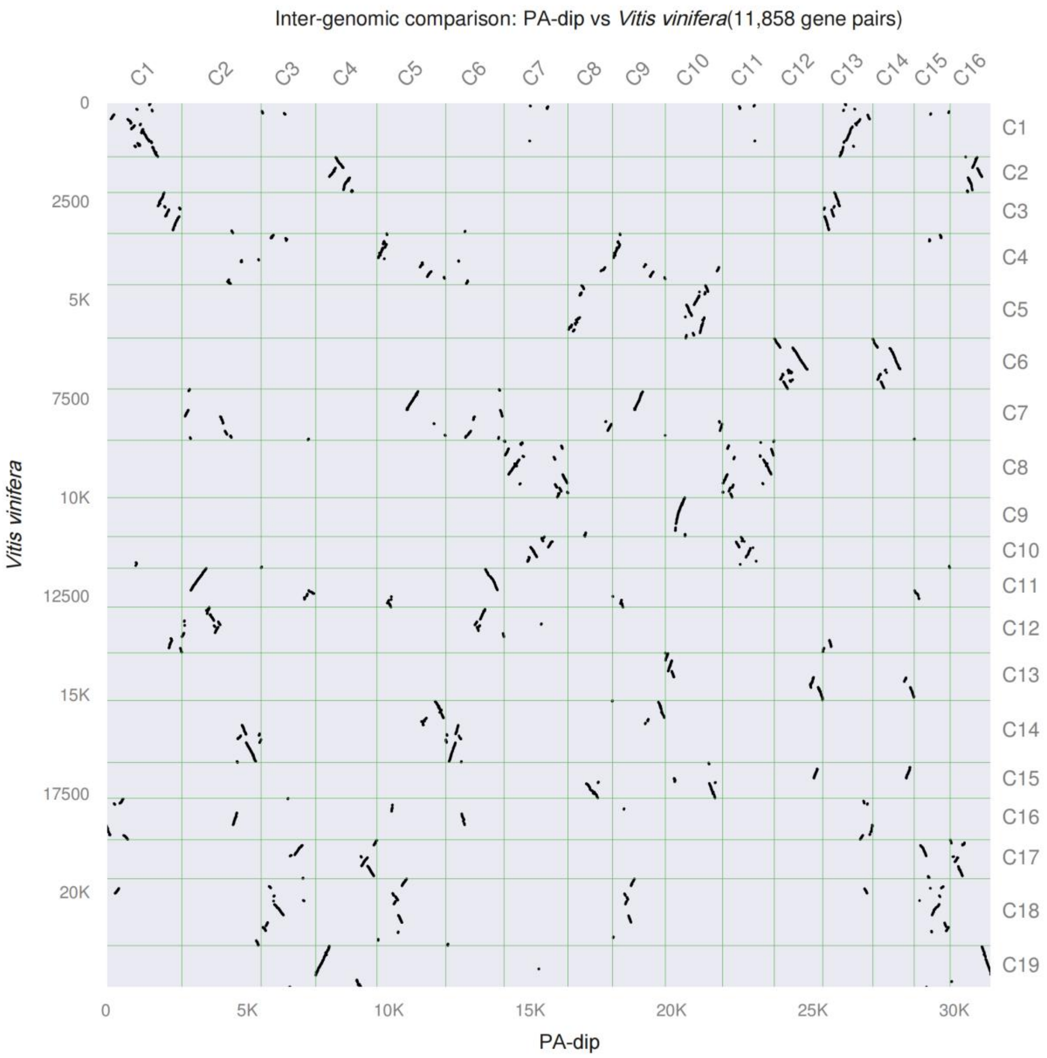


**Supplementary Figure 11. Synteny analysis between PA-dip and *Vitis vinifera* genomes.**

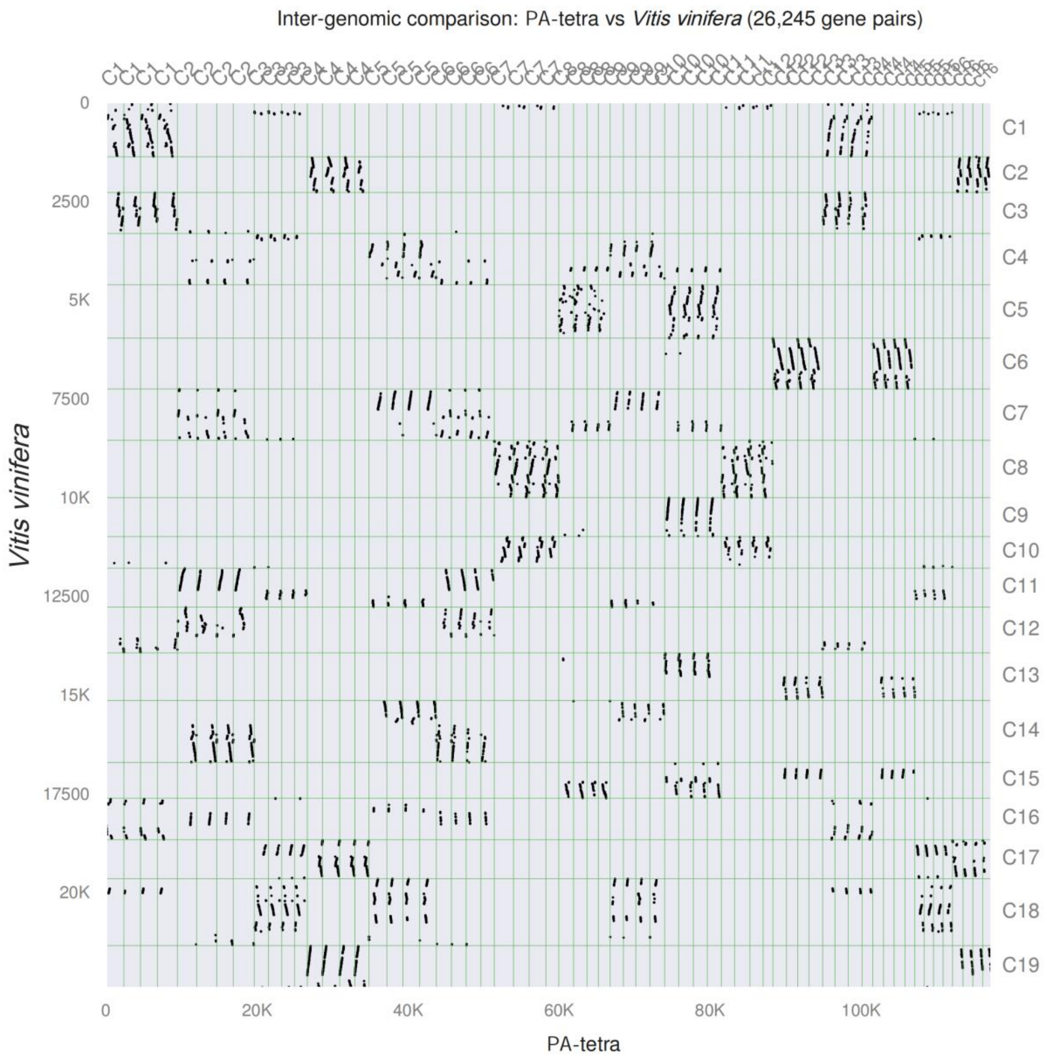


**Supplementary Figure 12. Synteny analysis between PA-tetra and *Vitis vinifera* genomes.**

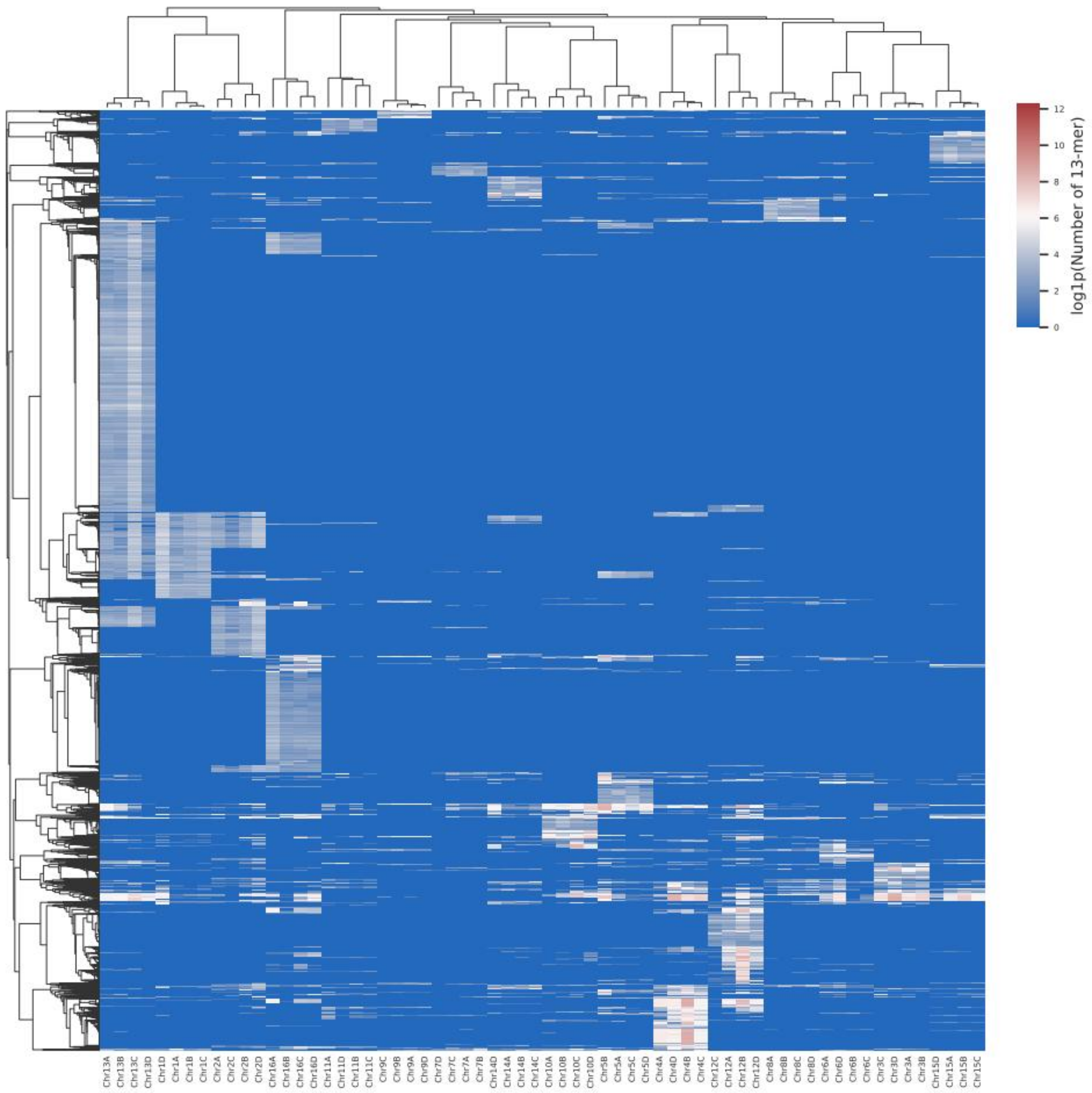


**Supplementary Figure 13. Clustering of counts of *K*-mers (*K*=13) enables the consistent partitioning of four haplotypes in each homologous chromosome into same group.**

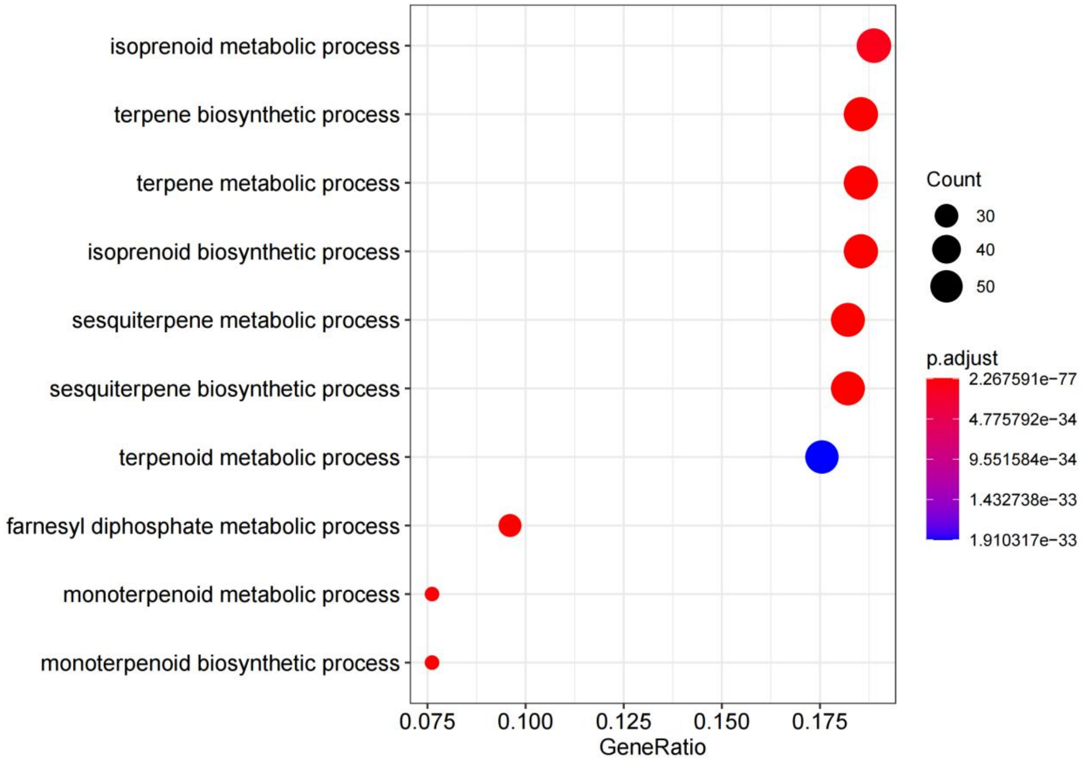


**Supplementary Figure 14. GO enrichment analysis of genes experienced expansion in *C. paliurus.***

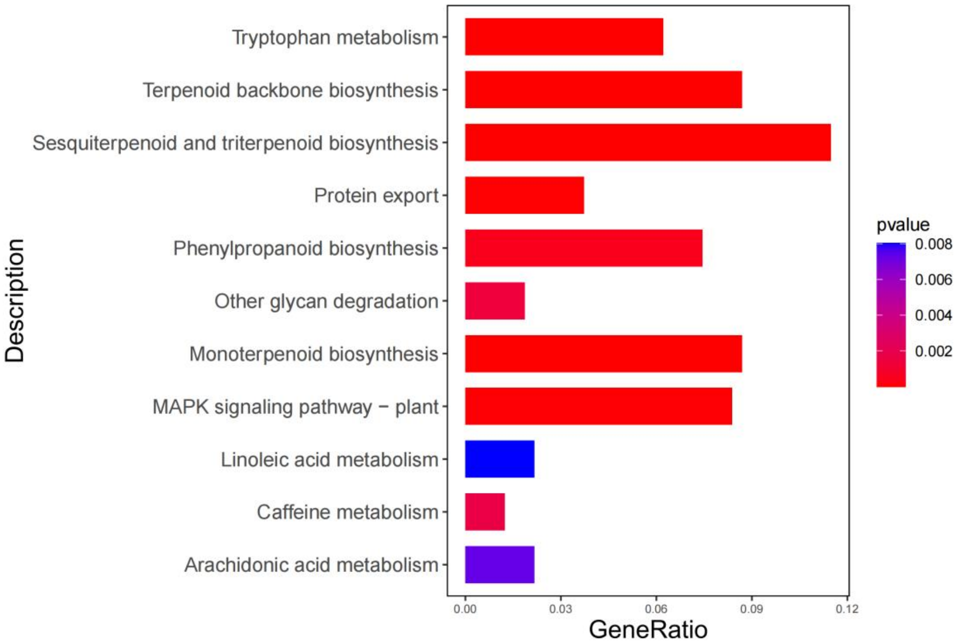


**Supplementary Figure 15. KEGG pathway analysis of genes experienced expansion in *C. paliurus.***

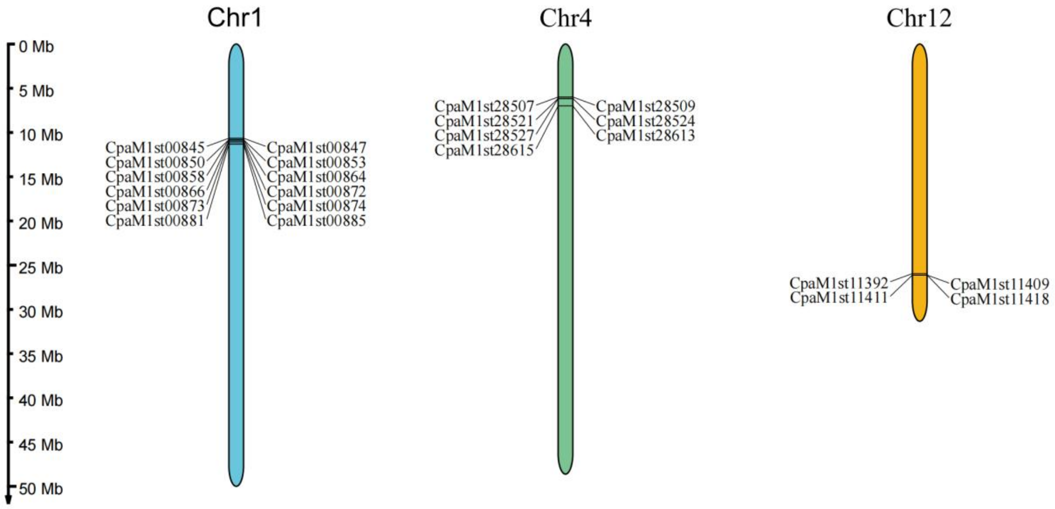


**Supplementary Figure 16. The P450s clusters of expanded genes are arranged on Chr1, Chr4, and Chr12.**

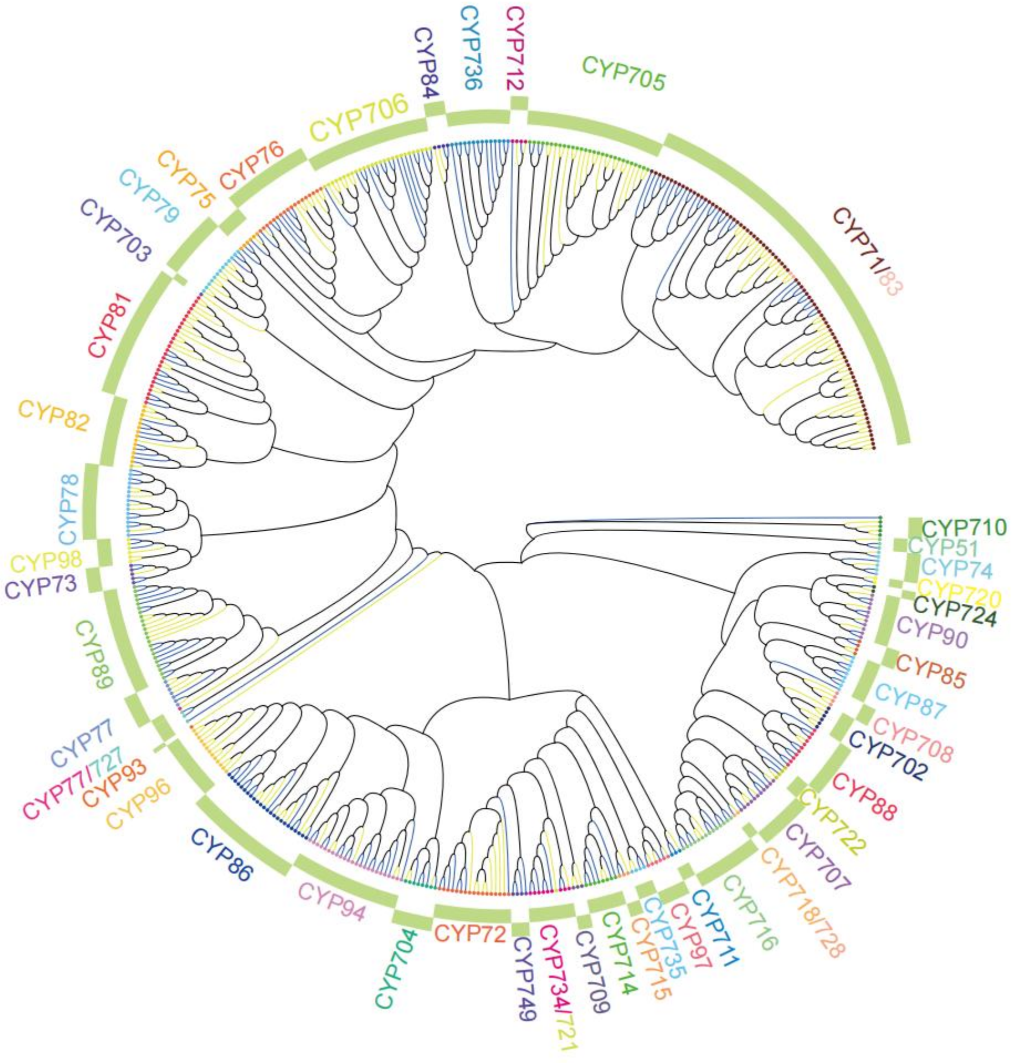


**Supplementary Figure 17. Phylogenic analysis of P450 families. The yellow and blue branches indicate the sequences from *Arabidopsis* and *C. paliurus*, respectively. The dots represent P450 genes. The outermost arc indicates the P450 gene family.**

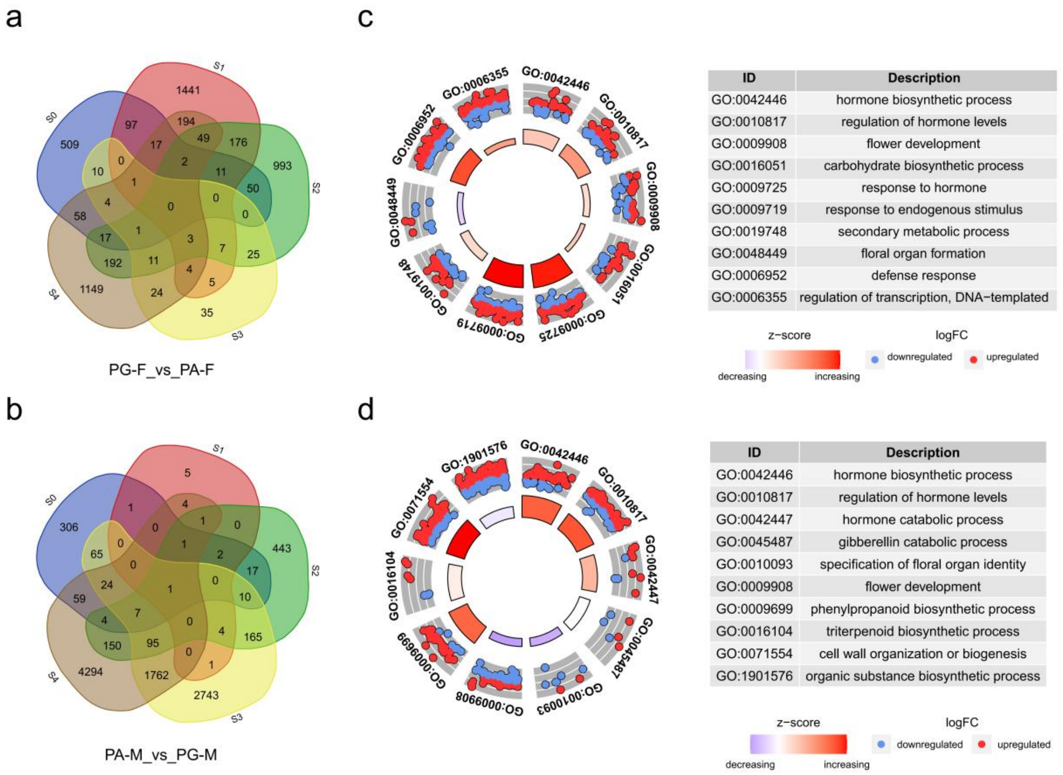


**Supplementary Figure 18. Identification and functional enrichment of DEGs between PG and PA *C. paliurus*.** Veen diagrams of DEGs in the female floral buds (a) and male floral buds (b) between PG-tetra and PA-tetra. In all subsequent figures, PG-F means female floral buds of PG, PA-F means female floral buds of PA, PG-M means male floral buds of PG, PA-M means male floral buds of PA.

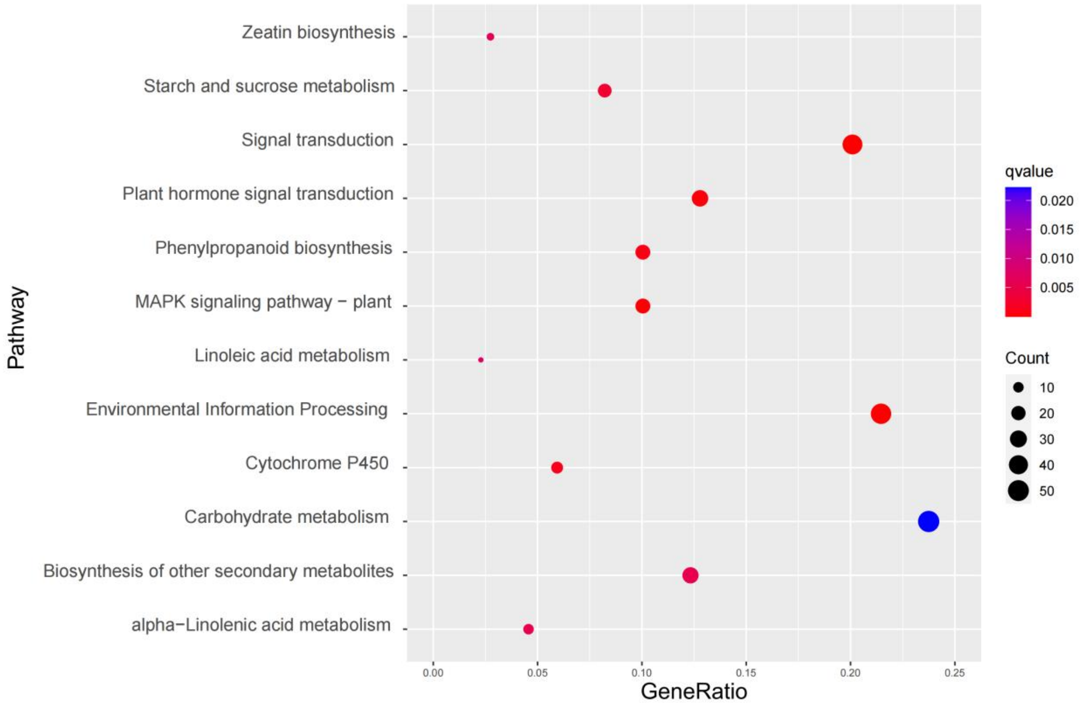


**Supplementary Figure 19. KEGG enrichment of the 958 differently expressed genes in female floral buds (PG-F_vs_PA-F) at five flowering time stages (S0-S4).**

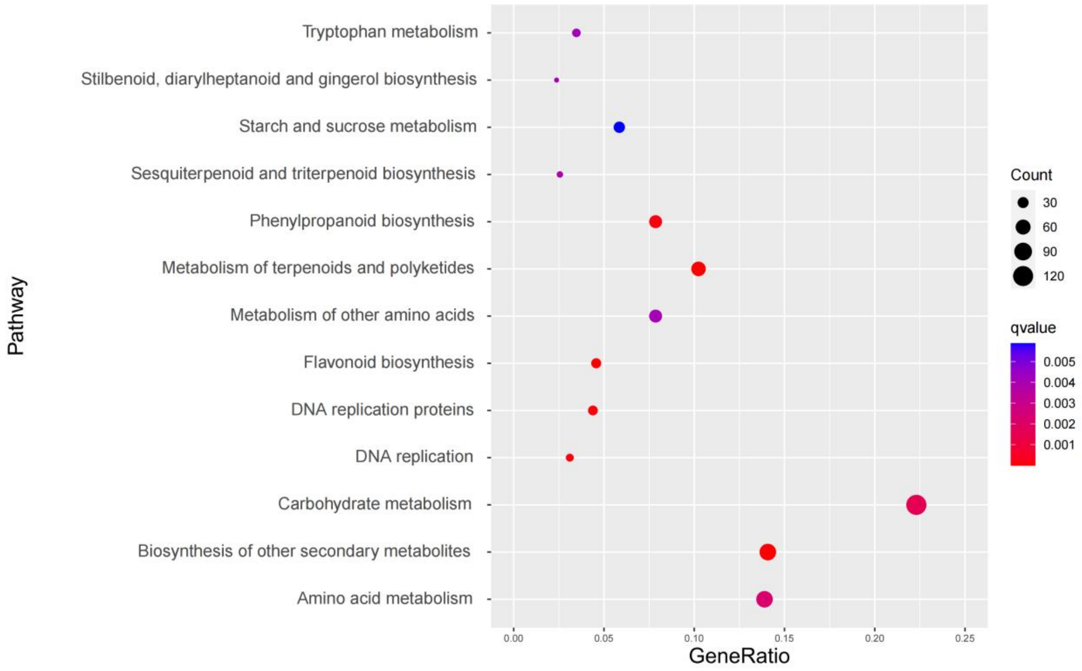


**Supplementary Figure 20. KEGG enrichment of the 2373 differently expressed genes in male floral buds (PA-M_vs_PG-M) at five flowering time stages (S0-S4).**

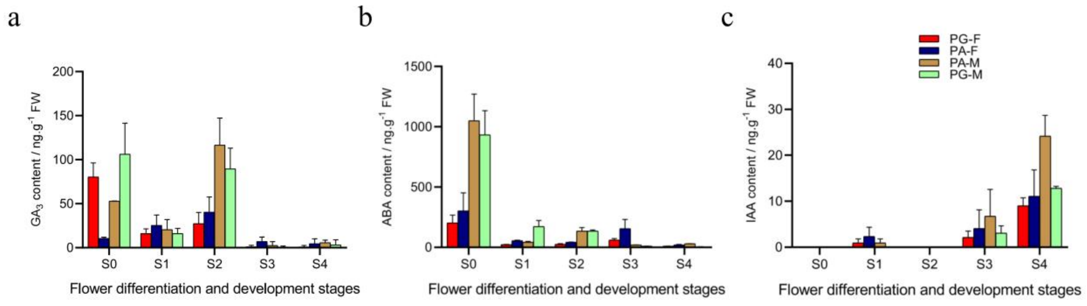


**Supplementary Figure 21. Hormones contents of *C. paliurus* during five flower differentiation and development stages.** (a) Identification of (gibberellin) $GA_3$ level in female and male floral buds from PG or PA type. (b) Identification of abscisic acid (ABA) level. (c) Identification of auxin (IAA) level.

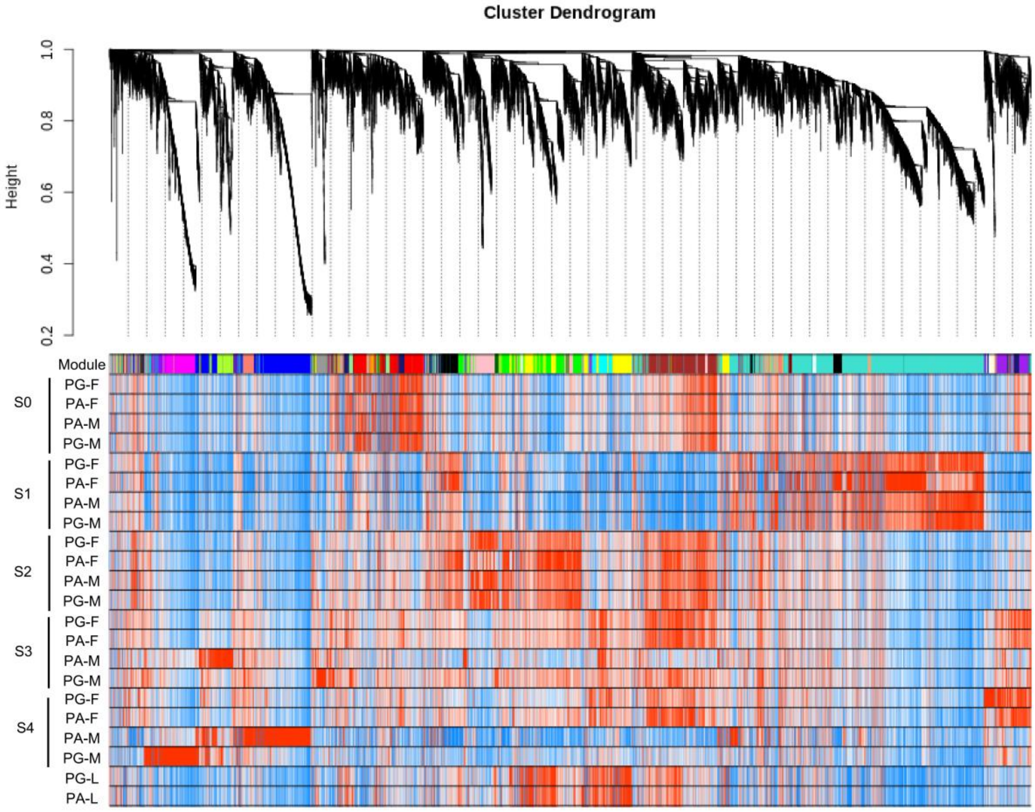


**Supplementary Figure 22. Hierarchical clustering tree (dendrogram) of genes based on coexpression network analysis in PG and PA individuals (female, male flora buds, and leaves) during five development stages.** Each individual's value is the average of the expression of three replicate samples. PG-L means leaves of PG, PA-L means leaves of PA.

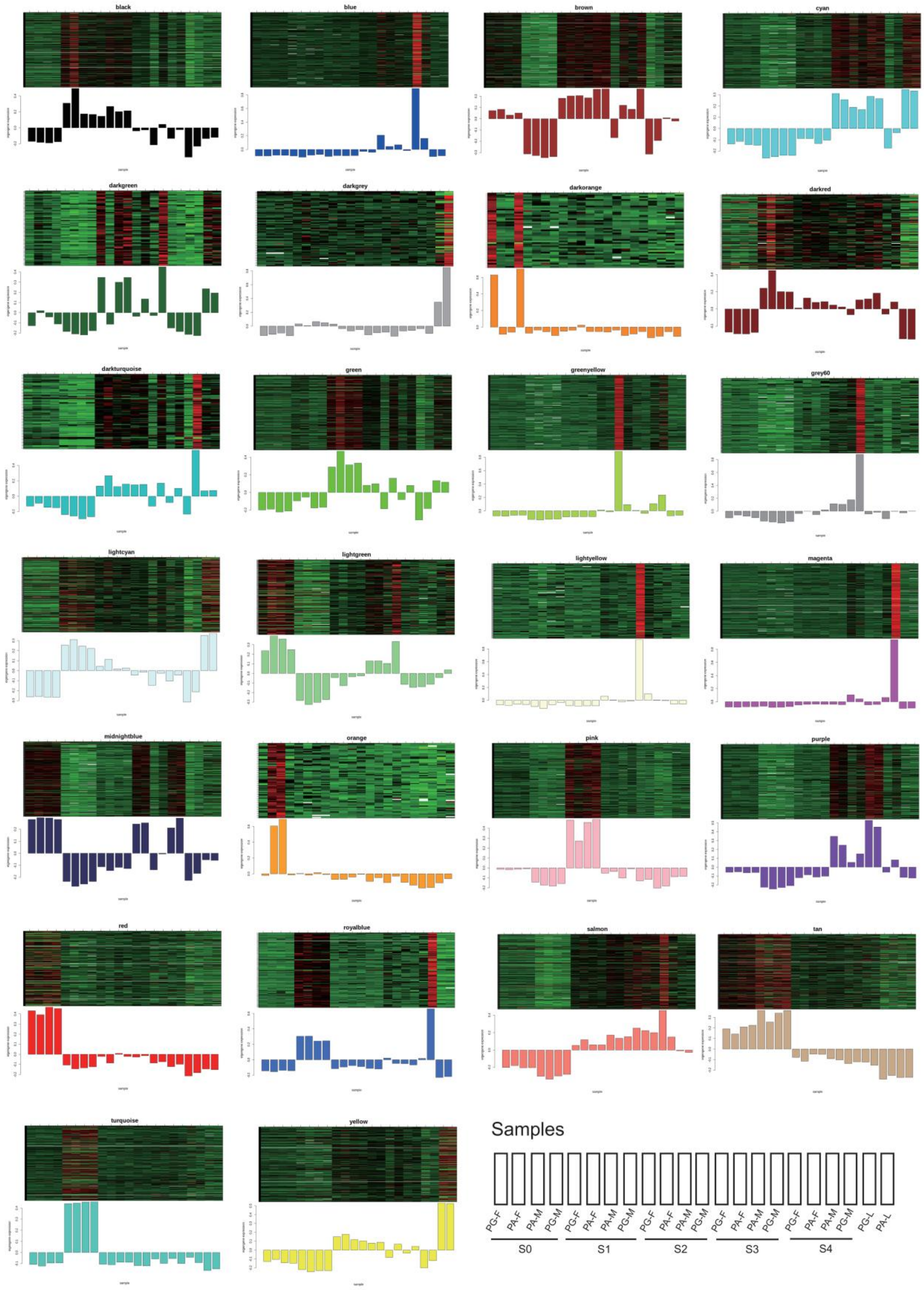


**Supplementary Figure 23. Co-expression modules identified by WGCNA.**

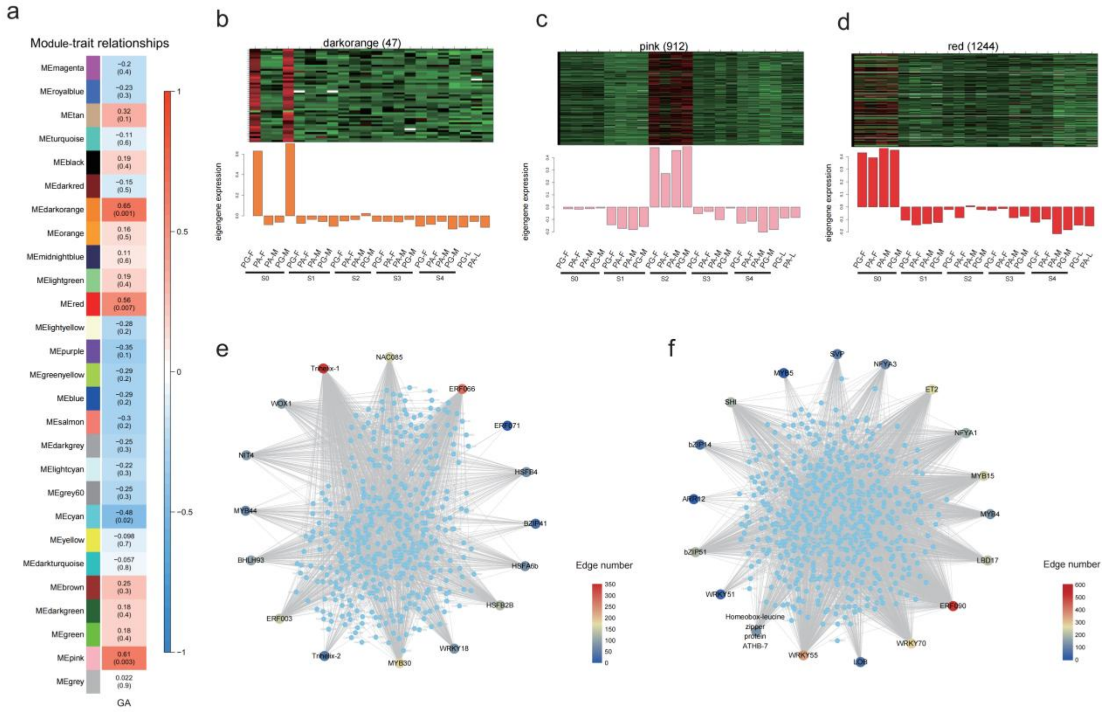

**Supplementary Figure 24. Functions and networks of coexpression module genes. (**a) Module-trait relationships. The column indicate GA, and the rows indicate the different modules. The red and blue colors indicate positive and negative correlations, respectively. The correlation coefficient (r) and P-value (p) are displayed in each cell. (b, c, d) Eigengene expression profiles in the darkorange, pink, and red modules. (e, f) Construction of the correlation network of the pink and red modules. The large circles represent transcription factors. The color is determined by the edge number of the gene.

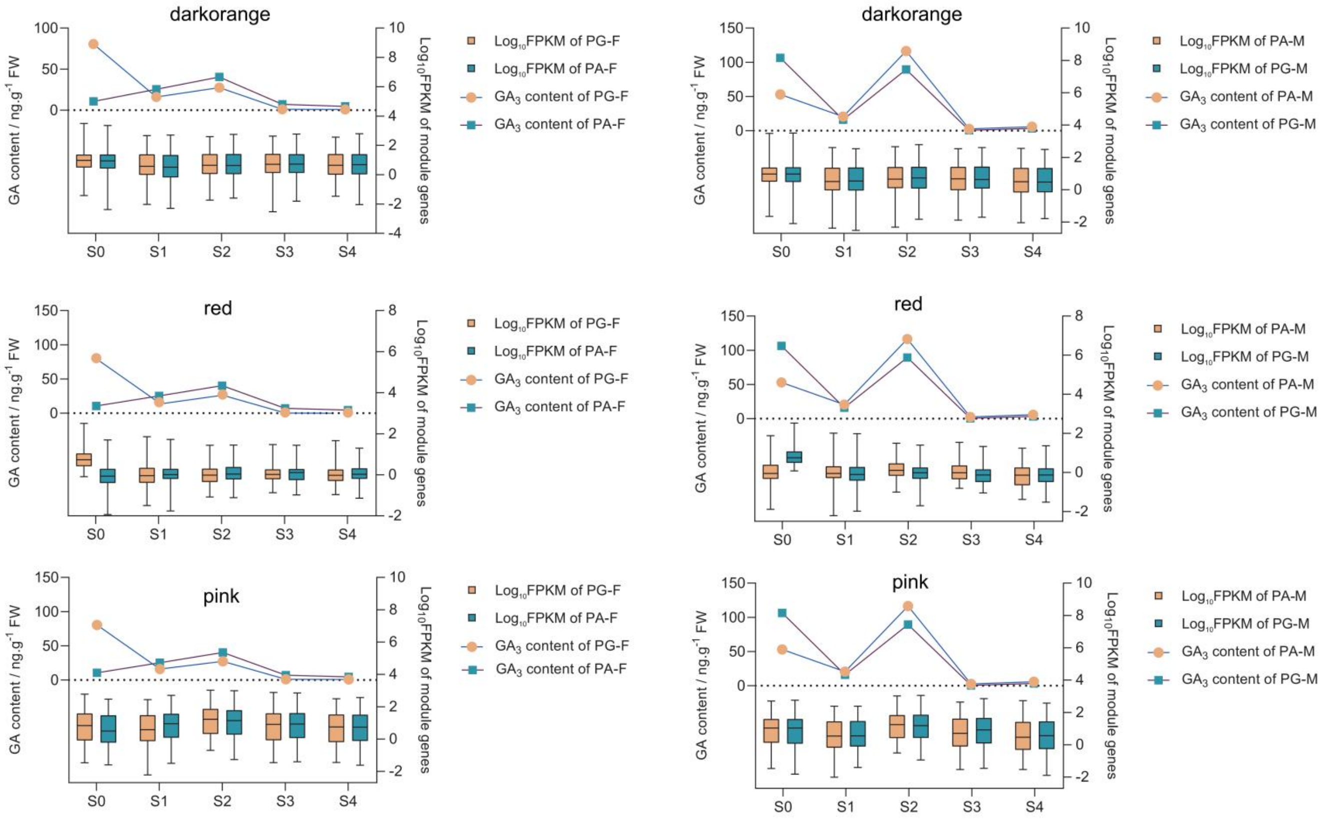


**Supplementary Figure 25. Expression profiles of three modules and $GA_3$ contents during five stages.**

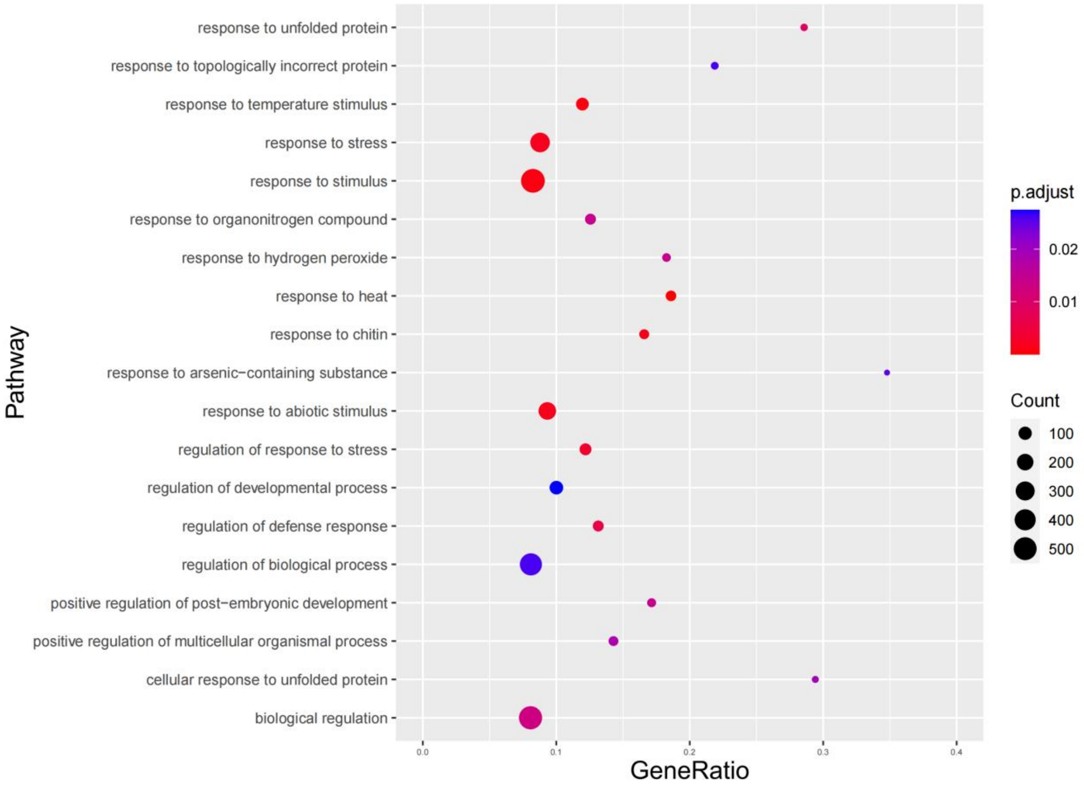


**Supplementary Figure 26. GO enrichment of three modules (darkorange, red, and pink) genes.**

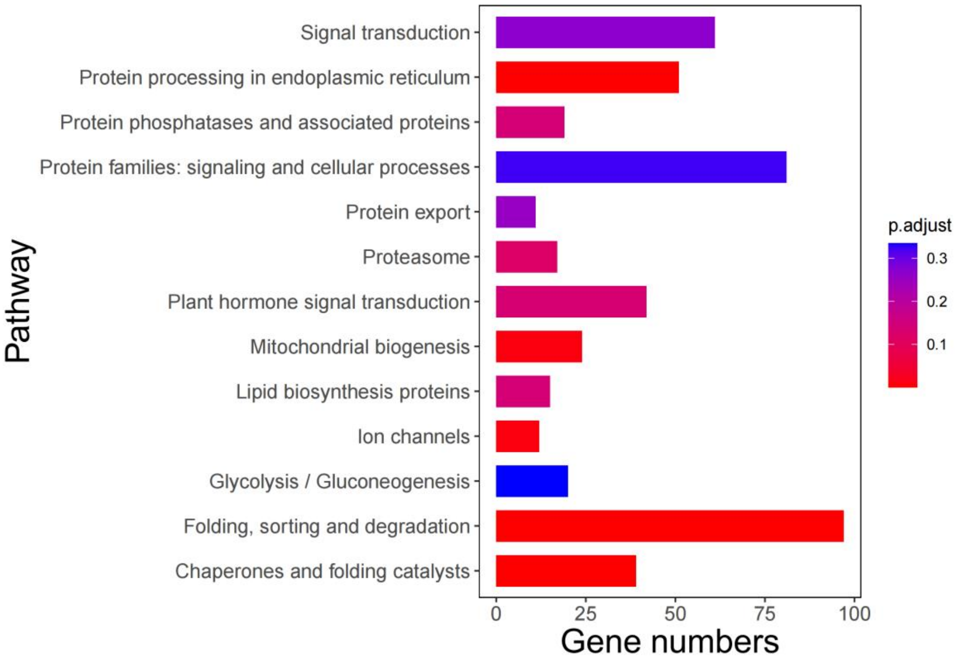


**Supplementary Figure 27. KEGG enrichment of the three modules (darkorange, red, and pink) genes.**

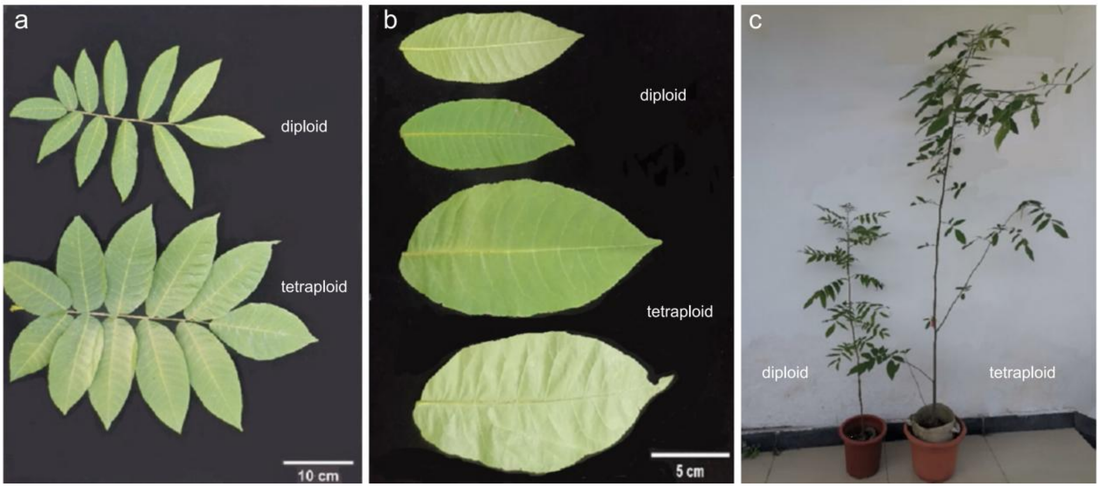


**Supplementary Figure 28. The morphological difference between diploid and tetraploid *C. paliurus* compound leaf (a), singe leaf (b), and seedlings (c).**

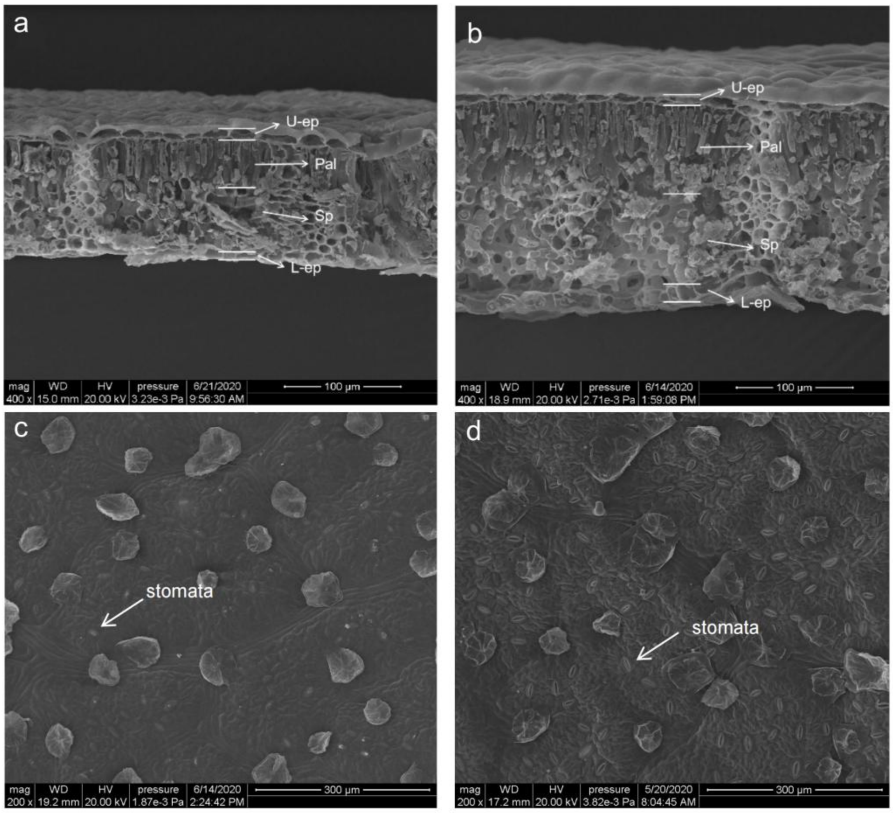


**Supplementary Figure 29. The comparison of leaf thickness and stomatal density between diploid and tetraploid *C. paliurus* based on scanning electron microscopy.** (a) Leaf thickness of diploid sample. (b) Leaf thickness of tetraploid sample. (c) Stomatal density of diploid sample. (d) Stomatal density of tetraploid sample.

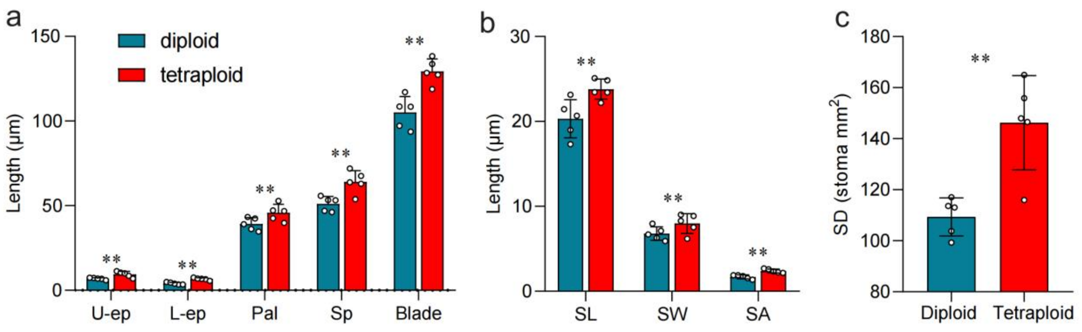


**Supplementary Figure 30. Quantification of anatomical structure values in different ploidy *C. paliurus*.** (a) Thickness of leaf tissues, U-ep (upper epidermal cells), L-ep (lower epidermal cells), Pal (palisade mesophyll), Sp (sponge tissue). (b) Stomatal size of leaf, (SL) stomatal length, (SW) stomatal width, (SA) stomatal aperture. (c) Stomatal density (SD). **, P value < 0.01.

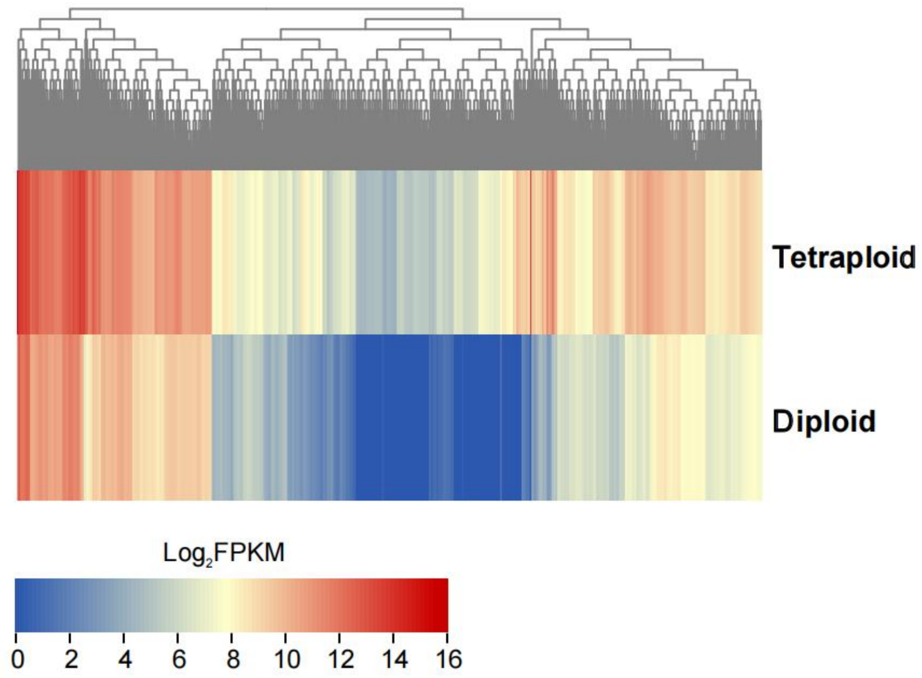


**Supplementary Figure 31. Heatmap showing the expression patterns of 691 dosage-effect genes among tetraploid and diploid samples.**

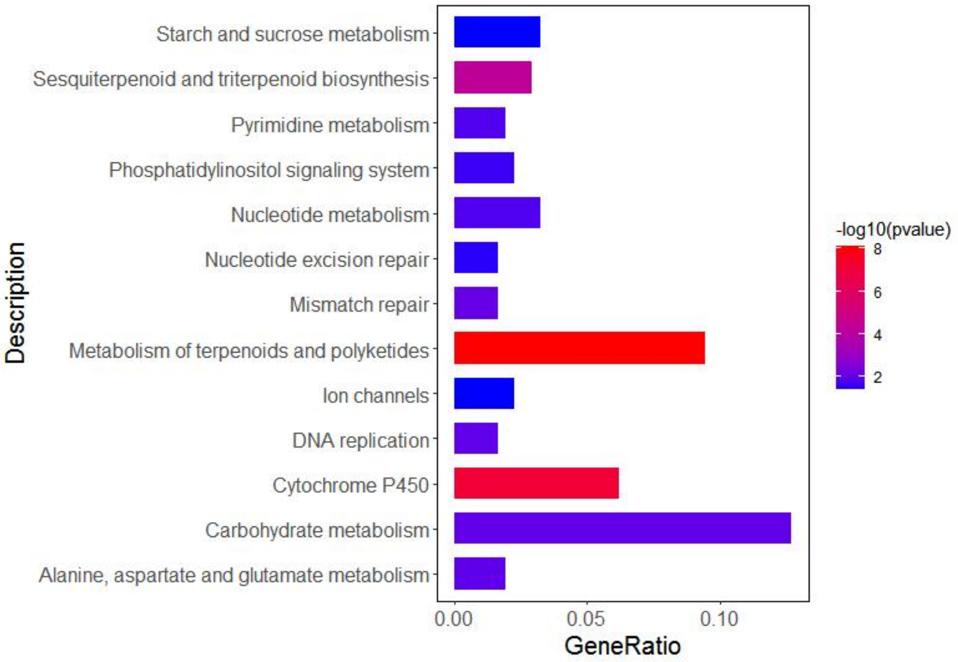


**Supplementary Figure 32. KEGG pathway enrichment analysis of up-regulated genes in tetraploid samples.**

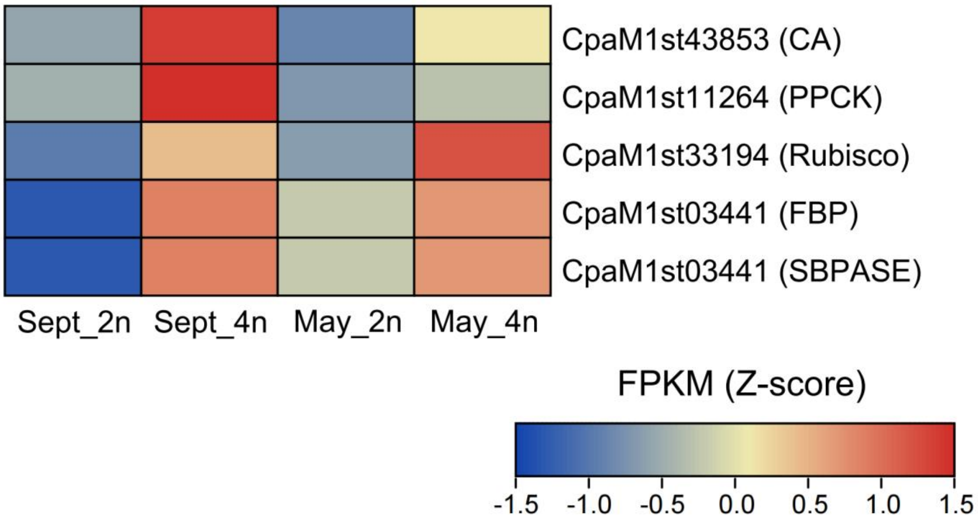


**Supplementary Figure 33. Heatmap showing the expression patterns of photosynthesis related genes. Sept_2n**: diploid samples collected in September; **Sept_4n**: tetraploid samples collected in September; **May_2n**: diploid samples collected in May; **May_4n**: tetraploid samples collected in May**.**

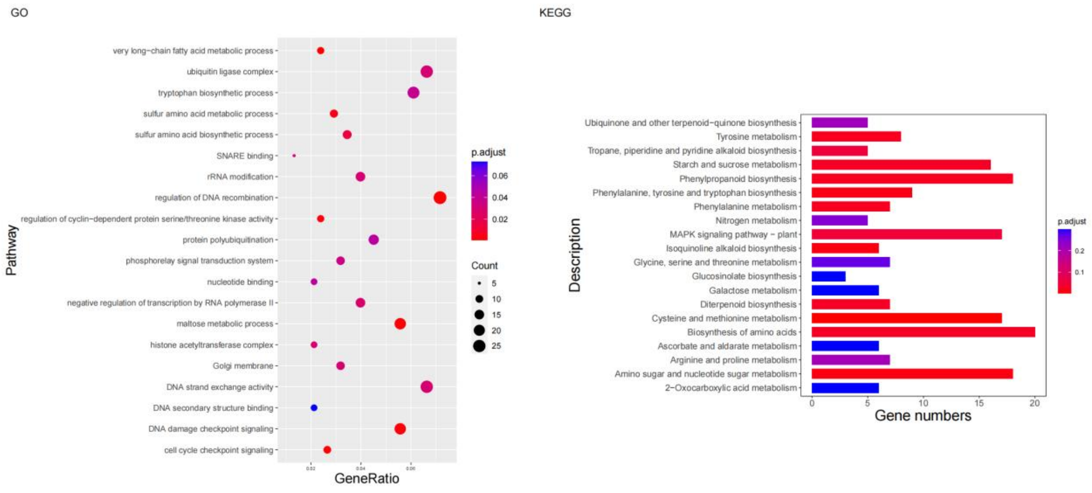


**Supplementary Figure 34. Functional enrichment analysis of dosage compensation effect genes in *C. paliurus.***

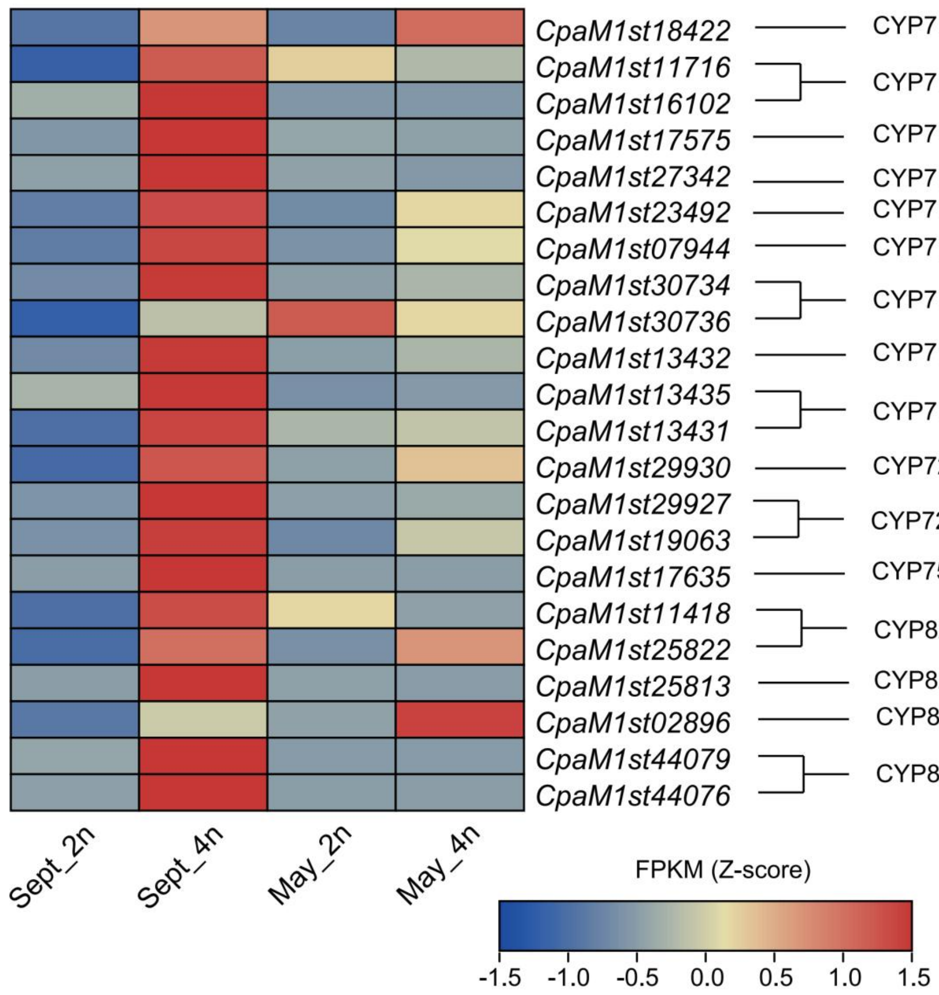


**Supplementary Figure 35. Heatmap showing the expression patterns of P450s family genes of dosage-effect genes across four samples (as in Supplementary Figure 28).**

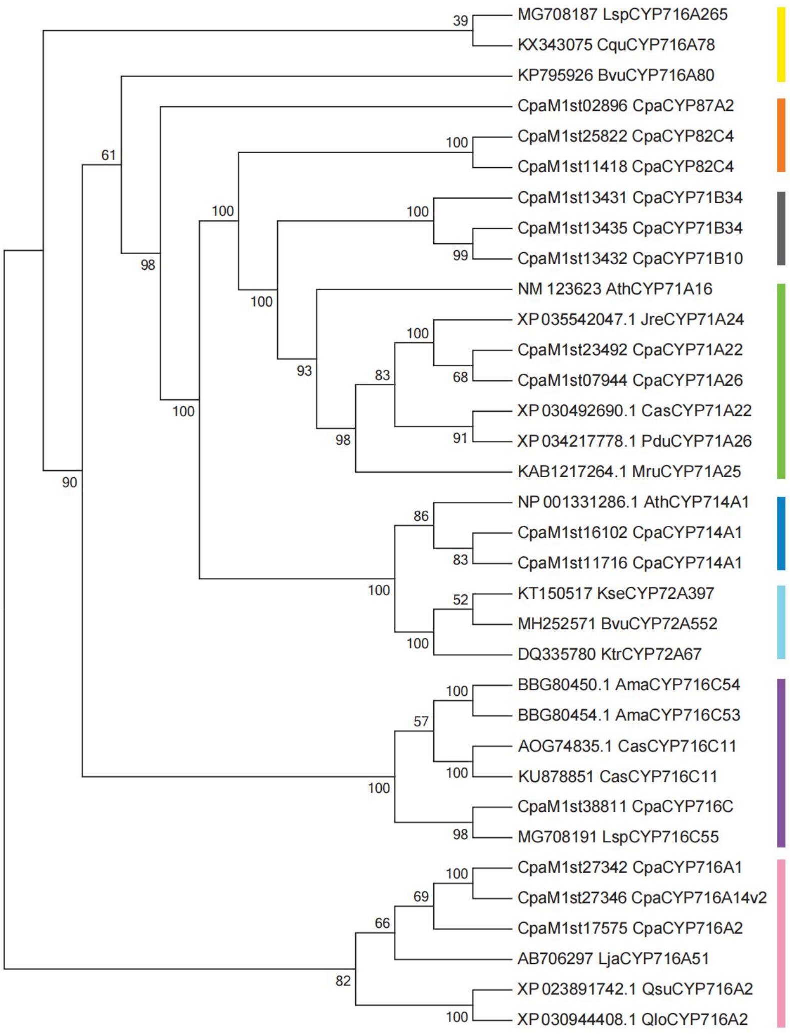


**Supplementary Figure 36. Phylogenetic tree of P450s genes family in multiple species, including *C. paliurus*, *Centella asiatica*, *Avicennia marina*, *Lagerstroemia speciosa*, *Moreua rubra*, *Juglans regia*, *Prunus duscis*, *Arabidopsis thaliana*, *Kalopanax truncatula*, *Barbarea vulgaris*, *Chenopodium quinoa*, *Lotus japonicus*, *Quercus suber*, *Kalopanax septemlobus*, and *Quercus lobata*.**

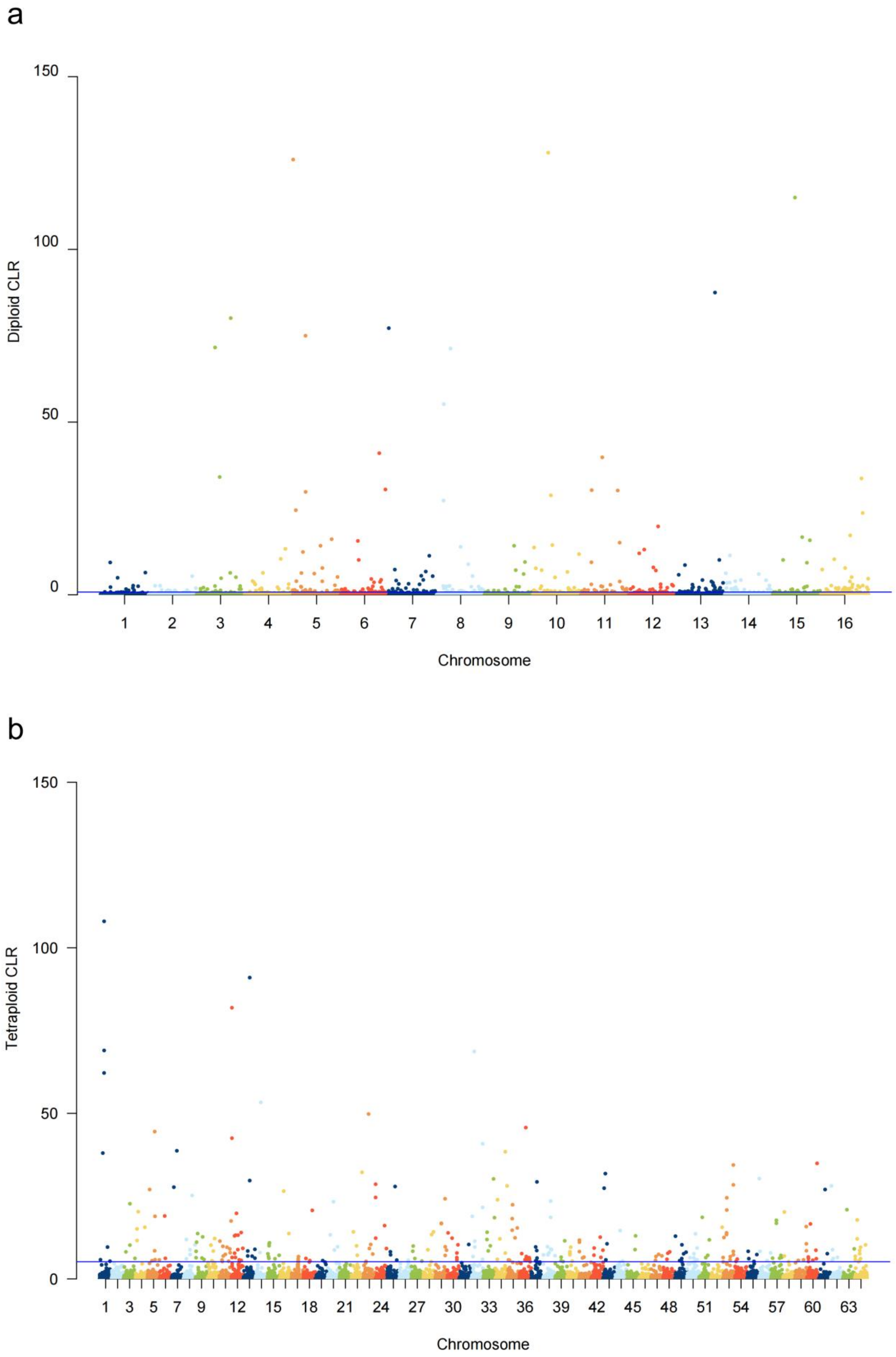


**Supplementary Figure 37. Distribution of selective sweep regions in *C. paliurus* genome.** (a) diploid and (b) auto-tetraploid.

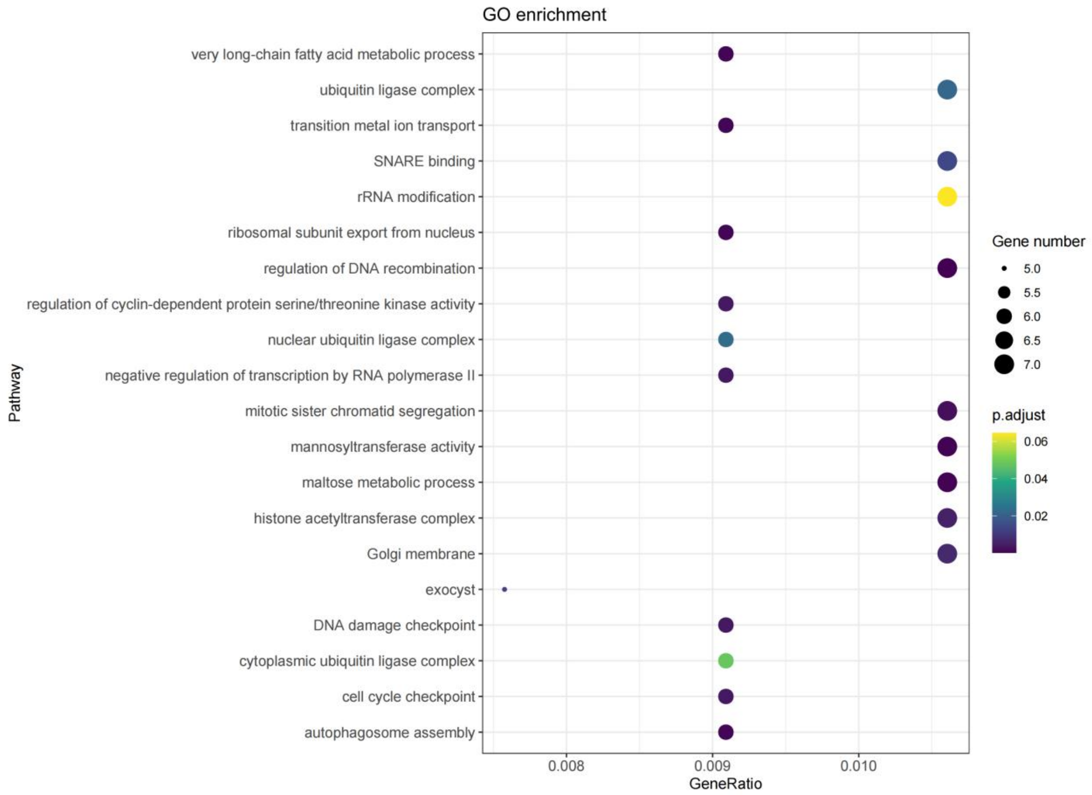


**Supplementary Figure 38. GO enrichment analysis of genes under strong selective sweep in diploid *C. paliurus*.**

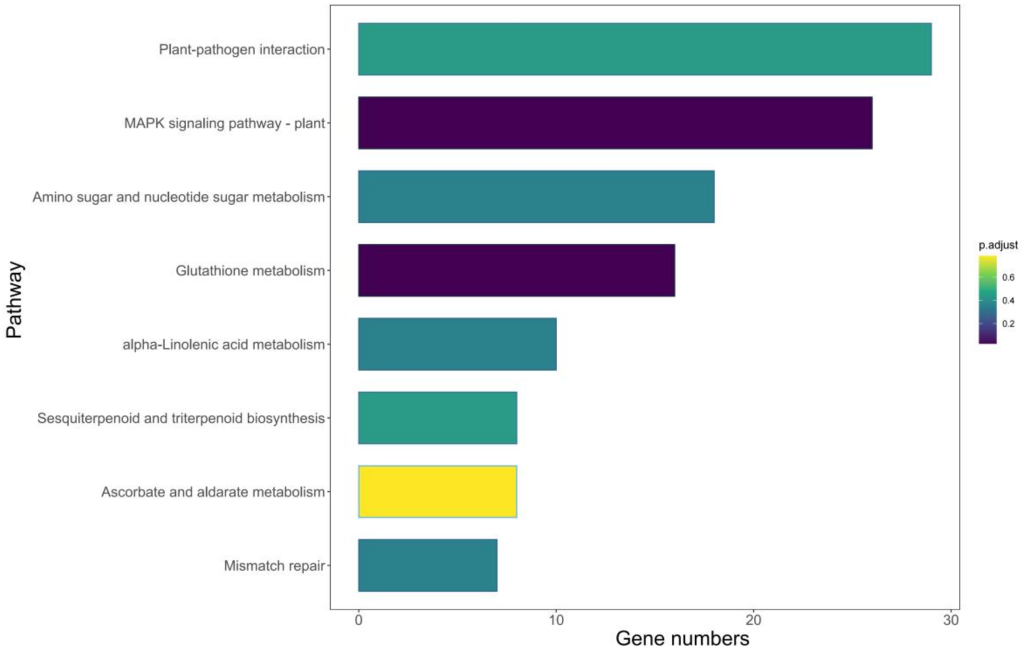


**Supplementary Figure 39. KEGG pathway analysis of genes under strong selective sweep in diploid *C. paliurus*.**

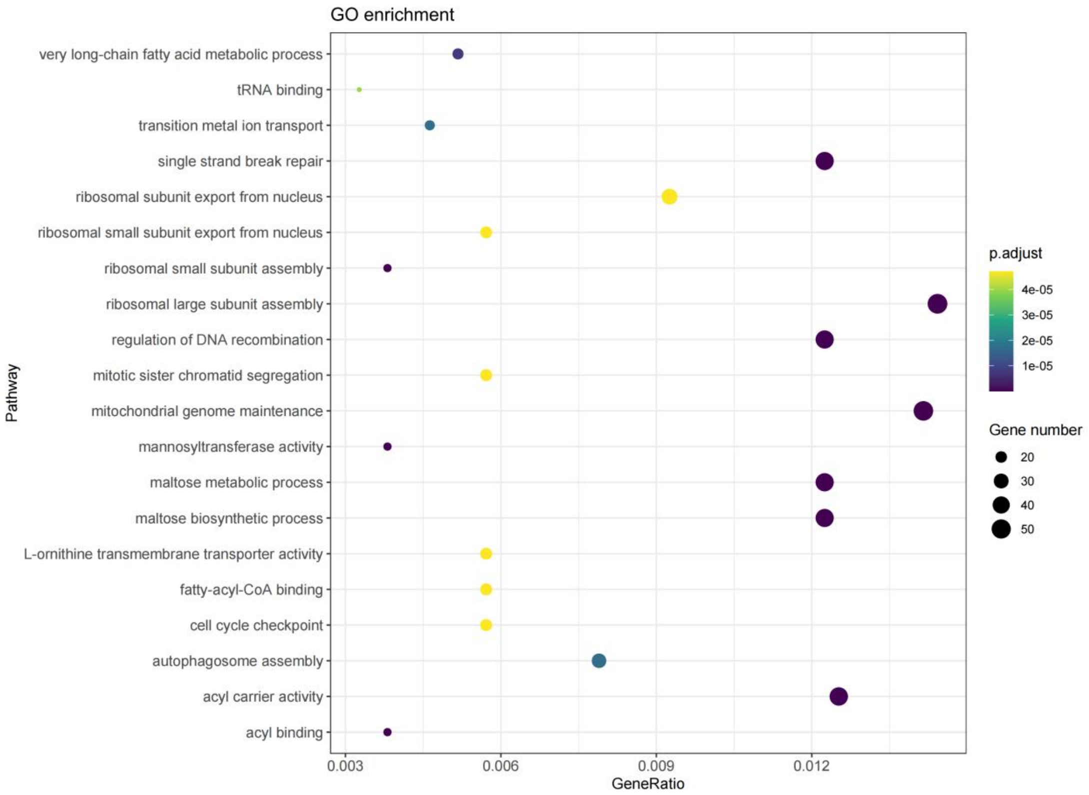


**Supplementary Figure 40. GO enrichment analysis of genes under strong selective sweep in tetraploid *C. paliurus*.**

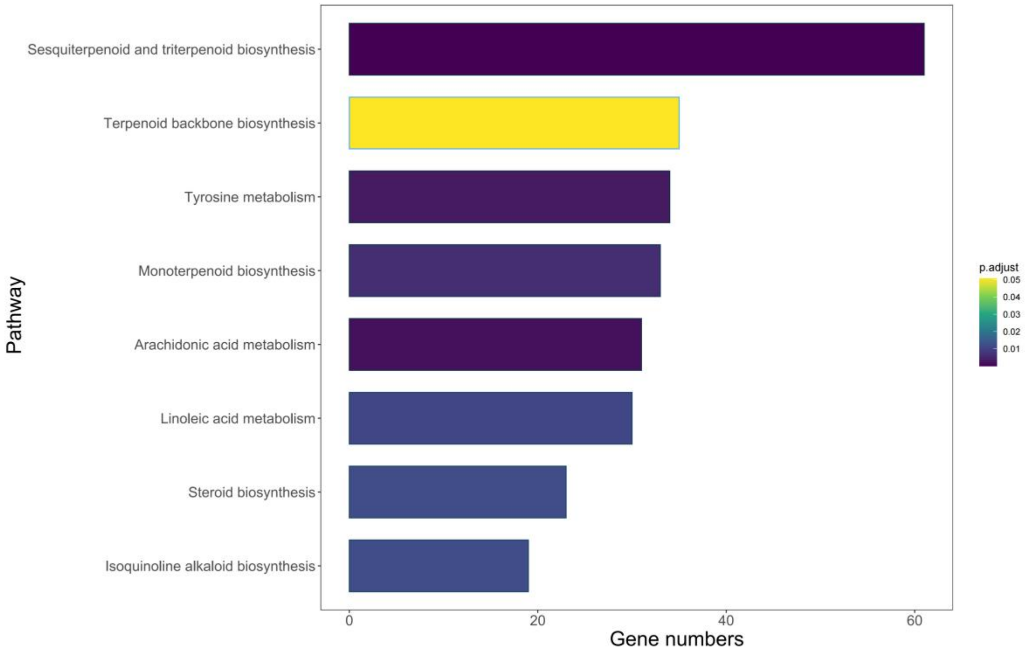


**Supplementary Figure 41. KEGG pathway analysis of genes under strong selective sweep in tetraploid *C. paliurus*.**

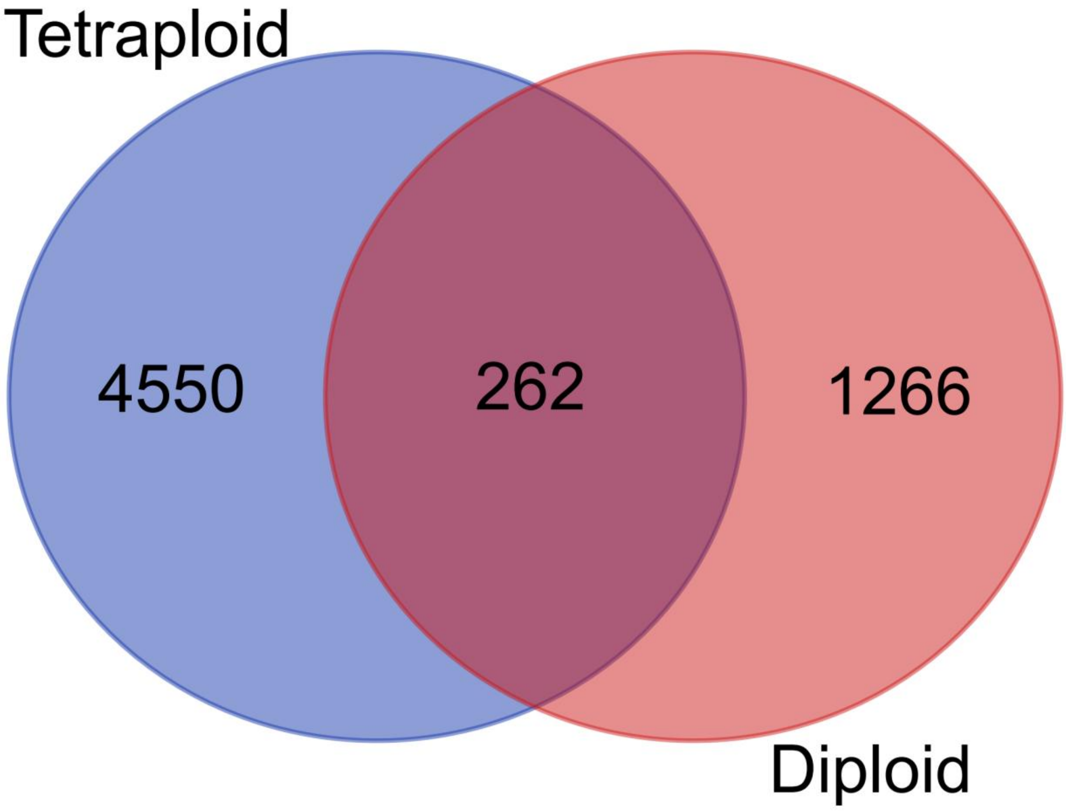


**Supplementary Figure 42. Veen diagrams of selective genes in diploid and tetraploid *C. paliurus.***

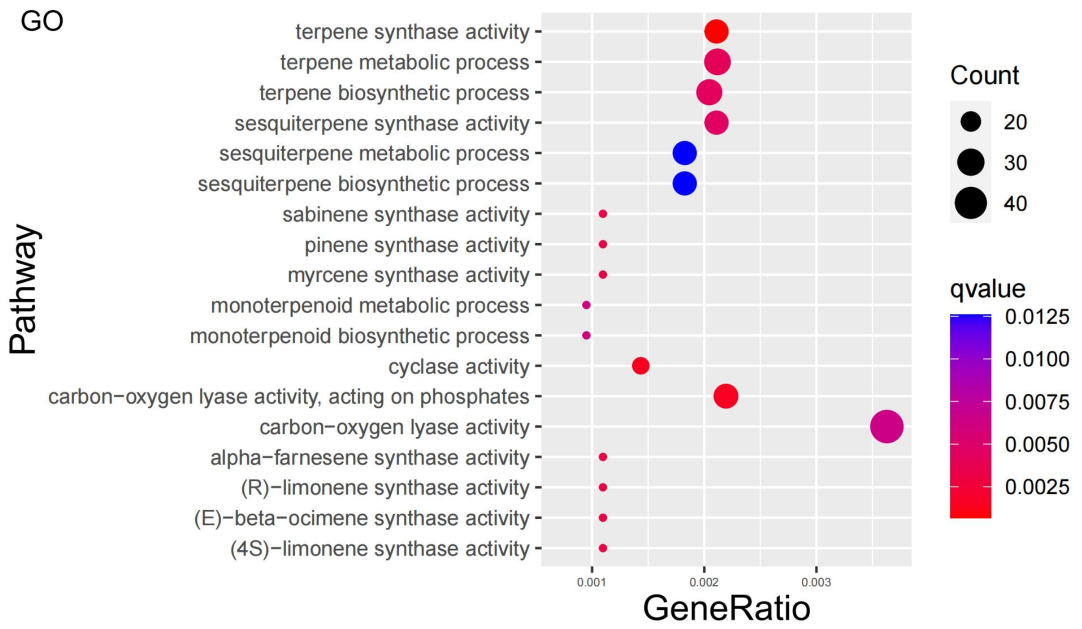


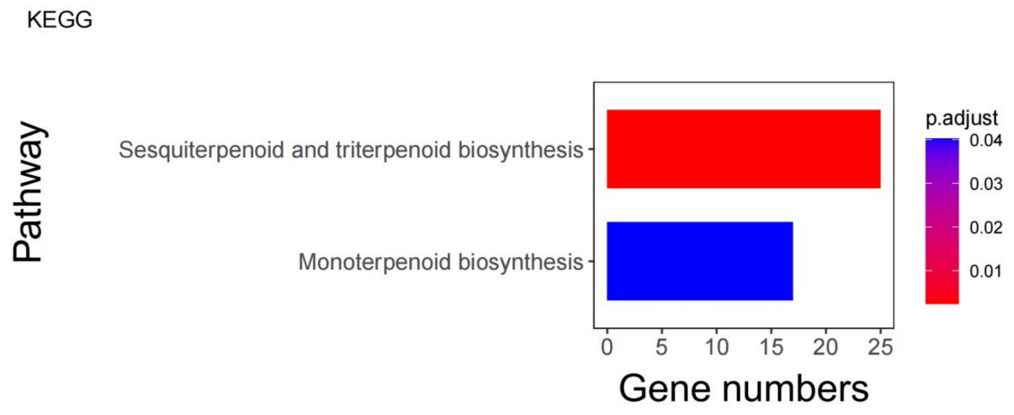


**Supplementary Figure 43. GO and KEGG enrichment of selective genes specific to tetraploid *C. paliurus*.**

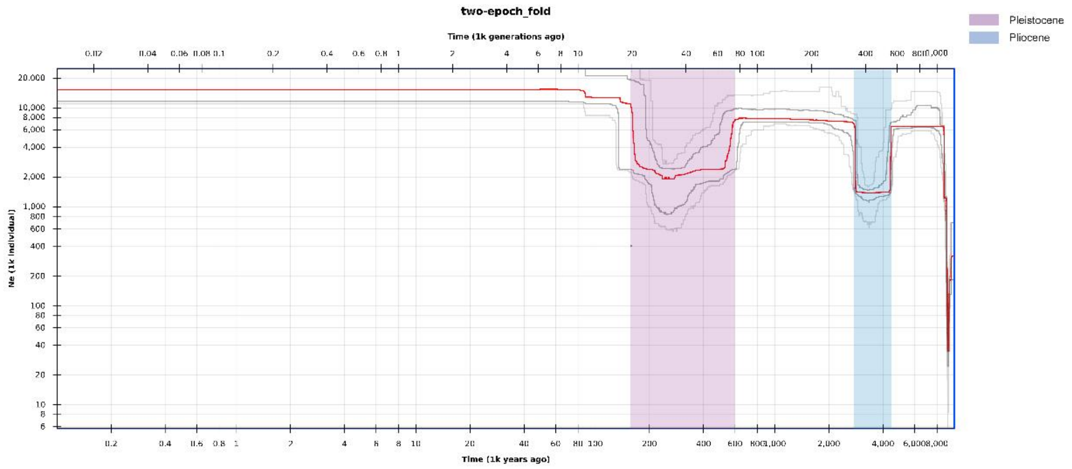


**Supplementary Figure 44.** Historical effective population size for *C. paliurus* beginning from 8 million years ago to present. Stairway plot showing that the *C. paliurus* population has undergone bottlenecks during two known periods of major climate upheaval: the Pleistocene (purple) and the Pliocene (blue).

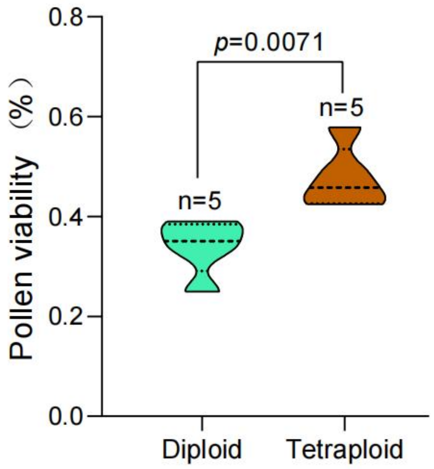


**Supplementary Figure 45. The comparison of pollen viability between diploid and tetraploid *C. paliurus.***
